# Identification of genetic modifiers of Huntington’s disease somatic CAG repeat instability by in vivo CRISPR-Cas9 genome editing

**DOI:** 10.1101/2024.06.08.597823

**Authors:** Ricardo Mouro Pinto, Ryan Murtha, António Azevedo, Cameron Douglas, Marina Kovalenko, Jessica Ulloa, Steven Crescenti, Zoe Burch, Esaria Oliver, Antonia Vitalo, Eduarda Mota-Silva, Marion J. Riggs, Kevin Correia, Emanuela Elezi, Brigitte Demelo, Jeffrey B. Carroll, Tammy Gillis, James F. Gusella, Marcy E. MacDonald, Vanessa C. Wheeler

## Abstract

Huntington’s disease (HD), one of >50 inherited repeat expansion disorders (Depienne and Mandel, 2021), is a dominantly-inherited neurodegenerative disease caused by a CAG expansion in *HTT* (The Huntington’s Disease Collaborative Research Group, 1993). Inherited CAG repeat length is the primary determinant of age of onset, with human genetic studies underscoring that the property driving disease is the CAG length-dependent propensity of the repeat to further expand in brain (Swami *et al*., 2009; GeM-HD, 2015; Hensman Moss *et al*., 2017; Ciosi *et al*., 2019; GeM-HD, 2019; Hong *et al*., 2021). Routes to slowing somatic CAG expansion therefore hold great promise for disease-modifying therapies. Several DNA repair genes, notably in the mismatch repair (MMR) pathway, modify somatic expansion in HD mouse models (Wheeler and Dion, 2021). To identify novel modifiers of somatic expansion, we have used CRISPR-Cas9 editing in HD knock-in mice to enable *in vivo* screening of expansion-modifier candidates at scale. This has included testing of HD onset modifier genes emerging from human genome-wide association studies (GWAS), as well as interactions between modifier genes, thereby providing new insight into pathways underlying CAG expansion and potential therapeutic targets.

## Results and Discussion

*Htt*^Q111^ knock-in mice models have proven to be excellent systems in which to study *HTT* CAG expansion, and we previously identified key modifiers of this process through genetic crosses with mice harboring null mutations (Wheeler *et al*., 2003; Dragileva *et al*., 2009; Kovalenko *et al*., 2012; Mouro Pinto *et al*., 2013; Roy *et al*., 2021), whose relevance has been directly validated through human GWAS (GeM-HD, 2015; Hensman Moss *et al*., 2017; Ciosi *et al*., 2019; GeM-HD, 2019; Hong *et al*., 2021). However, this approach is time-consuming, cost-inefficient and low-throughput. Repeat instability has been studied extensively in model systems such as bacteria, yeast and mammalian cell-based reporter assays (reviewed in Wheeler and Dion, 2021). While much higher throughput, the disease-relevance of these observations (*i.e.* relevance to *in vivo* instability in tissues) can be unclear. To bridge this gap, and with the goal of better understanding the factors and pathways that underlie somatic expansion, we have established a novel *in vivo* CRISPR-Cas9-based system for identifying somatic *HTT* CAG expansion modifier genes by systemic delivery of adeno-associated virus (AAV) expressing single guide RNAs (sgRNAs) targeting genes of interest in Cas9-expressing *Htt*^Q111^ knock-in mice (Wheeler *et al*., 1999; Lee *et al*., 2011; Platt *et al*., 2014) (**Fig.1, Fig.S1, Table S1**). Taking advantage of efficient AAV8 delivery to liver (Gao *et al*., 2002), and the high rate of expansion in liver paralleling that in disease-vulnerable striatum (**Fig.S2**) (Lee *et al*., 2011), we established a relatively high throughput screening platform that provides a sensitive readout of CAG expansion, eliminating the need for genetic crosses with constitutive null alleles and overcoming limitations of embryonic lethality.

**Figure 1.**
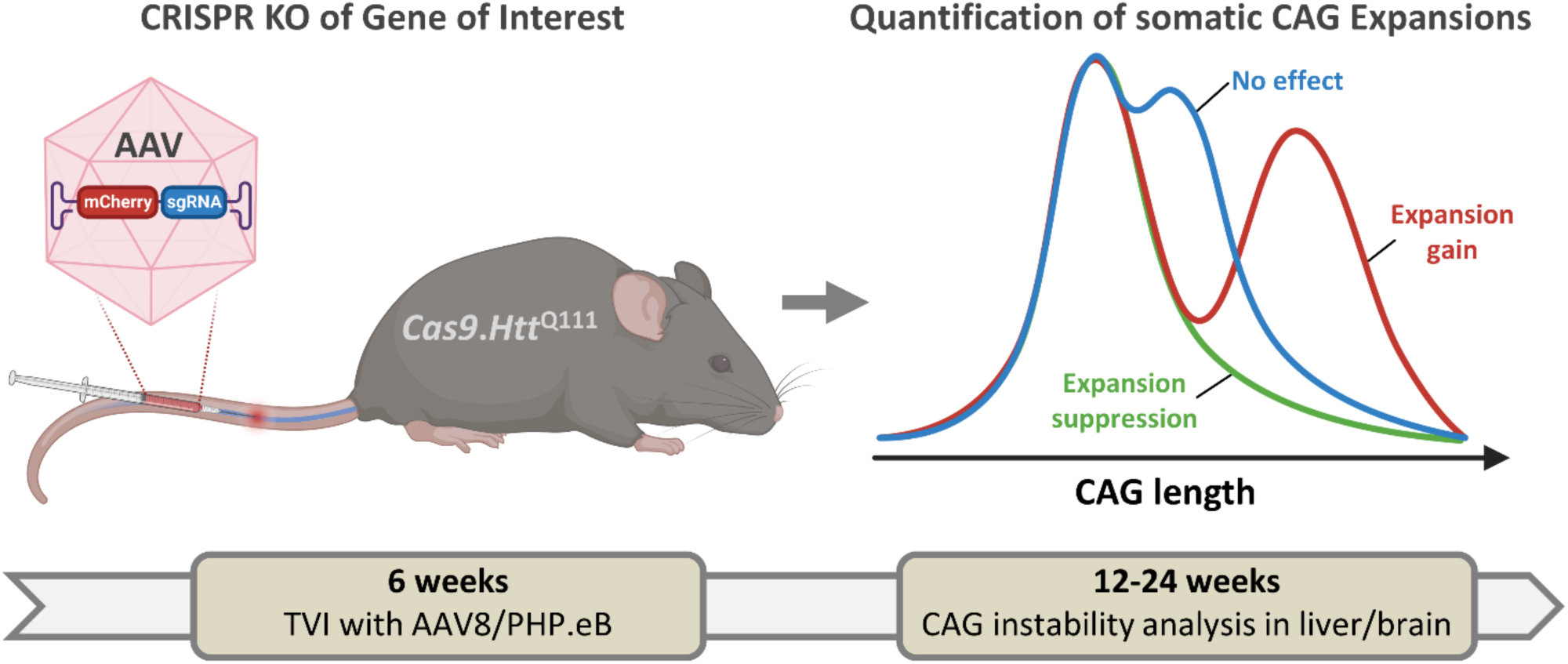
*In vivo* CRISPR editing platform. Adeno-associated virus (AAV8 or PHP.eB) expressing mCherry and a sgRNA targeting a gene of interest is administered to *Htt*^Q111^ mice, which constitutively express Cas9. Tail vein injections (TVI) are performed at 6 weeks of age and mice are aged to 12 or 24 weeks for determination of *Htt* CAG repeat expansion in the liver or striatum, respectively. The blue line depicts a typical profile of CAG repeat lengths in untreated mice that would be observed if there was no effect of the sgRNA, and the green and red lines depict hypothetical CAG length profiles induced by sgRNAs that suppress or promote repeat expansion, respectively. The AAV vector is shown in more detail in **Fig. S1**.

To validate this system we targeted known strong modifiers of *Htt*^Q111^ CAG expansion, namely MMR genes *Msh2*, *Msh3*, *Mlh1* and *Mlh3* (enhancers) and *Fan1* (suppressor) (Wheeler *et al*., 2003; Dragileva *et al*., 2009; Kovalenko *et al*., 2012; Mouro Pinto *et al*., 2013; Loupe *et al*., 2020; Roy *et al*., 2021), achieving efficient viral transduction and editing (**Fig.S3, Tables S1, S2**). Targeting these MMR genes at 6 weeks suppressed expansion, producing a readout at 12 weeks comparable to 6-week mice, whilst targeting *Fan1* enhanced expansions at 12 weeks beyond those in ∼20-week mice (**Fig.2, Table S2**). Control mice did not differ in CAG expansion within the range used (112-119) (**Fig.S4**). Expansion suppression was comparable to that achieved by the respective constitutional homozygous knockout alleles (Wheeler *et al*., 2003; Dragileva *et al*., 2009; Kovalenko *et al*., 2012; Mouro Pinto *et al*., 2013), indicating largely bi-allelic gene inactivation, supported by immunoblot analyses (**Fig.S5**). The impacts of CRISPR targeting were greater than anticipated by simulation based on mixing experiments with liver DNA (**Fig.S2**), likely due to the relatively high editing efficiencies in hepatocytes compared to whole liver (**Table S2, Fig.S6**) (Lee *et al*., 2011) and potential underestimation of inactivating mutations, *e,g*. large deletions and/or the functional impacts of non-frameshift mutations. Overall, we demonstrate highly efficient CRISPR/Cas9-mediated editing in liver allowing detection of both expansion suppressors and enhancers.

**Figure 2.**
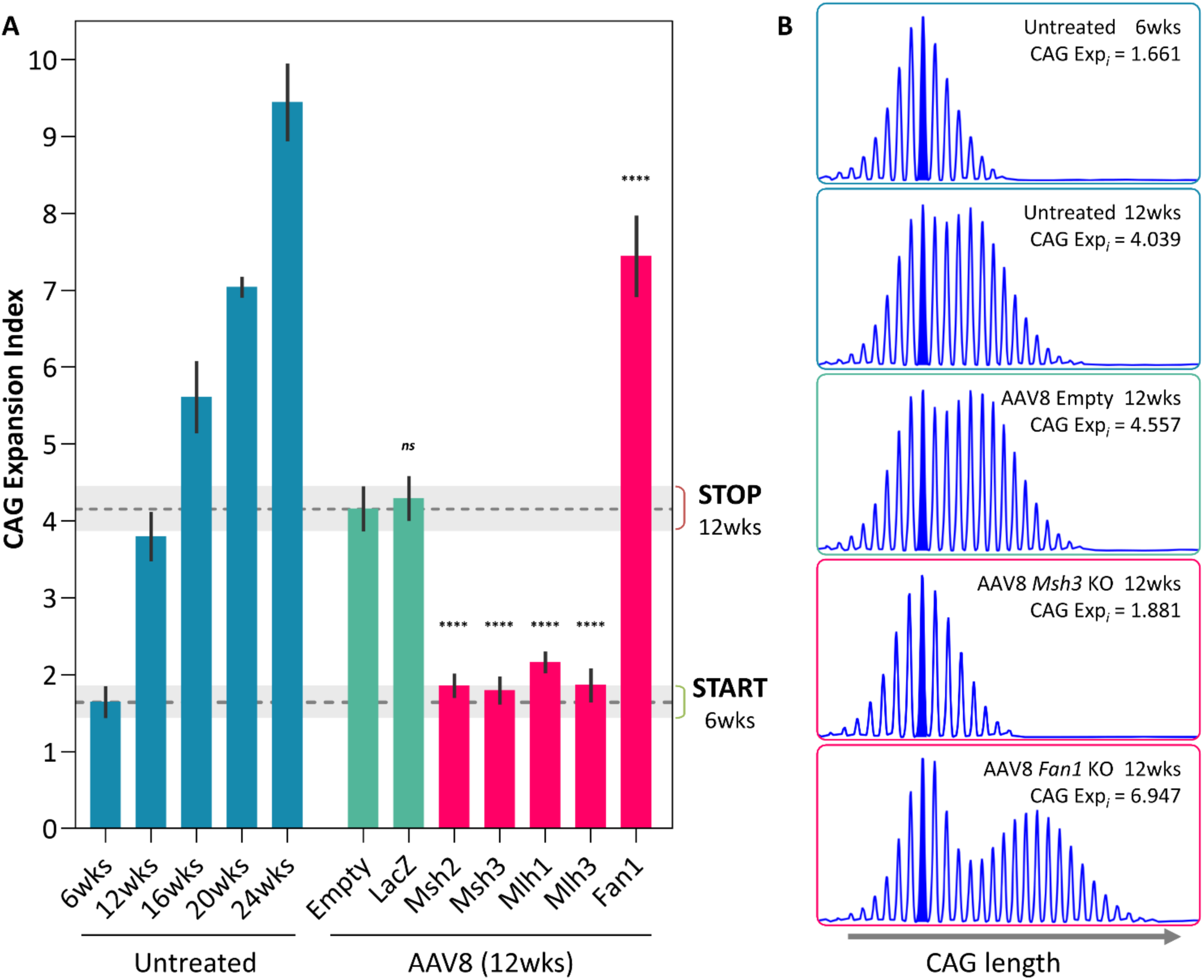
Validation of CRISPR editing platform with known strong modifier genes. **A.** Somatic CAG expansion indices, determined from fragment sizing of *Htt* CAG repeat-containing PCR amplicons of untreated *Htt*^Q111^ Cas9 mice livers from 6 to 24 weeks of age, and in 12-week mice injected at 6 weeks of age with control AAV8s (empty vector or expressing sgRNA targeting LacZ) or AAV8 expressing sgRNAs targeting genes in which null mutations are known to suppress (*Msh2, Msh3, Mlh1, Mlh3*) (Wheeler *et al*., 2003; Dragileva *et al*., 2009; Kovalenko *et al*., 2012; Mouro Pinto *et al*., 2013) or enhance (*Fan1*) (Loupe *et al*., 2020) expansion. Transduction with empty AAV8, or a sgRNA targeting *LacZ,* resulted in a slight background increase in expansion relative to 12-week untreated mice and therefore effects of target sgRNAs are compared to the empty AAV8 vector control. *Msh2, Msh3, Mlh1, Mlh3 and Fan1* relative to empty AAV8: ****p<0.0001 (One-way ANOVA with Dunnett’s multiple comparison correction) (**Table S2**). Bars show mean±s.d. Dotted lines/shaded grey regions show mean/95% confidence interval expansion indices in 6-week untreated mice and in empty AAV8 control-treated mice at 12 weeks of age. **B**. Examples of GeneMapper profiles of *Htt* CAG repeat PCR amplicons.

To gain new insight into the process of repeat expansion we then targeted a large number of genes with previously unknown roles in *Htt*^Q111^ CAG somatic expansion, including candidates from HD onset modifier GWAS (GeM-HD, 2015; GeM-HD, 2019), DNA repair/metabolic genes spanning different pathways, and genes implicated in repeat instability in various model systems (reviewed in Wheeler and Dion, 2021; Zhao *et al*., 2021) (**Fig.3, Tables S3, S4**). Targeting *Pms1, Pold1 and Pold3* elicited expansion suppression comparable to targeting *Msh2*, *Msh3*, *Mlh1* and *Mlh3,* while targeting *Pold2*, *Pold4, Pole, Polb, Pcna, Crebbp, Ercc1, Ercc5, Ercc3, Setd2,* and *Setdb1* resulted in moderate to mild suppression (**Fig.3**). Targeting *Pms2, Msh6, Hmgb1* and *Lig4* increased expansions, though to a lesser extent than *Fan1* (**Fig.3**). Impacts on expansion suppression were broadly similar in hepatocytes and whole liver and not tied to precise editing efficiency (**Fig.3, Fig.S5, Fig.S7, Table S2**). Therefore, lower expansion indices were not due to loss of expansion-harboring hepatocytes, with the exception of *Pcna*, whose targeting resulted in hepatocyte injury/loss and regeneration, precluding the interpretation of *Pcna* as a modifier in this system (**Fig.S8**).

**Figure 3.**
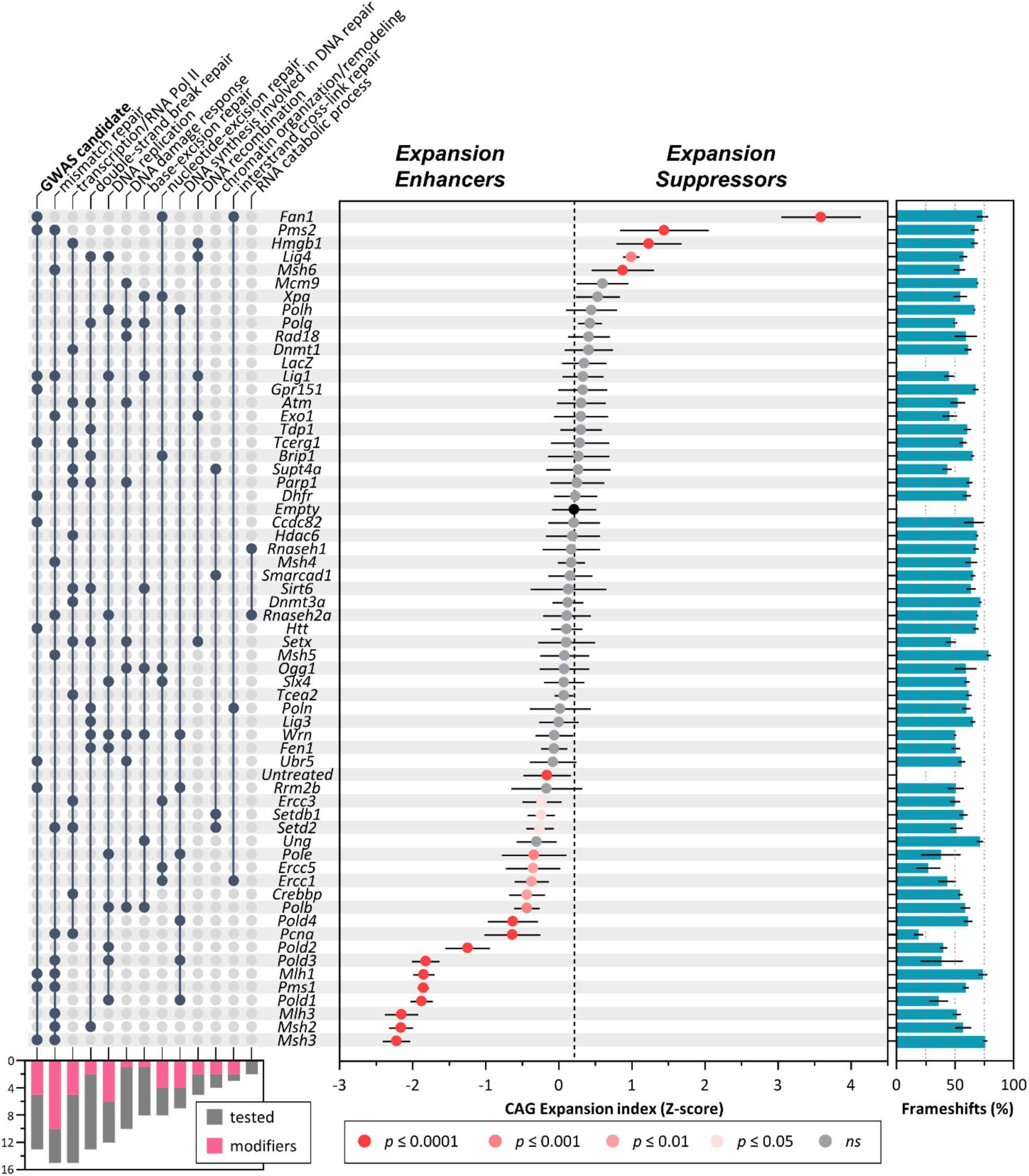
Candidate gene CRISPR knockout screen identifies novel modifiers of CAG expansion. Ranked mean±s.d CAG expansion indices (as Z-score) in livers from 12-week *Htt*^Q111^ Cas9 mice treated at 6 weeks of age with AAV8 expressing sgRNA targeting candidate genes of interest and including empty vector, LacZ and untreated controls. Adjusted p-values are determined in a one-way ANOVA with Dunnett’s multiple comparison test relative to empty vector. The bar graph on the right shows the % frameshift mutation (mean±s.d) for each targeted gene. See **Tables S1 and S2** for details. The UpSet panel on the left indicates candidate genes at genome-wide significant age at onset modifier loci (“GWAS Candidate”) (GeM-HD, 2019), followed by major Gene Ontology (GO) Biological Processes ranked in descending order of number of genes tested. To minimize redundancy, “transcription/regulation by RNA Polymerase II” and “chromatin organization/remodeling” are aggregated terms combined from standard GO terms (see **Table S3**). **Table S3** shows the full set of GO Biological Processes and **Table S4** indicates rationale for gene inclusion. The bottom left bar graph indicates, for each pathway, the number of genes modifying expansion (adjusted p<0.05) relative to the number of genes tested. Note that the reduced expansion index obtained targeting *Pcna* is likely due to hepatocyte loss (**Fig. S8**).

**Figure 4.**
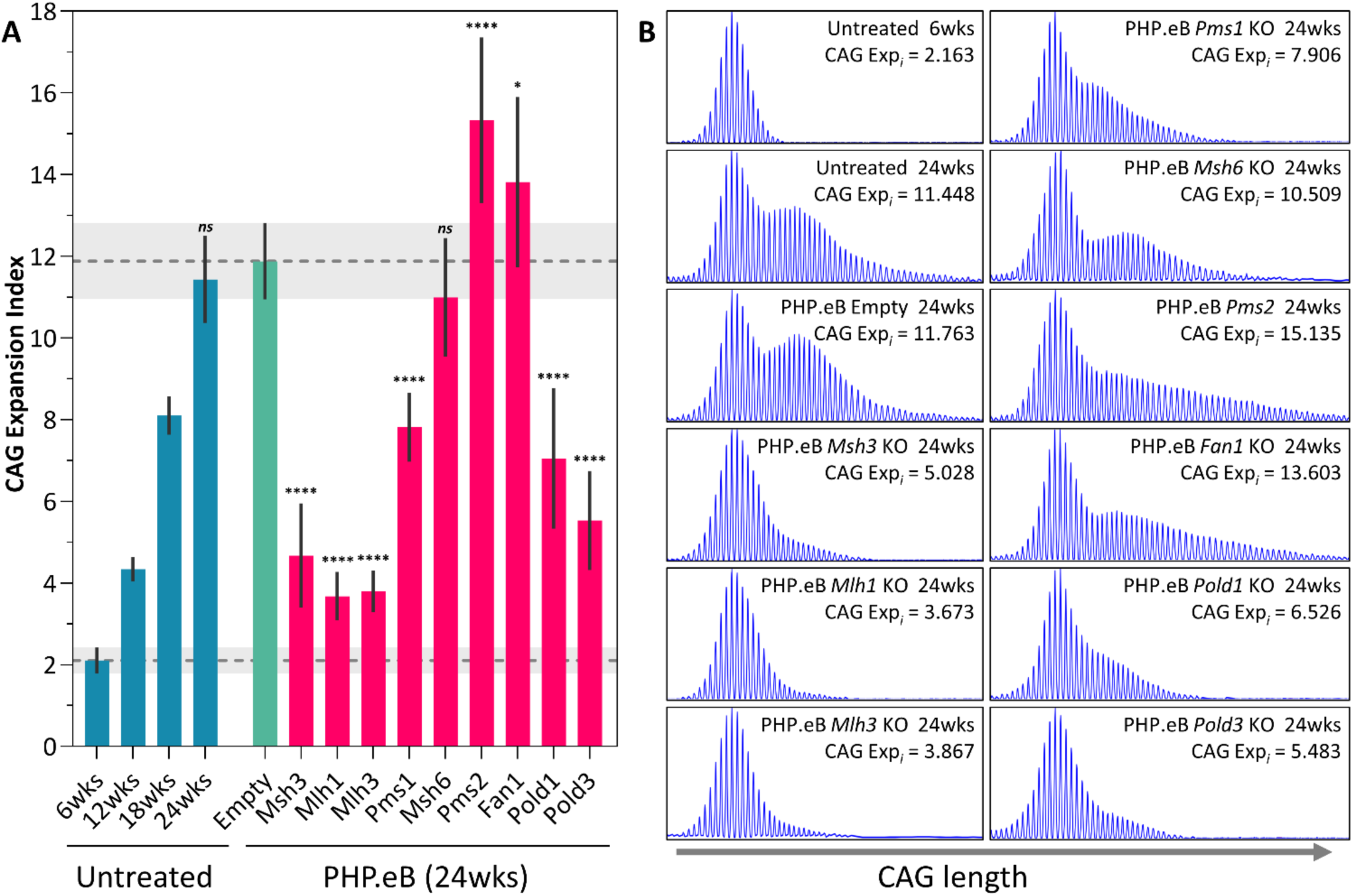
Modification of somatic CAG expansion in the striatum. **A.** Somatic CAG expansion indices, determined from fragment sizing of *Htt* CAG repeat-containing PCR amplicons of untreated *Htt*^Q111^ Cas9 mice striata from 6 to 24 weeks of age, and in 24-week mice injected at 6 weeks of age with control PHP.eB (empty vector) or PHP.eB expressing sgRNAs targeting genes of interest. *\*\*\*\** p<0.0001, * p<0.05 relative to empty vector (One-way ANOVA with Dunnett’s multiple comparison correction) (**Table S5**). Bars show mean±s.d. Dotted lines/shaded grey regions show mean/95% confidence interval expansion indices in 6-week untreated mice and in empty vector control-treated mice at 24 weeks of age. **B**. Examples of GeneMapper profiles of *Htt* CAG repeat PCR amplicons.

Somatic expansion in brain is critical for HD onset (GeM-HD, 2015; GeM-HD, 2019). We therefore tested the impact of targeting a subset of genes on CAG expansion in the striatum, which exhibits high levels of expansion (Lee *et al*., 2010) and is particularly affected in the human disease (Vonsattel *et al*., 1985), by systemic delivery of sgRNAs at 6 weeks using PHP.eB, a recombinant AAV9 derivative capable of crossing the blood brain barrier in C57BL/6 mice (Chan *et al*., 2017), with readout at 24 weeks (**Figs 1, 4, Table S5**). Modifier effects in striatum largely recapitulated those in liver, highlighting *Pms1* and *Pms2*, both GWAS onset modifiers, as well as *Pold1* and *Pold3*, as novel expansion modifiers in brain. The weak *Msh6* modifier effect at 12 weeks in AAV8-treated liver (**Fig.3**, **Table S2)** was not seen in 24-week striatum or liver with PHP.eB treatment **(Fig. 5, S9, Table S5**). Notably, in contrast to the liver (**Fig.3, Fig. S9**, **Table S5**), targeting *Pms2* in striatum promoted expansion to a greater extent than *Fan1*, providing evidence for tissue-dependent effects.

**Figure 5.**
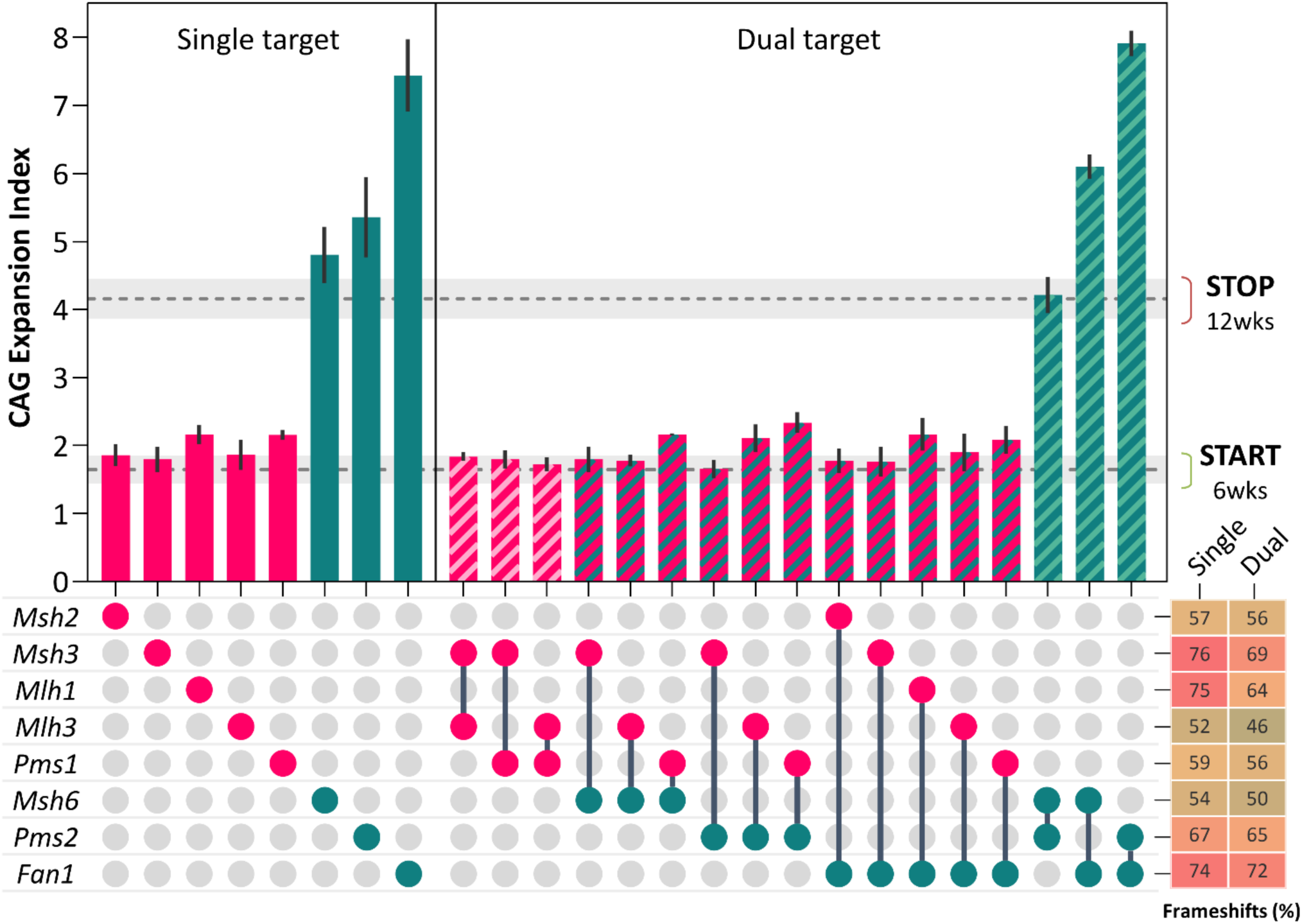
Interactions between modifier genes. Interactions between pairs of expansion modifiers *Msh2, Msh3, Mlh1, Mlh3, Pms1, Msh6, Pms2* and *Fan1* were tested by co-injecting two AAV8s, each targeting a different gene. Expansion enhancers are depicted in red and expansion suppressors are depicted in green. Bars show mean±s.d CAG expansion indices, of *Htt*^Q111^ Cas9 mice livers at 12 weeks of age following injection with either one (single target) or two (dual target) AAV8s. Dotted lines/shaded grey regions show mean/95% confidence interval expansion indices in 6-week untreated mice and in empty vector control-treated mice at 12 weeks of age. The bottom right panel shows the average % frameshift mutation for each guide when injected in a single or dual gene targeting experiment. Refer also to **Tables S1 and S2.** In dual guide targeting with any of the expansion enhancers (*Msh2, Msh3, Mlh1, Mlh3, Pms1)*, expansion indices were not significantly different to those obtained when targeting each enhancer alone (**Table S6**; p>0.999 in all cases). In combinations of expansion enhancers, *Fan1+Pms2* dual guide targeting resulted in an expansion index that was not significantly different to that obtained targeting *Fan1* alone (**Table S6**; p=0.8219). In contrast, targeting *Fan1+Msh6* resulted in a significantly lower expansion index than that targeting *Fan1* alone (**Table S6**; p<0.0001) and a significantly greater expansion index than that targeting *Msh6* alone (**Table S6**; p<0.0001). Targeting *Pms2+Msh6* resulted in a significantly lower expansion index than that targeting *Pms2* alone (**Table S6**; p<0.0001) and a slightly greater expansion index than that targeting *Msh6* alone (**Table S6**; p=0.0632). It appears therefore that the effect of the *Pms2* knockout is redundant to that of the *Fan1* knockout and the effects of both *Fan1* and *Pms2* knockouts are at least partially dependent on the presence of *Msh6*.

We then extended this system to investigate genetic interactions in 12-week liver, since simultaneously targeting two genes with two AAV8s did not appreciably alter editing efficiency compared to that achieved by single gene targeting (**Fig.5, Table S6**). We previously reported that *Fan1*’s expansion suppression was dependent on *Mlh1* (Loupe *et al*., 2020), an effect recapitulated here by CRISPR targeting *Fan1*+*Mlh1* (**Fig.5, Table S6**). Extending this paradigm to multiple suppressor/enhancer combinations, the effects of suppressors *Fan1*, *Pms2* or *Msh6* were fully dependent on enhancers *Msh2, Msh3, Mlh1, Pms1* and *Mlh3* (**Fig.5, Table S6**). In combinations of two suppressors, targeting *Fan1+Pms2* resulted in a phenotype indistinguishable from the *Fan1* knockout, while targeting *Fan1+Msh6* or *Pms2+Msh6* reduced the impacts of the respective *Fan1* or *Pms2* knockouts. (**Fig.5, Table S6**), indicating complex interactions among the suppressors. We also probed potential functional redundancies that could underlie the lack of effect of single targeting, performing dual targeting for *Lig1+Lig3*, *Dnmt1+Dnmt3a* and *RNaseh1+RNaseh2a*, though none impacted CAG expansion (**Fig.S10**).

Our *in vivo Htt*^Q111^ CRISPR-Cas9 editing platform thus provides a highly efficient means of screening and identifying novel expansion modifying genes and their interactions, though it is worth noting that the absence of an effect in this system does not necessarily preclude a role for a particular gene in CAG expansion due to potential insensitivity to detect modifiers of weak effect (**Fig.S2**), compensatory responses, functional redundancies or cell type-dependent modifier effects. Importantly, we have identified a number of novel modifiers, including modifiers of CAG expansion in the striatum. We highlight several observations: Firstly, targeting the orthologues of HD onset modifier candidates (GeM-HD, 2015; GeM-HD, 2019) (**Fig.3, Tables S3, S4**) provides the first *in vivo* evidence for *PMS1* and *PMS2* modifying HD via an impact on *HTT* CAG expansion. In contrast, *TCERG1* and *CCDC82* may more likely modify HD via other mechanisms (GeM-HD, 2019; Hong *et al*., 2021). Constitutional *Rrm2b* knockout was previously shown to slightly suppress expansion in liver and striatum (Loupe *et al*., 2020), consistent with the weak (non-significant) effect seen here. *LIG1* seems likely to alter HD onset via an effect on repeat instability, therefore the lack of effect targeting *Lig1* here may suggest functional redundancy, consistent with a human *LIG1* mutation having a greater impact than a knockout (Bentley *et al*., 2002), highlighting the need to dissect the functional variants from human GWAS. Of all the other genes tested, only *POLD1* has subsequently emerged as a genome-wide significant disease modifier in the most recent HD GWAS (GeM-HD Consortium, *in submission*).

Secondly, our data do not support an equally important role for all DNA repair pathways in somatic *HTT* CAG expansion (Fig.3); rather, members of the MMR pathway, together with FAN1, are clearly highlighted as the key players. The involvement of LIG4, as observed in the liver of a Fragile X-related disorders mouse model (CGG repeats) (Gazy *et al*., 2019), suggests a suppressive role for a double-strand break repair process that warrants further study. We also expose roles for transcription and chromatin-related processes, supporting previous observations in different systems (**Table S4**) (reviewed in Wheeler and Dion, 2021), and consistent with the idea that transcription through the repeat and/or an open chromatin structure are important for repeat instability. ERCC1, ERCC3 (XPB) and ERCC5 (XPG) are involved in transcription-coupled repair (Iyama and Wilson, 2013), CBP *(Crebbp)*, SETD2 and SETDB1 modify histones (Vempati *et al*., 2010; Wagner and Carpenter, 2012; Markouli, Strepkos and Piperi, 2021) and HMGB1, a non-histone chromatin-associated protein, is involved in several DNA repair pathways including MMR (Yuan *et al*., 2004; Zhang *et al*., 2005; Genschel and Modrich, 2009; Tang *et al*., 2023) and binds slipped repeat structures (Slean *et al*., 2013). HMGB1 promoted CAG expansion in a cell-free system (Liu *et al*., 2009), in contrast to its suppressor role identified here, implying a different mechanism *in vivo*. SETD2 recruits MutSα (MSH2-MSH6) to chromatin (Huang and Li, 2020); however, the opposing directions of effects targeting *Setd2* and *Msh6* do not obviously support such a role *in vivo*. CBP, SETD2 and SETDB1 may alternatively act via post-translational modification of DNA repair proteins themselves (Kovalenko *et al*., 2020; Williams *et al*., 2020).

Thirdly, our data indicate non-redundant functions of MutLβ (MLH1-PMS1) and MutLγ (MLH1-MLH3) that may be distinguished by MLH3’s endonuclease activity, critical for expansion (Kadyrova *et al*., 2020; Roy *et al*., 2021), and a possible chromatin structural role of PMS1 that lacks endonuclease activity (Duroc *et al*., 2017). Interestingly in this regard, PMS1 possesses a high-mobility group (HMG) box (Stros, Launholt and Grasser, 2007). In contrast, *Pms2* suppresses expansion, indicating a distinct role for MutLα (MLH1-PMS2). Notably, *Pms1* enhanced *HTT* CAG and Fragile X GGC expansion in cell-based models (Miller *et al*., 2020; Ferguson *et al*., 2023; McLean *et al*., 2024), while *Pms2* variably enhanced or suppressed expansion in cell or mouse models of Fragile X, myotonic dystrophy and Friedreich ataxia (Gomes-Pereira *et al*., 2004; Bourn *et al*., 2012; Ezzatizadeh *et al*., 2012; Miller *et al*., 2020; Ferguson *et al*., 2023). Therefore, the precise roles of MutL proteins appear dependent on the disease-associated repeat context and/or cell-type, with important therapeutic implications.

Finally, the role of DNA polymerase delta (*Pold*) in CAG expansion supports a novel role for Exonuclease 1-independent MMR, dependent on POLδ’s strand displacement activity (Amin *et al*., 2001; Kadyrov *et al*., 2009; Desai and Gerson, 2014; Goellner, Putnam and Kolodner, 2015). Consistent with this, *Pold1* and *Pold3*, encoding POLδ’s catalytic and strand-displacement activities, were stronger modifiers than *Pold2* and *Pold4*, the latter promoting proofreading whilst limiting strand-displacement (Stith *et al*., 2008; Meng *et al*., 2010; Prindle and Loeb, 2012; Lin *et al*., 2013). We also identify minor roles for POLβ, previously implicated in repeat instability and proposed to interact with MSH3 in base-excision repair (Lokanga *et al*., 2015; Guo *et al*., 2016; Lai *et al*., 2016), and POLε, which can act in MMR *in vitro* (Bowen and Kolodner, 2017), but our data do not obviously support a noncanonical form of MMR dependent on POL11 (*Polh*) (Peña-Diaz *et al*., 2012).

We propose a model (Fig.6) that incorporates the strongest modifiers, integrating and extending existing observations. The model is centered on the idea that a predominant MutSβ (MSH2-MSH3)- and MutLγ (MLH1-MLH3)-dependent mechanism underlies the somatic expansion bias of the repeat mutation as consequences of: a) the preferential binding of MutSβ to CAG/CTG loop-outs (Owen *et al*., 2005; Panigrahi *et al*., 2010; Pluciennik *et al*., 2013), b) the preferential recruitment of MutLγ to MutSβ-bound DNA (Rogacheva *et al*., 2014; Cannavo *et al*., 2020; Kadyrova *et al*., 2020), and c) MutLγ endonucleolytic activity that is critical for expansion (Roy *et al*., 2021) and which is biased towards the strand opposite the loop-out, thus permitting the incorporation of the additional DNA (Kadyrova *et al*., 2020; Roy *et al*., 2021). While PMS1 and POLδ promote this pathway, other factors, *e.g.* PMS2, MSH6, FAN1, HMGB1 suppress it, either directly or indirectly. *e.g*. processing of loop-outs by MutLα, whose exonuclease activity, directed to either strand does not favor expansions (Gomes-Pereira *et al*., 2004; Pluciennik *et al*., 2013; Kadyrova *et al*., 2020), or actions of FAN1 binding to loop-outs or to MLH1 (Goold *et al*., 2021; Phadte *et al*., 2023). This model predicts that the likelihood of an expansion will depend on the chance of a CAG/CTG loop-out being processed via this expansion-promoting pathway, which will in turn depend on the relative levels of the various protein components and their complexes in cells. This may in part underlie cell type differences in repeat expansion (Gonitel *et al*., 2008; Lee *et al*., 2011; Mätlik *et al*., 2024; Pressl *et al*., 2024) as well as the sensitivity with which human HD modifier variants capture different HD phenotypes (GeM-HD Consortium, *in submission*).

**Figure 6.**
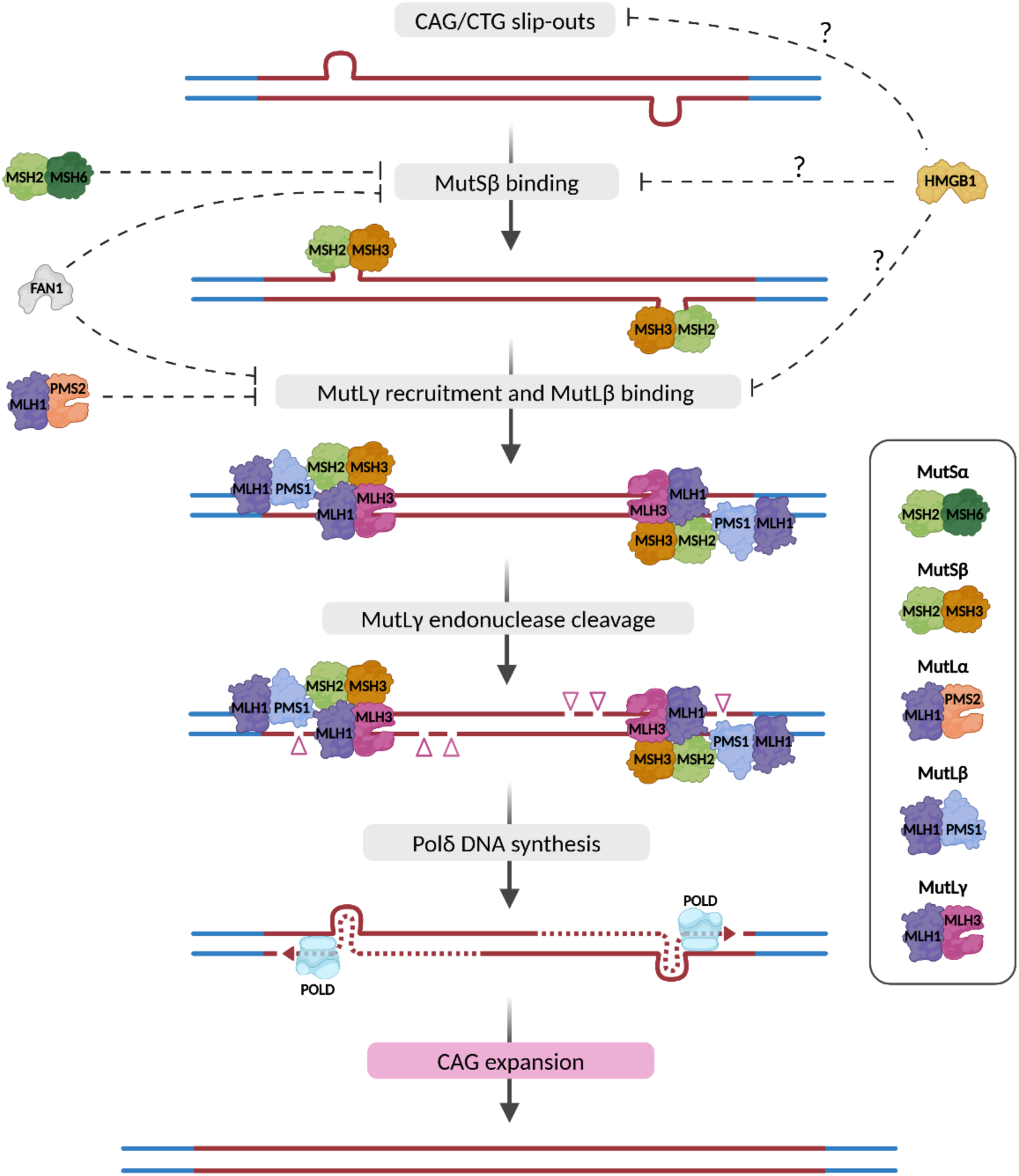
Model integrating major modifiers of repeat expansion. The model depicts the principal pathway driving repeat expansion, with main expansion enhancers shown. CAG/CTG loop-outs are generated *e.g*. in the process of transcription, chromatin remodeling or breathing and are preferentially recognized by MutSβ (MSH2-MSH3) which preferentially recruits MutLγ (MLH1-MLH3) (Owen *et al*., 2005; Panigrahi *et al*., 2010; Pluciennik *et al*., 2013; Rogacheva *et al*., 2014; Cannavo *et al*., 2020; Kadyrova *et al*., 2020). The role of PMS1 is unclear but it may play a facilitating role as part of the MutLβ (MLH1-PMS1) dimer at the level of chromatin structure, potentially stabilizing MutSβ or the subsequent MutSβ-MutLγ complex. MutLγ cleaves the DNA strand opposite the loop-out, driving an expansion bias (Kadyrova *et al*., 2020; Roy *et al*., 2021). Strand displacement synthesis by POL8 results in gap-filling that includes the looped-out DNA. Exonuclease-dependent strand excision cannot be ruled out, and minor roles for DNA POLε and POLβ are also implicated (**Fig.3**). DNA ligase seals the resulting nick, and if the loop-outs on each strand are both processed independently (Gomes-Pereira *et al*., 2004), the length of the loop-out will be incorporated as an expansion. Expansion suppressors such as FAN1 PMS2, MSH6 and HMGB1 may interfere at various steps in this pathway. MutSα (MSH2-PMS6) may decrease MutSβ complex formation and/or and binding to loop-outs. MutLα (MLH1-PMS2) to loop-outs bound by MutSβ results in endonucleolytic cleavage on either strand, with no expansion bias (Pluciennik *et al*., 2013; Kadyrova *et al*., 2020). FAN1 may inhibit MutSβ binding by its direct binding to loop-outs (Phadte *et al*., 2023) or may inhibit MutLγ recruitment by sequestering MLH1 (Goold *et al*., 2021). The suppressive function of HMGB1 is unclear but could potentially act at several steps. Other factors (**Fig.3**) may also play roles enhancing or suppressing this pathway. Thus, the likelihood of an expansion event will depend on the steady state levels of the different expansion enhancer and suppressor proteins/complexes, which are likely to differ by cell type. Genetic interactions (**Fig.5**) indicate that PMS2 is redundant to FAN1 in its expansion-suppressing function in liver. The reduced impacts of the *Fan1* and *Pms2* knockouts in the absence of *Msh6* (**Fig.5**) are not readily explained, and we speculate that this may implicate a direct role for MSH6 and the existence of a MSH6-dependent factor that enhances FAN1 and PMS2’s expansion-suppressing effect. The model is based on an assumption that MSH and MLH subunits function as part of canonical heterodimeric complexes; however, there is evidence for functions of PMS2 and MLH3 that are independent of MLH1 (summarized in (Pannafino and Alani, 2021)), and similarly, noncanonical roles for MMR subunits may also be plausible in mechanisms underlying CAG instability. Dark red lines are repeat sequence; blue lines are flanking non-repeat sequence. Triangles mark endonuclease cleavage.

In summary, we have developed a novel CRISPR-Cas9-based screening approach, providing a paradigm for *in vivo* genetic modifier studies of other repeat expansion disorders that represent a growing class of currently intractable human diseases. This has allowed us for the first time to systematically test a large number of genes, under standardized conditions *in vivo*, for their role in CAG expansion, and represents the first *in vivo* study to analyze modifiers of repeat instability in any disease at this scale. Our results emphasize the importance of studying repeat instability in mice tissues, where modifiers may act differently to other model systems. We have demonstrated high efficiency of genome editing following systemic PHP.eB delivery of sgRNAs to the brain, with relevance to understanding the molecular underpinnings of neurological and neurodegenerative diseases more broadly. We highlight the power of our CRISPR editing platform for the functional validation of modifier genes identified in human GWAS and for uncovering genetic interactions between modifiers. Our genetic interaction data make predictions that can be further tested in humans in the context of genetic interactions between modifier variants. Significantly, we have identified novel modifiers of somatic *HTT* CAG expansion, the key driver of the rate of HD onset, and provided new insight into underlying pathways that can be dissected by further genetic and biochemical studies. Our data reinforce and extend the pool of potential therapeutic targets to slow somatic expansion, which must also be weighed against their broader safety and druggability profiles (Benn, Gibson and Reynolds, 2021). We highlight MSH3, MLH3, PMS1 and POL8 as potential targets for reducing expression or activity, while FAN1, PMS2 and HMGB1 are possible targets for promoting expression or activity.

## Materials and Methods

### Mouse lines and injection with AAV vectors

Mouse experiments were carried out in accordance with the recommendations in the Guide for the Care and Use of Laboratory Animals, NRC (2010). All animal procedures were carried out to minimize pain and discomfort, under approved IACUC protocols of the Massachusetts General Hospital (MGH) or The Jackson Laboratory (JAX). *Htt*^Q111^ mice on the C57BL/6J (B6J) genetic background (Neto *et al*., 2017) were crossed with Cas9 knock-in mice constitutively expressing Cas9 nuclease in a widespread fashion under the direction of a CAG promoter (JAX strain 026179; B6J) (Platt *et al*., 2014). Male and female mice heterozygous for each knock-in allele and with CAG repeat lengths ranging from 112 to 119 (determined from tail at weaning at Laragen, Inc) were shipped to MGH between 4 and 5 weeks of age. At 6 weeks of age mice were administered via tail vein injection (200 µl) with either 3E+11 AAV8 or 1-3E+12 PHP.eB vector genomes (VG). Mice were aged until either 12 weeks of age for AAV8 injections or until 24 weeks of age for PHP.eB injections, and sacrificed for tissue harvesting. Dual guide targeting experiments for genetic interactions were performed with 3E+11 VG of each of two AAV8s, in a single injection (200 µl) at 6 weeks of age, followed by tissue collection at 12 weeks of age.

### Guide design and *in vitro* testing

Single guide RNAs (sgRNAs) were designed using the Broad Institute design tool (Doench *et al*., 2016; Sanson *et al*., 2018), prioritizing: 1) recommended guidelines of on-target scores >0.6 and target range of 5-65% of the primary transcript; 2) targeting other potential transcripts in addition to the primary transcript; 3) minimizing off-targets. To assess guide efficacy, the top two candidate sgRNA sequences for each gene were cloned into lentiCRISPRv2 (Addgene # 52961) (Sanjana, Shalem and Zhang, 2014). A total of 2.5µg plasmid DNA were transfected into NIH 3T3 cells (∼80-100k) using Lipofectamine 3000 (Thermo Fisher Scientific). After 48-72 hours, transfected cells were selected using 2-3 µg puromycin (Thermo Fisher Scientific) until all cells in non-transfected control were dead. Surviving cells were allowed to recover until 70-90% confluency was achieved. DNA was extracted from surviving cells (Lucigen QuickExtract or Qiagen DNeasy DNA blood and tissue kit) for characterization and quantification of CRISPR-Cas9-induced mutations. The top performing sgRNA for each gene, based on highest frameshift percentage, was selected for AAV packaging and used *in vivo*.

### Quantification of CRISPR/Cas9 editing efficiency

PCR amplicons surrounding the sgRNA target site were generated using Phusion High-Fidelity DNA Polymerase (Thermo Fisher Scientific), purified using MiniElute PCR Purification kit (Qiagen) and subjected to deep 2 x 150 bp paired-end next generation sequencing (NGS) on the Illumina MiSeq platform (DNA core, Center for Computational and Integrative Biology, MGH). Genomic editing was determined in each sample from a minimum of 2,000 NGS reads using CRISPResso v2 (Clement *et al*., 2019), using batch command with default settings and the following parameters: amplicon_min_alignment_score: 90; quantification_window_center: −3; quantification_window_size: 5. Editing efficiency was summarized as percentages of frameshift (non-multiple of 3bp indels) and non-frameshift (multiple of 3bp indels and SNVs) mutations. Guide sequences, editing frequencies and NGS primer sequences are shown in **Tables S1, S2 and S5**.

### AAV constructs and viral production

We generated a plasmid vector for packaging in AAV (pAAV), expressing a sgRNA under the control of the human U6 promoter and an mCherry reporter under the control of the CAGGS promoter (**Fig. S1**), which is a hybrid promoter composed of the CMV immediate-early enhancer, CBA promoter, and CBA intron 1/exon 1 (Niwa, Yamamura and Miyazaki, 1991). sgRNA sequences were cloned into pAAV and then packaged into either AAV8 or PHP.eB capsids at the Massachusetts Eye and Ear Infirmary Gene Transfer Vector Core, the UMass Chan Medical School Viral Vector Core, or SignaGen Laboratories.

### Liver tissue collection and hepatocyte enrichment

Mice were sacrificed by CO2 inhalation and an ∼25 mg piece of liver taken from the tip of the right medial lobe, rinsed in cold phosphate-buffered saline (PBS) and immediately frozen on dry ice or liquid N2 for subsequent DNA extraction for analyses of CAG expansion and guide editing. As needed, an additional 3-5 mm section of liver was cut from the right medial lobe, rinsed in cold PBS and fixed in 10% formalin for histological processing. The remaining liver tissue was rinsed in cold PBS, and frozen on dry ice or liquid N2 for banking for additional analyses as needed. Hepatocyte enrichment was performed similarly to as previously described by us (Lee *et al*., 2011). Briefly, mice were anesthetized with ∼250 mg/Kg avertin (Tribromoethanol, Sigma-Aldrich) by intraperitoneal injection, followed by transcardial perfusion with Dulbecco’s PBS. A small piece of liver was collected for DNA extraction as above for comparison to hepatocyes, prior to transcardial perfusion with 50 ml collagenase solution: 43.24 mg collagenase type IV (Thermo Fisher Scientific), 38.35 mL Leibovitz’s L-15 media (Thermo Fisher Scientific), 6.65 ml 0.15M MOPS buffer at pH 7.2-7.4 (Sigma-Aldrich), and dH2O to final total volume of 50 ml (pre-heated to 37°C). Liver lobes were further incubated in 15 ml of collagenase solution at 37°C for 1-2 hrs with gentle rocking. Cells were pelleted by centrifugation at 100 *g* for 3 minutes, then resuspended in 30 ml PBS and filtered through a 500µm cell strainer. Cells were pelleted by centrifugation at 50 *g* for 3 minutes, resuspended in 30 ml PBS and filtered through a 70 µm cell strainer. Finally, cells were washed four times with 10 ml PBS (50*g* for 3 minutes each cycle) and frozen on dry ice for future DNA extraction.

### *HTT* CAG repeat expansion analysis

Genomic DNA was isolated from tissues using the DNeasy DNA blood and tissue kit (Qiagen). Somatic CAG instability analysis was performed using a human-specific PCR assay that amplifies the *HTT* CAG repeat from the knock-in allele (Mouro Pinto *et al*., 2013). The forward primer was fluorescently labeled with 6-FAM (Thermo Fisher Scientific) and products were resolved using the ABI 3730xl DNA analyzer (Thermo Fisher Scientific) with GeneScan 500 LIZ as internal size standard (Thermo Fisher Scientific). GeneMapper v5 (Thermo Fisher Scientific) was used to generate CAG repeat size distribution traces. CAG expansion indices were calculated from the GeneMapper peak height data as previously described, using a 5% relative peak height threshold cut-off (Lee *et al*., 2010). For AAV8 experiments, expansion indices in both liver and hepatocytes were determined relative to the modal allele in the liver trace from the respective mouse. For PHP.eB, striatum and liver expansion indices were determined relative to the modal allele in the respective trace.

### Western blot analyses

Protein lysates were prepared in RIPA buffer supplemented with 5 mM EDTA and protease inhibitors (Halt Protease Inhibitor Cocktail, Thermo Scientific) by mechanical grinding with disposable pestle and two 10-second sonication pulses (Branson sonifier, power level 3.5), on ice. The homogenates were kept on ice for 20 min and then clarified by centrifugation at 4 °C for 30 minutes at 14000 rpm. Protein concentration was determined using the DC protein assay kit (Bio-Rad). Protein extracts were resolved on 4–12 % Bis-Tris polyacrylamide gels (NuPAGE, Life Technologies) and transferred to 0.45 mm nitrocellulose membrane (Thermo Scientific). The membranes were blocked with 5% dry non-fat milk in TBST (Tris-buffered saline with 0.1% Tween 20) and incubated with primary antibody diluted in 5% dry non-fat milk/TBST. The following primary antibodies were used: MLH1 (1:1000, ab92312, Abcam), MSH2 (1:1000, D24B5, Cell Signaling), MSH3 (1:5, 2F11, conditioned medium from hybridoma obtained from DSHB), MSH6 (1:1000, ab92471, Abcam). Horseradish peroxidase-conjugated goat anti-rabbit and anti-mouse (1:5000; NA934 and NA931 respectively, GE Healthcare) were used as secondary antibodies. Signals were visualized using enhanced chemiluminescence (ECL) detection system (Thermo Scientific). Densitometric analysis of protein levels was performed using Quantity One software (Bio-Rad). Following background subtraction, protein levels were normalized by total protein amount in the same lane as determined by Novex Reversible Membrane Protein Stain (Thermo Scientific) performed prior to blocking.

### Histological analyses

Formalin-fixed fragments of the right medial liver lobe were cryo-embedded in OCT, sectioned at 6 mm, and mounted on glass slides. Liver AAV transduction efficiency was evaluated by fluorescence microscopy for mCherry with DAPI counterstain, with auto-stitching of 10x images captured on Leica DMi8 epifluorescence microscope. For overall evaluation of liver toxicity, slides were stained with Hematoxylin and Eosin or Ki67 antibody. For Ki67 immunostaining, the slides were treated for 30 min with 0.3% H_2_O_2_ in methanol, blocked for 1 h with 3% normal horse serum (NHS) in TBS, incubated overnight at 4°C with Ki67 antibody (1:200 in 1% NHS/TBS, ab16667, Abcam), then for 1 h with biotinylated goat anti-rabbit secondary antibody (1:200, BA-1000, Vector Laboratories), followed by standard avidin-biotin complex (ABC) staining (VECTASTAIN^®^ Elite^®^ ABC Kit, Vector Laboratories) with DAB chromogenic substrate (Sigma-Aldrich). Counterstain for Ki67 was done with methyl green (Sigma-Aldrich). Brightfield microscopy was performed with Olympus BX51 microscope (PlanFL N 10x/0.30 objective), equipped with Q-Color 5 Olympus camera and QCapture Pro image acquisition software.

### Statistical analyses

Z-scores of liver CAG expansion indices were calculated for each of a total 620 *Htt*^Q111^.Cas9 mice by subtracting the global average expansion index (3.954) followed by dividing by the standard deviation (0.9709). Statistical significance was determined by comparing the mean of each group with the mean of the “empty” group by one-way ANOVA, with Dunnett’s multiple comparison correction, using GraphPad Prism v10. Summary statistics and p-values are shown in **Tables S2 and S5**.

For the genetic interactions, statistical significance was determined by comparing raw expansion indices in an “all-by-all” one-way ANOVA, with Dunnett’s multiple comparison correction, using GraphPad Prism v10 (**Table S6**).

## Data Availability

The datasets generated during and/or analyzed during the current study are available on reasonable request.

## Acknowledgments

This research was supported by: the Berman/Topper Family HD Career Development Fellowship from the Huntington’s Disease Society of America (R.M.P.); Young Investigator Award from the National Ataxia Foundation (R.M.P.); Harvard NeuroDiscovery Center grant (R.M.P.); National Institutes of Health grants to R.M.P. (R01 NS126420), V.C.W. (R01 NS049206; R21 NS111066) and J.F.G. (R01 NS091161); CHDI Foundation, Inc. (V.C.W, M.E.M and J.F.G.); and Pfizer, Inc (R.M.P and V.C.W.). We thank: the MGH CCIB DNA Core for assistance with Next-Generation Sequencing; the UMass Chan Medical School Viral Vector Core and the MEEI Gene Transfer Vector Core for AAV production; the Rodent Histopathology Core at Dana-Farber/Harvard Cancer Center for tissue OCT embedding, cryosectioning and H&E staining services (Dana-Farber/Harvard Cancer Center is supported in part by a NCI Cancer Center Support Grant # NIH 5 P30 CA06516); Stephani Dempsey and Britt Callahan from The Jackson Laboratory for assistance with breeding and genotyping the Cas9.Q111 mice; Brenda Lager at CHDI for mouse logistics support; Julieanne Brandolini and Zahara Yee from MGH CCM for technical support; and Feng Zhang and Randall Platt from the Broad Institute for sharing the Cas9 mice and early discussions; Anna Pluciennik, Seung Kwak and Ramee Lee for helpful discussions. Figures 1 and 6 were partly generated using BioRender.

