## Supplementary Figures for "Identification of genetic modifiers of Huntington’s disease somatic CAG repeat instability by in vivo CRISPR-Cas9 genome editing"

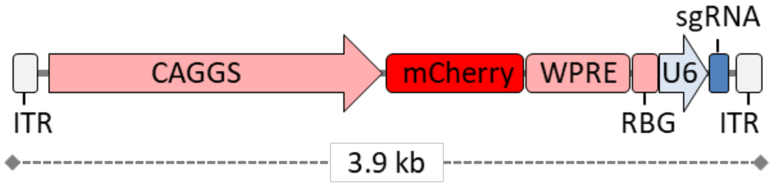


**Figure S1. AAV vector expressing sgRNA and mCherry reporter**

ITR, Inverted Terminal Repeat; CAGGS, hybrid promoter composed of the CMV immediate-early enhancer, CBA promoter, and CBA intron 1/exon 1 (Niwa, Yamamura and Miyazaki, 1991); WPRE, Woodchuck Hepatitis Virus Posttranscriptional Regulatory Element; RBG, rabbit beta-globin polyadenylation signal; U6, human U6 promoter; sgRNA, single guide RNA.

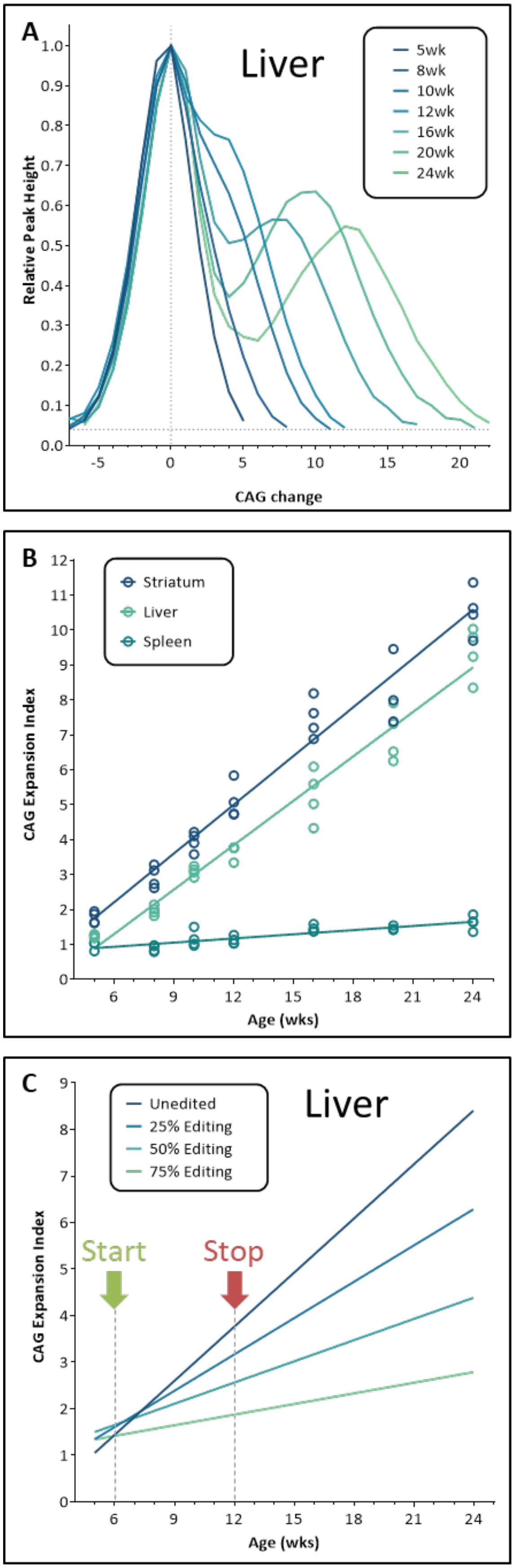


**Figure S2. CAG expansion trajectories over time and simulated impact of modifier gene editing**

**A.** Normalized GeneMapper peak height data from *Htt* CAG repeat-containing PCR products from liver, averaged across mice (N=1-4) at different ages. **B**. CAG expansion indices from liver, striatum and spleen from the same mice (N=3-4). **C**. Genomic DNA mixing experiment to simulate editing. Liver genomic DNA from 5-week mice (stable) was mixed in different proportions with liver genomic DNA from mice between 9 and 24 weeks of age. Proportions were 0% 5-week:100% other, 25% 5-week:75% other; 50% 5-week:50% other, and 75% 5-week:25% other, simulating the impact of no editing or 25%, 50% and 75% bi-allelic editing of a strong-effect modifier gene suppressing expansion. These results were used to guide the experimental paradigm of injecting AAV8 at 6 weeks of age (Start) and analyses of CAG expansion at 12 weeks of age (Stop). Analyses of 12-week mice permits relatively fast *in vivo* analyses, minimizes variation in expansion between untreated animals that increases with age (panel B) and is predicted to provide the sensitivity to detect the impact of a strong effect modifier suppressing expansion with an editing efficiency as low as 25%.

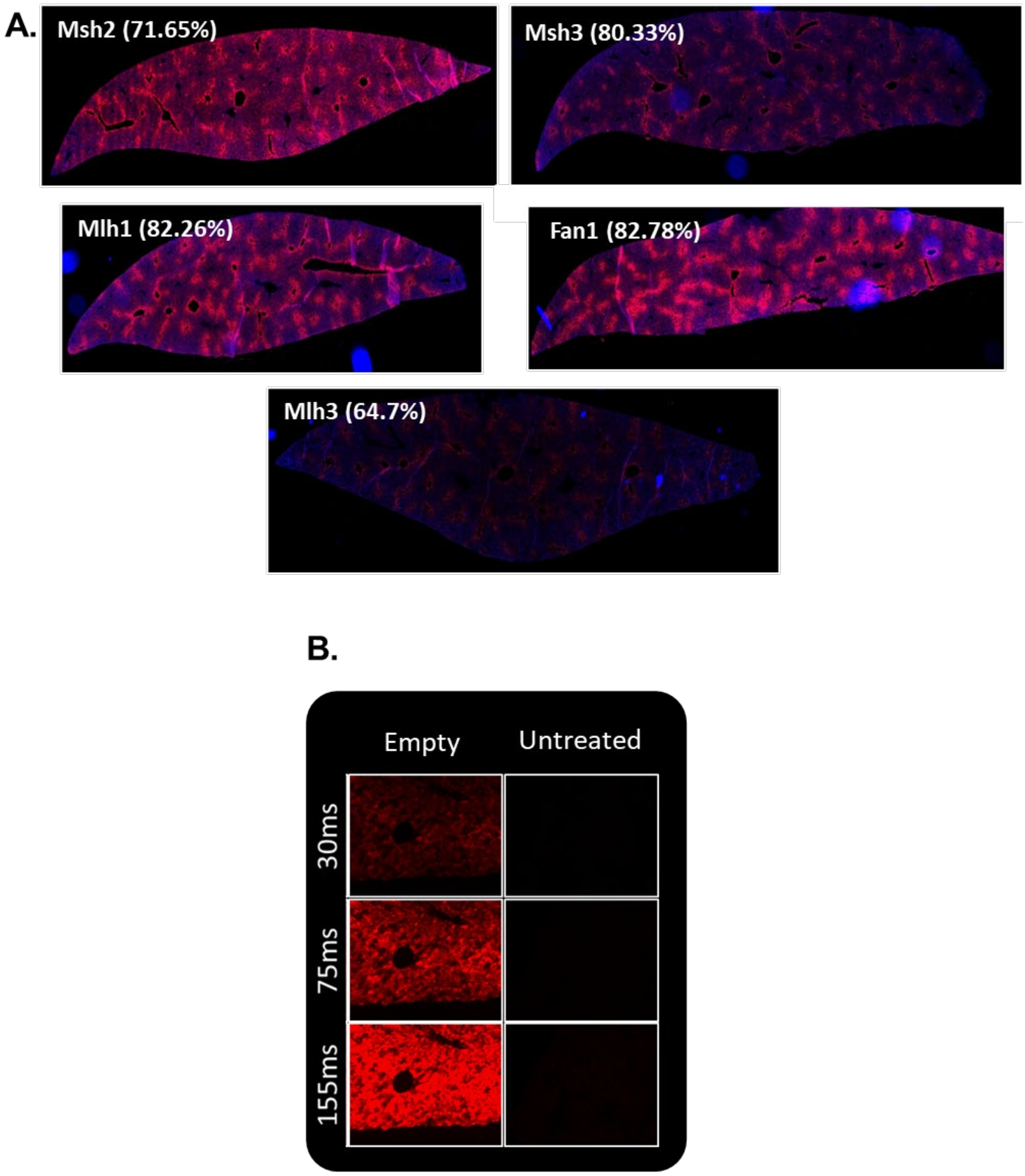


**Figure S3. AAV8 transduction efficiency in the liver**

**A**. mCherry immnofluorescence in liver sections from 12-week *Htt*$^{Q111}$ Cas9 mice treated at 6 weeks with AAV8 vectors expressing mCherry and sgRNAs targeting various modifier genes. The % of gene editing (frameshift + non-frameshift) is shown. **B**. Magnified mCherry immunofluorescence images at different exposure times from empty vector-treated or untreated mice.

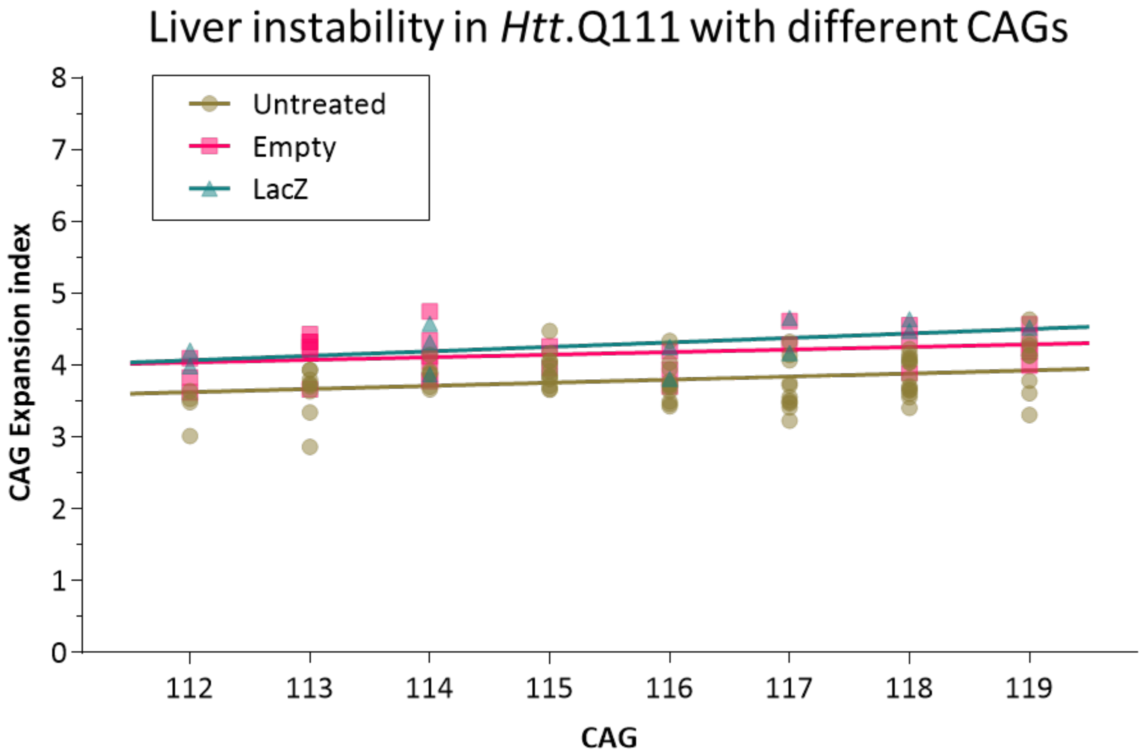


| Simple Linear Regression | Untreated | Empty | LacZ |
|---|---|---|---|
| Number of animals (*n*) | 77 | 30 | 12 |
| Goodness of Fit ($R^2$) | 0.07896 | 0.08743 | 0.2464 |
| Slope | 0.04348 | 0.03567 | 0.06203 |
| Is slope significantly non-zero? | | | |
| F | 6.430 | 2.682 | 3.270 |
| P value | 0.0133 | 0.1127 | 0.1007 |
| Deviation from zero? | Significant | Not Significant | Not Significant |

**Figure S4. Relationship of CAG expansion in the liver to inherited repeat length**

Liver expansion indices measured at 12 weeks versus inherited CAG length (determined in tail at weaning at Laragen) in untreated, mice and control (empty vector and LacZ)-treated mice over the range of CAGs (112-119) used in this study. The lack of/very minimal impact of inherited CAG length in this range (apparent to the greatest extent in the untreated mice due to the large number of mice analyzed) indicates that CAG length variation within this range does not confound our interpretation of the effects of modifier genes.

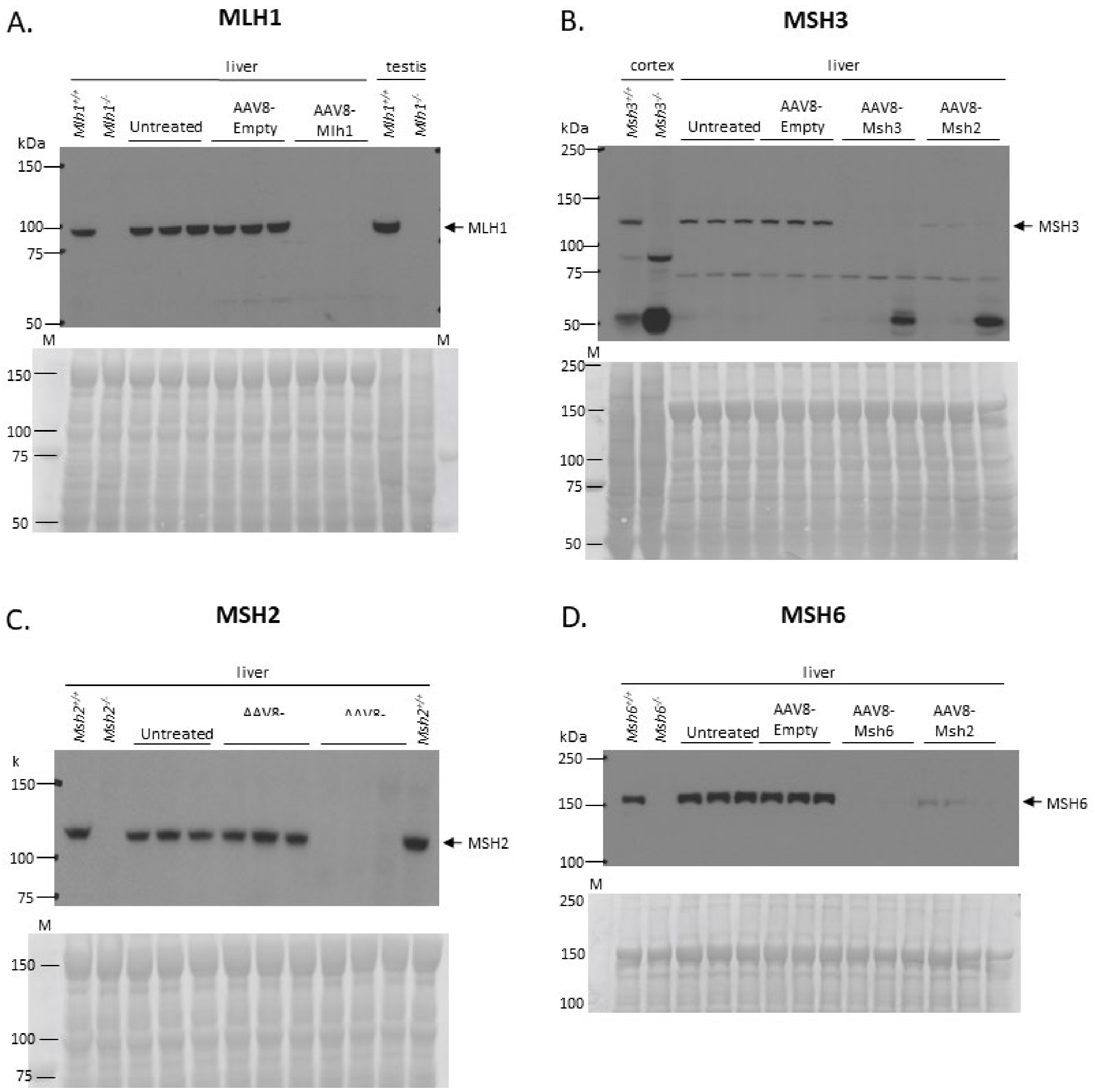


**Figure S5. Western blot analyses**

Western blot analyses in 12-week livers from $\mathit{Htt}^{Q111}$ Cas9 mice, either untreated or treated at 6 weeks with empty AAV8 vector or AAV8 vectors expressing sgRNAs targeting *Mlh1, Msh3, Msh2* or *Msh6*. Western blots are probed with antibodies to MLH1 (A), MSH3 (B), MSH2 (C) or MSH6 (D). Also loaded as controls are extracts from 6-month liver, cortex or testis of constitutional null mice and their littermate controls ($\mathit{Mlh1}^{-/-}$, $\mathit{Mlh1}^{+/+}$, $\mathit{Msh3}^{-/-}$, $\mathit{Msh3}^{+/+}$, $\mathit{Msh2}^{-/-}$, $\mathit{Msh2}^{+/+}$, $\mathit{Mshs6}^{-/-}$, $\mathit{Msh6}^{+/+}$) (Dragileva *et al.*, 2009; Kovalenko *et al.*, 2012; Mouro Pinto *et al.*, 2013). Total protein stain (Novex) is shown under each western blot (A=75 µg, B=75 µg, C=60 µg, D=35 µg total protein/well). In panels B and D, CRISPR-Cas9 knockout of *Msh2* also results in dramatically reduced levels of MSH3 and MSH6, respectively, due to the destabilization of MutSβ and MutSα dimers.

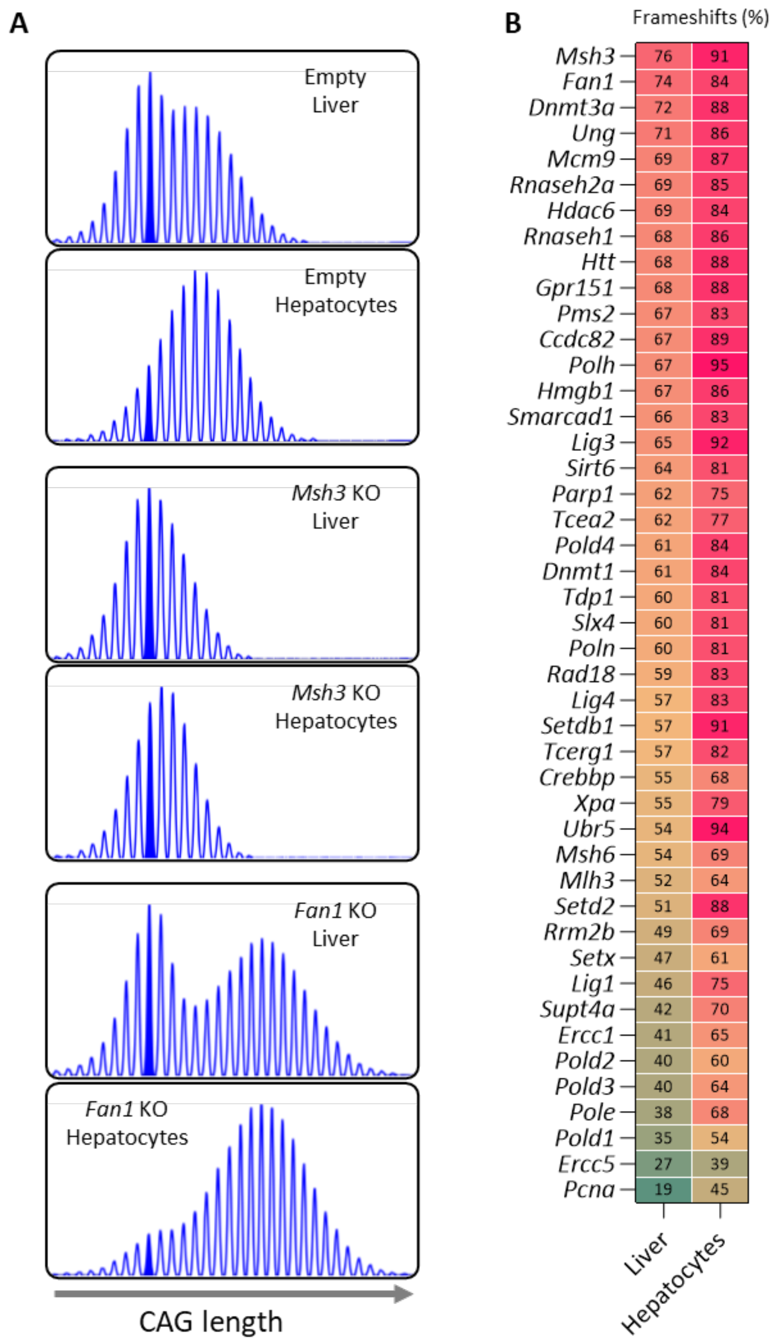


**Figure S6. Enhanced editing in hepatocytes relative to whole liver**

**A.** Example GeneMapper traces from paired hepatocyte and liver DNAs of empty vector, *Msh3* guide and *Fan1* guide-treated mice. The modal peak in liver is filled in blue. **B**. Mean expansion % frameshift mutation in liver compared to hepatocytes. Refer also to **Table S2.**

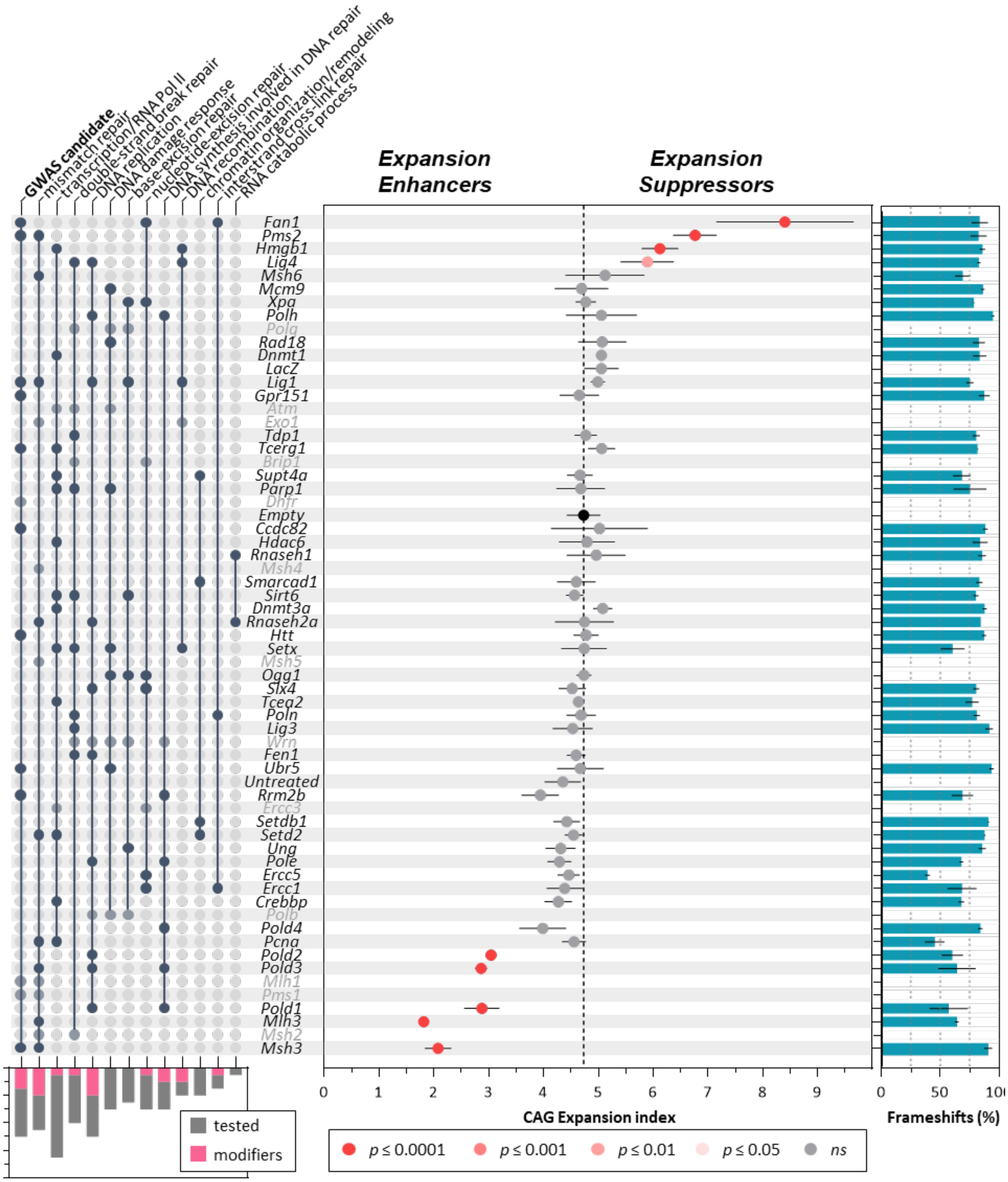

GWAS candidate
mismatch repair
transcription/RNA Pol II
double-strand break repair
DNA replication
DNA damage response
base-excision repair
nucleotide-excision repair
DNA synthesis involved in DNA repair
DNA recombination
chromatin organization/remodeling
interstrand cross-link repair
RNA catabolic process
Expansion Enhancers
Expansion Suppressors
Fan1
Pms2
Hmgb1
Lig4
Msh6
Mcm9
Xpa
Polh
Polq
Rad18
Dnmt1
LacZ
Lig1
Gpr151
Atm
Exo1
Tdp1
Tcerg1
Brip1
Supt4a
Parp1
Dhfr
Empty
Ccdc82
Hdac6
Rnaseh1
Msh4
Smarcad1
Sirt6
Dnmt3a
Rnaseh2a
Htt
Setx
Msh5
Ogg1
Slx4
Tcea2
Poln
Lig3
Wrn
Fen1
Ubr5
Untreated
Rrm2b
Ercc3
Setdb1
Setd2
Ung
Pole
Ercc5
Ercc1
Crebbp
Polb
Pold4
Pcna
Pold2
Pold3
Mlh1
Pms1
Pold1
Mlh3
Msh2
Msh3
tested
modifiers
0 1 2 3 4 5 6 7 8 9
CAG Expansion index
0 50 100
Frameshifts (%)
p ≤ 0.0001
p ≤ 0.001
p ≤ 0.01
p ≤ 0.05
ns

**Figure S7. Expansion indices in enriched hepatocytes**

Mean±s.d CAG expansion indices in hepatocytes enriched from livers of 12-week *Htt*$^{Q111}$ Cas9 mice treated at 6 weeks of age with AAV8 expressing sgRNA targeting candidate genes of interest and including empty vector, LacZ and untreated controls. Enriched hepatocytes were analyzed in a subset of mice for the majority of genes tested. The order of genes/conditions represented matches that in Figure 3. Adjusted p-values are determined in a one-way ANOVA with Dunnett's multiple comparison test relative to empty vector. The bar graph on the right shows the % frameshift mutation (mean±s.d) for each targeted gene. See **Tables S2 and S3** for details. The panel on the left indicates major Gene Ontology (GO) Biological Processes represented by these genes and "GWAS Candidate" indicates candidate genes at genome-wide significant age at onset modifier loci (GeM-HD, 2019). To minimize redundancy, "transcription/regulation by RNA Polymerase II" is combined from GO terms "negative regulation of transcription by RNA polymerase II", "positive regulation of transcription by RNA polymerase II" and "transcription elongation from RNA polymerase II", and "chromatin organization/remodeling" is combined from GO terms "chromatin organization" and "chromatin remodeling". **Table S3** shows the full set of GO Biological Processes and **Table S4** indicates the rationale for gene inclusion. The bottom left bar graph indicates, for each pathway, the number of genes modifying expansion (adjusted $p<0.05$) relative to the number of genes tested.

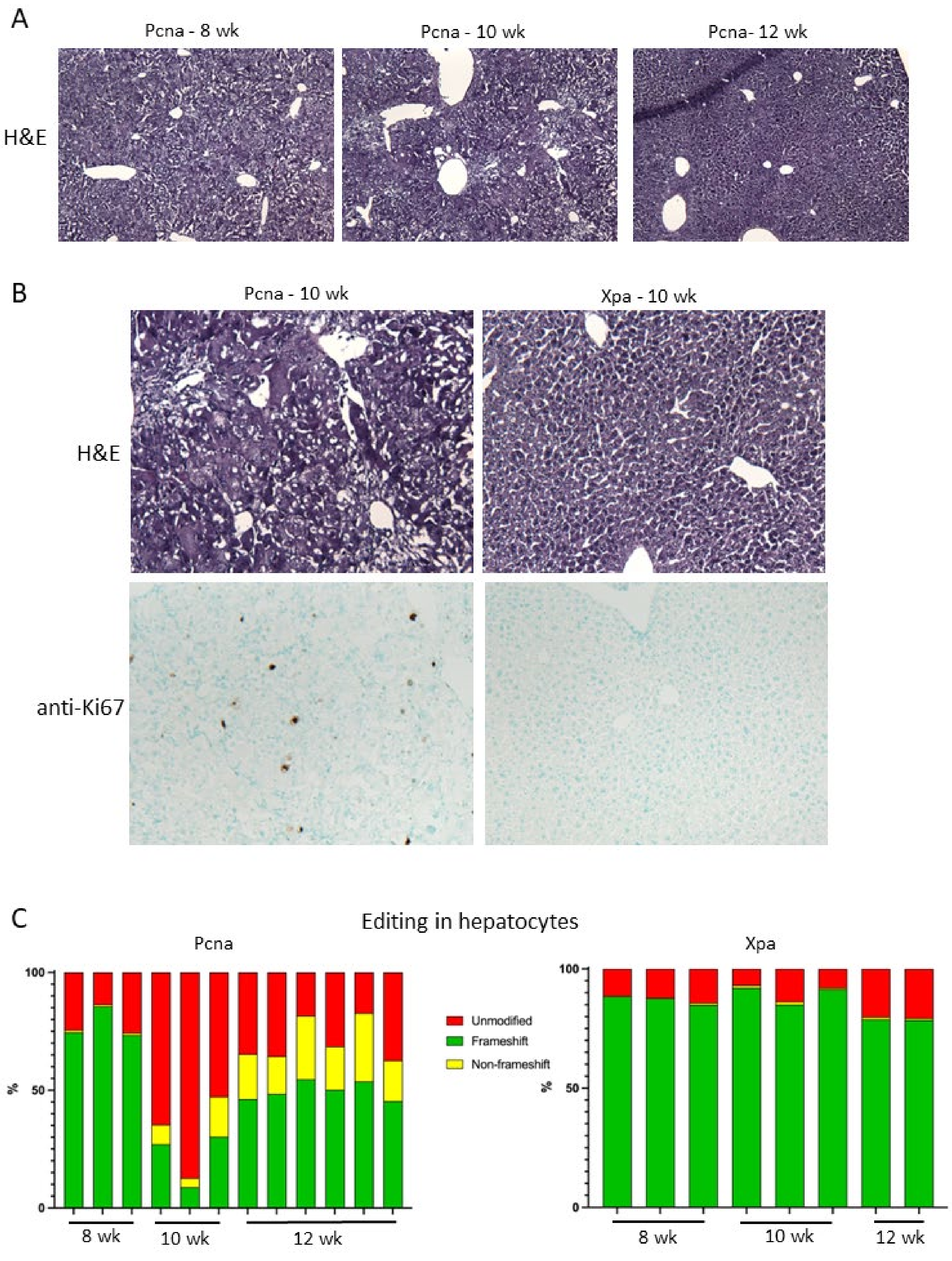
A
Pcna - 8 wk
Pcna - 10 wk
Pcna- 12 wk
H&E
B
Pcna - 10 wk
Xpa - 10 wk
H&E
anti-Ki67
C
Editing in hepatocytes
Pcna
Xpa
Unmodified
Frameshift
Non-frameshift
%
100
50
0
8 wk
10 wk
12 wk

**Figure S8. The impact of CRISPR-Cas9 editing of *Pcna***

**A**. Representative images of Hematoxylin and Eosin (H&E) staining performed on liver sections of $\mathit{Htt}^{Q111}$ Cas9 mice at 8 (N=3), 10 (N=4) and 12 weeks (N=6) of age treated with AAV8 targeting *Pcna* at 6 weeks of age. At 12 weeks of age the tissue appears largely intact, with hepatocyte nuclei clearly discernable. At younger ages mice show evidence of hepatocyte/liver damage with indistinct nuclear morphology and disorganized tissue structure with enlarged trabeculae. **B**. Zoomed in images of H&E staining in 10-week *Pcna* guide-treated liver compared to 10-week *Xpa* guide-treated liver as a control, and immunostaining for Ki67, a proliferation marker, in the same mice. In *Pcna* guide-treated mice, Ki67-positive cells were observed at 10 weeks, to a lesser extent at 8 weeks, were largely absent at 12 weeks and were not observed in *Xpa*-guide treated livers at any age. **C**. Editing in hepatocytes over the 8>10>12-week time course in *Pcna*- and *Xpa*-guide treated mice. CRISPR-Cas9 targeting of *Pcna*, but not *Xpa,* resulted in a shift from a high percentage of frameshift mutations (8 weeks), to a majority of unedited reads (10 weeks) and then to an increased fraction of edited reads with a relatively high proportion of non-frameshift mutations (12 weeks). Taken together with the histological analyses, these data indicate that CRISPR-Cas9 targeting of *Pcna* results in hepatocyte cell degeneration and regeneration. Patterns of gene editing over time may reflect the degeneration of hepatocytes harboring homozygous loss of function mutations and the subsequent selection in the regenerating liver of hepatocytes harboring either heterozygous loss of function mutations or non-frameshift mutations compatible with cell division.

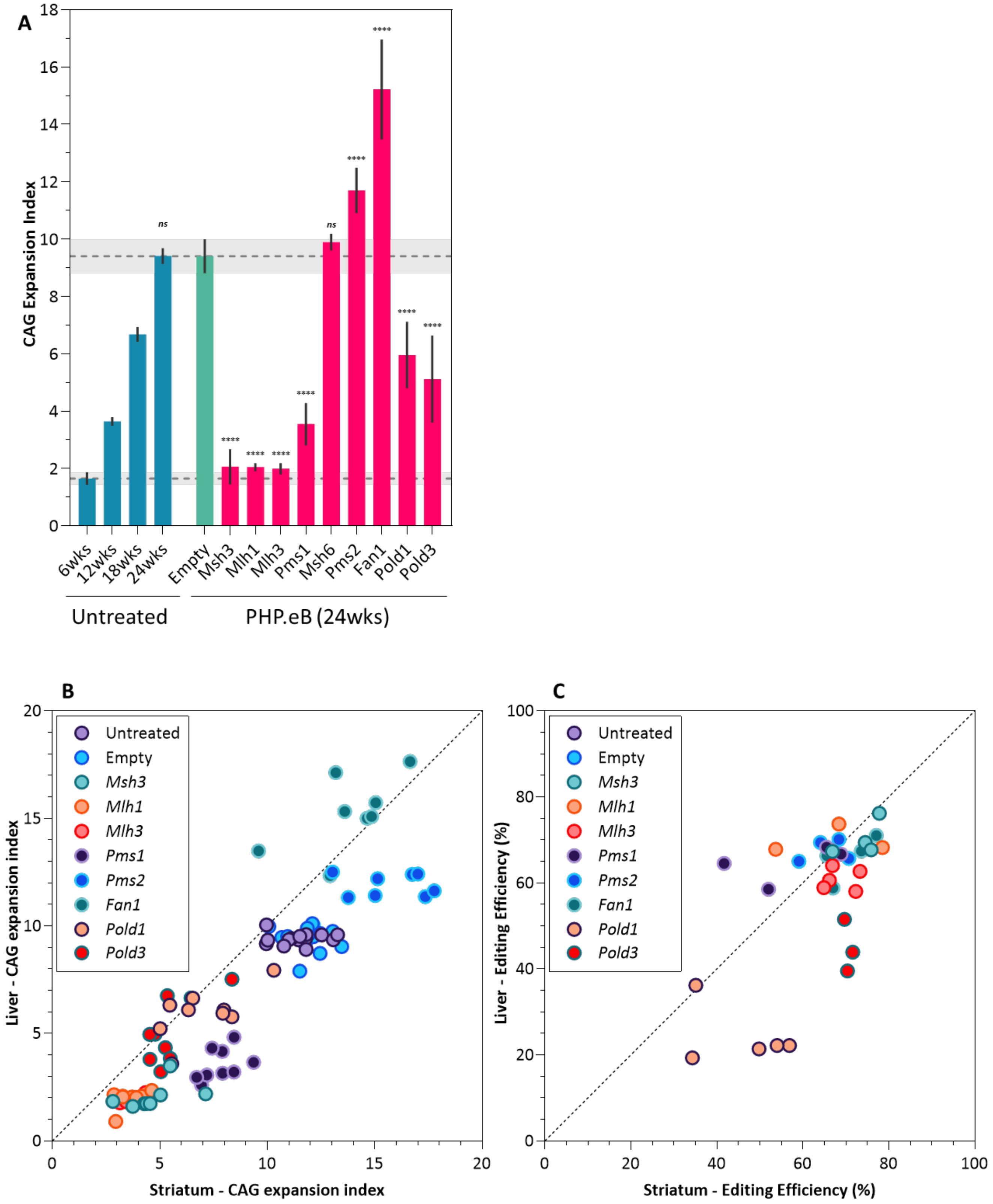

A
CAG Expansion Index
0
2
4
6
8
10
12
14
16
18
ns
ns
****
6wks
12wks
18wks
24wks
Empty
Msh3
Mlh1
Mlh3
Pms1
Msh6
Pms2
Fan1
Pold1
Pold3
Untreated
PHP.eB (24wks)
B
Liver - CAG expansion index
Striatum - CAG expansion index
0
5
10
15
20
Untreated
Empty
Msh3
Mlh1
Mlh3
Pms1
Pms2
Fan1
Pold1
Pold3
C
Liver - Editing Efficiency (%)
Striatum - Editing Efficiency (%)
0
20
40
60
80
100
Untreated
Empty
Msh3
Mlh1
Mlh3
Pms1
Pms2
Fan1
Pold1
Pold3

**Figure S9. Liver expansion in PHP.eB injected mice**

**A.** Somatic CAG expansion indices, determined from fragment sizing of *Htt* CAG repeat-containing PCR amplicons of untreated *Htt*$^{Q111}$ Cas9 mice livers from 6 to 24 weeks of age, and in 24-week mice injected at 6 weeks of age with control PHP.eB (empty vector) or PHP.eB expressing sgRNAs targeting genes of interest. **** $p<0.0001$ relative to empty vector (One-way ANOVA with Dunnett's multiple comparison correction) (**Table S5**). Bars show mean±s.d. Dotted lines/shaded grey regions show mean/95% confidence interval expansion indices in 6-week untreated mice and in empty vector control-treated mice at 24 weeks of age. **B**. Liver CAG expansion indices plotted against striatal expansion indices determined in the same 24-week mice. Dotted line = line of identity. Expansion indices tend to be lower in the liver than in the striatum (points to the right of the identity line), with the exception of *Fan1* guide-treated mice where expansion indices are greater in the liver (points to the left of the identity line), indicating a relatively large expansion-promoting impact of *Fan1* knockout in this tissue. **C**. CRISPR editing efficiencies in liver plotted against striatum as determined in the same 24-week PHP.eB-treated mice. Dotted line = line of identity. Editing efficiency seems comparable between the two different tissues, with exception for *Pold1* and *Pold3* which seem to have weaker editing in the liver (points to the right of the identity line), potentially due to POLD's role in replication.

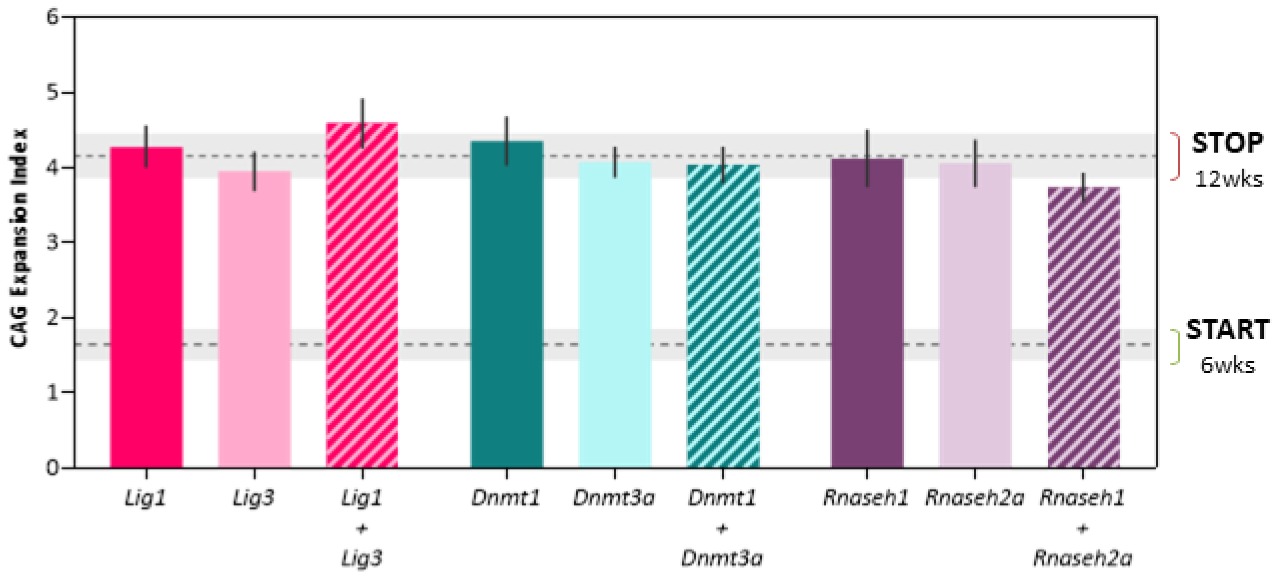


**Figure S10. Dual guide targeting of *Dnmt1/Dnmt3a*, *Rnaseh1/Rnaseh2a* and *Lig1/Lig3***
Potential functional redundancies were tested in dual guide experiments targeting *Dnm1+Dnmt3*, *Rnaseh1+Rnaseh2a* and *Lig1+Lig3*. Bars show mean±s.d CAG expansion indices, of *Htt*$^{Q111}$ Cas9 mice livers at 12 weeks of age following injection with either one (single target) or two (dual target) AAV8s. Dotted lines/shaded grey regions show mean/95% confidence interval expansion indices in 6-week untreated mice at and in empty vector control-treated mice at 12 weeks of age.
