## Supplementary Tables for "Identification of genetic modifiers of Huntington’s disease somatic CAG repeat instability by in vivo CRISPR-Cas9 genome editing"

***Supplementary Table 1: Genes targeted, CRISPR guides and Next Generation MiSeq (NGS) assay primers used***

| Gene | sgRNA ID | sgRNA sequence (5' - 3') | sgRNA coordinates (GRCm39/mm39) | Exon targeted (target cut %) | NGS assay | |
|---|---|---|---|---|---|---|
| | | | | | F primer (5'-3') | R primer (5'-3') |
| *Atm* | Atm-JD3.1 | TGCAAGATACACATGAATCG | chr9:53,390,984-53,391,003 (-) | 37/63 (61%) | TGCCATGTCCTAGGTGAAAACC | GCCCTGTAACTTACGGAACCA |
| *Brip1* | Brip1-JD2.6 | ACTGTTAAGAAAGATCACGG | chr11:86,048,619-86,048,638 (-) | 7/20 (20%) | TGATGCGTTCTGAGAGATGTTCT | TGTGTCCGCGTCCCAAAATA |
| *Ccdc82* | Ccdc82-JD3.1 | ATCACTGTCATACATCACAG | chr9:13,252,194-13,252,213 (-) | 3/9 (31%) | ATGATGAAGCTCACCCTGGTC | ACATTCATCCTCGACCACCCT |
| *Crebbp* | Crebbp-JD3.2 | CATACTCGAGATATCCGATG | chr16:3,911,349-3,911,368 (+) | 26/31 (60%) | AATGACACCGCTTTCTATCCT | TCAGCAAACTTCTGAACCAACC |
| *Dhfr* | Dhfr-JD3.1 | GACATGGTTTGGATAGTCGG | chr13:92,502,261-92,502,280 (+) | 4/6 (62%) | TGCCAAAAGTTTGGATGATGCC | CCCTTTGGCACCTGTGTAGT |
| *Dnmt1* | Dnmt1-JD3.1 | CATCACGGCTCACTTCACGA | chr9:20,851,776-20,851,795 (-) | 5/39 (10%) | GCCACCTGCAGTCACTTGTT | CAGGGACATCCGCTCAGAC |
| *Dnmt3a* | Dnmt3a-JD3.1 | GCACTGCAGAAGGAGCTCAG | chr12:3,946,602-3,946,621 (-) | 9/23 (38%) | CAGGGAAGGACCGATCTAAGG | TTGGCTAACACAGCTCCGA |
| *Ercc1* | Ercc1-JD3.1 | TGTGCCCTGGGAATTCGGTG | chr7:19,084,118-19,084,137 (+) | 4/10 (41%) | GTCACCCCCATAAGTGGCTT | TAACTGTAGGCCACAACCGC |
| *Ercc3* | Ercc3-JD2.1 | TGTCGCTTGAGGAACGCCGA | chr18:32,378,609-32,378,628 (+) | 5/15 (26%) | CCCACCCTGATGTTATCCAGC | ACCATGAACTGAAGCTGTGCT |
| *Ercc5* | Ercc5-JD3.2 | GTAAAGGAGTGTGTAACCAG | chr1:44,206,436-44,206,455 (+) | 8/15 (39%) | GAGGAGCATGCTGATGAAACG | TGCTTCTTTCACGGCACGAG |
| *Exo1* | Exo1-JD2.1 | TCTTACAGCCAAATGCGAGG | chr1:175,716,406-175,716,425 (-) | 6/15 (21%) | CCTAGGAGTGGATTGCCTCG | ACGAAGATTTTAGCTCCTCAGCA |
| *Fan1* | Fan1-JD2.1 | ACATGTCTGACATGACTACG | chr7:64,022,241-64,022,260 (-) | 2/14 (33%) | CCAAAACGCTGGTGTCTGG | CTCCGCAGGTAGTACGGGT |
| *Fen1* | Fen1-JD3.1 | GAACTGGTAGATGCTCATGG | chr19:10,178,317-10,178,336 (+) | 3/3 (10%) | TGCTACCATGGGAATTCACGG | CATGCGGATGGTACGGTAGAA |
| *Gpr151* | Gpr151-JD3.1 | AGCACCACCAGACGTCACGC | chr18:42,712,120-42,712,139 (-) | 1/1 (44%) | GCTTCGCGTATGCAAGTGAC | CGCCAGCCAGAAGTAAAGGA |
| *Hdac6* | Hdac6-JD3.2 | TCCCTGCACCGGTATGACCG | chrX:7,802,469-7,802,488 (-) | 22/29 (59%) | TGAGGATGACCCTAGGTGAGT | ATCTCGGCCTACTTGGCTAC |
| *Hmgb1* | Hmgb1-JD3.2 | TCATAAGCTCATACTCACGG | chr5:148,987,087-148,987,106 (+) | 3/5 (46%) | AGGCTGACAAGGCTCGTTATG | CCCCTCAAATACACGACCCAA |
| *Htt* | Htt-ex6.1 | GACCCGAACAAGCAAAAGAC | chr5:34,956,751-34,956,770 (+) | 6/67 (6%) | CCTGACAGGGAACCATGAGC | ACTCATGCCAAGAGGCAACA |
| *LacZ* | LacZ-JD3.1 | TGCGAATACGCCCACGCGAT | | | | |
| *Lig1* | Lig1-JD3.2 | AGAGTGATTCTCCAGTGAAG | chr7:13,019,218-13,019,237 (+) | 4/29 (6%) | GGAAGAGAAGGGAATGCAGAGT | CAGGGGCTTCATCTTCGTCC |
| *Lig3* | Lig3-JD3.1 | GCTGCTCTGCCATCGCACAG | chr11:82,674,433-82,674,452 (-) | 2/20 (9%) | AACAAATCTCCTCGGTGGGT | CTTTGCCAATGCGGCACAC |
| *Lig4* | Lig4-JD3.1 | TTAGACGTCCTAATTGTGGG | chr8:10,022,376-10,022,395 (-) | 2/2 (51%) | ACCCTCTGTCCATTTACAAGCC | CGGCACAGAGTGTGGAATACA |
| *Mcm9* | Mcm9-JD3.1 | TGAATAATGAGCAATCCTCG | chr10:53,492,024-53,492,043 (-) | 5/13 (24%) | CCTCACCATCTATGGGGTTGTA | TAGGCCCTTCATCACCAACCT |
| *Mlh1* | Mlh1-JD4 | AGGCAGCTGATTCTACCAGA | chr9:111,078,358-111,078,377 (+) | 10/19 (35%) | GTTCCTGGGCATCTGATAAGG | TACAGGAATGGGTGTGTGTTT |
| *Mlh3* | Mlh3-JD2 | CATTGATTACATATATCCCG | chr12:85,315,257-85,315,276 (+) | 2/12 (22%) | TTATATGCAGGCCAAAGAATGGC | CTTCCTGAATACAAATCAACACGGT |
| *Msh2* | Msh2-JD1 | GCGCCGTGTAAAAGTCGCCG | chr17:87,980,140-87,980,159 (-) | 1/16 (4%) | AAATGGCGGTGCAGCCTAAG | ATGTACTTGATCACGCCCTGG |
| *Msh3* | Msh3-JD2.3 | CTTACTCCGAGCACTCATCG | chr13:92,411,305-92,411,324 (-) | 14/24 (57%) | TTTGGTTTCTAGTGCTCTACCCA | GGAAGCCCGCACTTACTTGG |
| *Msh4* | Msh4-JD3.1 | CTGGATGAAGAACCCCCGTG | chr3:153,580,351-153,580,370 (+) | 12/20 (58%) | GGAATGATAGCACAGCTCGC | GCCTGGAAAGTCTACCGTCAA |
| *Msh5* | Msh5-JD3.1 | TGGCTACTTTGTACACCGAG | chr17:35,261,104-35,261,123 (+) | 9/25 (29%) | GGGATGCATAGAAATTCAGACTCC | CCTGCGCATGGCTATAGAGT |
| *Msh6* | Msh6-JD2.2 | CATCAGTGACCGTCTAGATG | chr17:88,293,598-88,293,617 (+) | 4/10 (58%) | GGATACTTGCCATACGCCCTTTG | CAGGGGAGACCCAACATTATGA |
| *Ogg1* | Ogg1-JD3.1 | AAGAGAAAAGGCACTCAGTG | chr6:113,306,215-113,306,234 (-) | 3/7 (40%) | TGATTTCTCGACCCGAGCAG | ATGAGTCGAGGTCCAAAGGC |
| *Parp1* | Parp1-JD3.1 | GCTGTGATTGAGAAACTCGG | chr1:180,414,894-180,414,913 (+) | 9/23 (41%) | TTGAGTGGTCGTAGCCTGTTG | ATCAGGCTACAGTGGTGAGTC |
| *Pcna* | Pcna-JD3.1 | ATACGTGCAAATTCACCCGA | chr2:132,093,306-132,093,325 (+) | 4/6 (54%) | GTCTTGCATTCTGAGTTCTGGTTA | GCACAGGATATCACAACAGCATC |
| *Pms1* | Pms1-JD2.2 | GGGGTCAATATGTAATGTTG | chr1:53,314,234-53,314,253 (-) | 3/13 (11%) | GTGAGGGCATCAAGGCTGTA | AAGCTACTCTGACAAACAATGAAT |
| *Pms2* | Pms2-JD2.2 | AGTTTCAGACAATGGATGTG | chr5:143,850,257-143,850,276 (+) | 3/15 (8%) | AGAGTGTAGGCATGTCTGCTAA | TCTGAGACACGTGGAAATGACT |
| *Polb* | Polb-JD3.1 | ACTATAGGCGCAGAGTCGAG | chr8:23,130,008-23,130,027 (-) | 10/14 (56%) | TCTAGGTTGGCCTAATTAGTAGGTT | CCTACATGGTGAAAATACCTAACCA |
| *Pold1* | Pold1-JD3.1 | GTTCATCAAGACACCCACTG | chr7:44,187,495-44,187,514 (-) | 16/27 (58%) | GGAGCTGGGTAGTTCGGAAG | TCATCTAAGGCAGCCACAGC |
| *Pold2* | Pold2-JD3.1 | GGGTGGATATATTTGCTCCG | chr11:5,824,841-5,824,860 (+) | 4/11 (26%) | ACGTGAGCTCTGAGACTCCT | GCTGCAATGGGCCTCTCTAA |
| *Pold3* | Pold3-JD3.1 | TCTGGCAGTCTTATACAGAA | chr7:99,761,487-99,761,506 (-) | 3/12 (15%) | GAGTCACTTCTGTCTTGTTTGGC | ATACCATTCACTCTTCCTTCTTCC |
| *Pold4* | Pold4-JD3.2 | CACCCAGTCCCTCAGCCAGG | chr19:4,282,524-4,282,543 (+) | 2/4 (37%) | GGATGGCAAAGGCAAATGGG | GAGGGACTGGAAATGGGGTG |
| *Pole* | Pole-JD3.1 | CATTATGGTCACCTACAATG | chr5:110,443,144-110,443,163 (+) | 11/49 (16%) | AGTCCTGGCAAGTTGCTGAA | TGAGAAAGCCACCAACAGCA |
| *Polh* | Polh-JD3.1 | TGTCACTAGAAACATGTGGG | chr17:46,509,639-46,509,658 (-) | 3/11 (9%) | TTGAACTGCTGTGTCTGGGATTG | TTCCACGGGACTCACGAACTT |
| *Poln* | Poln-JD3.1 | TGAAATGATAGCTTACCCCG | chr5:34,270,637-34,270,656 (+) | 10/25 (53%) | GAGTGGGGCTCTTTAGTCTTCTG | TGATCTTCCCCTCTAGGAGTACAG |
| *Polq* | Polq-JD3.1 | TAATTGAAATGGGAGTACGG | chr16:36,880,497-36,880,516 (+) | 16/30 (35%) | GGTGACTGGCAAGGGCTTAT | TGGGGACGGCTTGTCATTAG |
| *Rad18* | Rad18-JD3.2 | TCAGTGCCCAACTTGTTGCG | chr6:112,664,940-112,664,959 (-) | 3/12 (12%) | TTGTGTCTTTTTGCAGATTGTTCAC | ACCCATGTCTCTTGTGCATACTT |
| *Rnaseh1* | Rnaseh1-JD3.1 | TGGGCGACAGACAAACCAGA | chr12:28,707,292-28,707,311 (+) | 5/8 (63%) | CCTTTGAGAAGAGTTGCTTTGGG | GGCAGGAAAGGAAGGGGAAA |
| *Rnaseh2a* | Rnaseh2a-JD3.1 | TGTCACACGATACAGCTGCG | chr8:85,691,805-85,691,824 (-) | 4/8 (40%) | TGAGGCGTCCCTTCATGAGT | AGAAGGTGACCAAAGGGCTG |
| *Rrm2b* | Rrm2b-JD2.1 | GAGAATGTACAAGCAAGCAC | chr15:37,953,869-37,953,888 (-) | 2/9 (17%) | CAATTGTGTTCGGAGACAGAAGAA | TTCTGTAAGGCAAAGGTGAGTG |
| *Rrm2b* | Rrm2b-JD3.1 | GGCATCTTTCTGGACAGCAG | chr15:37,953,848-37,953,867 (-) | 2/9 (19%) | CAATTGTGTTCGGAGACAGAAGAA | TTCTGTAAGGCAAAGGTGAGTG |
| *Setd2* | Setd2-JD3.1 | GCATTCGCTTAATATCCCGG | chr9:110,382,189-110,382,208 (-) | 4/21 (58%) | CCACCTTAGTGTTGTCTTTTGAGT | ACCATTCAATCATGAGGAGACGA |
| *Setdb1* | Setdb1-JD3.1 | AGGACTAAGACATGGCACAA | chr3:95,254,336-95,254,355 (-) | 6/22 (16%) | AGCTACAGCCCATCCTACCT | GTCCTTCCTCCAGCCTGAAC |
| *Setx* | Setx-JD3.2 | AAAGGCTTGCTACTGTCATG | chr2:29,036,322-29,036,341 (-) | 10/26 (35%) | CTCCATGTGAGGGTCATGTAAAGA | GCAGCACTGTTTTGGTTAGCTG |
| *Sirt6* | Sirt6-JD3.1 | CATGCGTTCTAGCTGAACCA | chr10:81,461,615-81,461,634 (+) | 3/8 (29%) | CTATCCTCAGAGGCCCCCAT | GGCACCCTGGTCCAATAAGG |
| *Slx4* | Slx4-JD3.1 | GACTGTGTGAGGTCAATGGG | chr16:3,804,012-3,804,031 (+) | 11/15 (59%) | CAACTAAGCAGAGGAGGGGC | ACTCTGTCTTGGTGTGCTCC |
| *Smarcad1* | Smarcad1-JD3.1 | AACAGAGCACATTTAAACTG | chr6:65,071,149-65,071,168 (-) | 13/24 (55%) | ATCATGAGGGGTAGTTTTCATAGTG | TGAAGCTTACTTTGATTATATTTATCG |
| *Supt4a* | Supt4a-JD3.1 | ACCTGGCTTAAAGTTACCTG | chr11:87,633,881-87,633,900 (-) | 4/5 (66%) | GCTTAATTCTTTCTTGCTTTTCAGT | CCCGCACGATTCCTATGACA |
| *Tcea2* | Tcea2-JD3.1 | AGTTCACACCAACGGCCACG | chr2:181,327,990-181,328,009 (-) | 6/9 (52%) | GTGTAAACAGAGGGGTGGCA | GGAAAGGTCAAGGGACCACC |
| *Tcerg1* | Tcerg1-JD3.1 | TAGAACACTAGAATCCACGT | chr18:42,669,394-42,669,413 (+) | 7/22 (42%) | GGCCCCTCCTATAGTACCCAT | TATGTCCTTAGTCACAAAACAACTC |
| *Tdp1* | Tdp1-JD3.1 | ACGCCGTTAGATAACTGGTG | chr12:99,876,022-99,876,041 (-) | 8/16 (53%) | GGTTGAGCCCTTTGTACCCA | CATGCAGCACAGCCGAATAC |
| *Ubr5* | Ubr5-JD3.1 | TGTGGTAAGAGATCCCACAG | chr15:38,009,409-38,009,428 (+) | 24/59 (37%) | GAAAATGGTTAGGGGTATGTGCC | TTAATTCAAAAAGCCCAAGTAGACA |
| *Ung* | Ung-JD3.2 | TGAAGAAAGGAACCACCACA | chr5:114,270,259-114,270,278 (+) | 2/6 (36%) | TAAATCATGCGCGCACACAC | TTCCTCTCTAGCCAGACCCC |
| *Wrn* | Wrn-JD3.1 | TGTGTCCGAAATAGGTCTTG | chr8:33,809,193-33,809,212 (+) | 12/34 (36%) | AACATTTTCCTCTTCTTGCATAGC | ACAGAGTGATGGGTGAAACCG |
| *Xpa* | Xpa-JD3.1 | AAAATGATTGACACCAAAGG | chr4:46,193,112-46,193,131 (-) | 2/6 (26%) | TTTCCCCGCAGGAGAGTCTA | ACCAATGTCCATACTTCTTTTGTGG |

***Supplementary Table 2: In vivo CRISPR editing and CAG instability analysis following AAV8 treatment***

| Gene | sgRNA ID | Liver | | | | | | | | | | Hepatocytes | | | | | | | | |
|---|---|---|---|---|---|---|---|---|---|---|---|---|---|---|---|---|---|---|---|---|
| | | CAG instability analysis | | | | | | CRISPR editing quantification by NGS | | | | CAG instability analysis | | | | | CRISPR editing quantification by NGS | | | |
| | | Index (+ve) | z-score | ANOVA | p-value (vs Empty) | Number of animals | CAG | Frameshift (%) | Non-frameshift (%) | Unedited (%) | Number of animals | Index (+ve) | ANOVA | p-value (vs Empty) | Number of animals | CAG | Frameshift (%) | Non-frameshift (%) | Unedited (%) | Number of animals |
| *Atm* | Atm-JD3.1 | 4.255 ±0.324 | 0.310 ±0.334 | ns | 0.9995 | 4 (4M) | 117.3 ±1.50 | 52.4% ±6.3 | 1.2% ±0.8 | 46.4% ±6.7 | 4 (4M) | | | | | | | | | |
| *Brip1* | Brip1-JD2.6 | 4.220 ±0.409 | 0.274 ±0.421 | ns | 0.9996 | 7 (7M) | 116.1 ±2.54 | 65.1% ±1.3 | 7.4% ±0.2 | 27.5% ±1.2 | 2 (2M) | | | | | | | | | |
| *Ccdc82* | Ccdc82-JD3.1 | 4.157 ±0.343 | 0.209 ±0.353 | ns | >0.9999 | 12 (5M, 7F) | 116.2 ±1.34 | 66.1% ±8.4 | 1.4% ±0.4 | 32.5% ±8.7 | 12 (5M, 7F) | 5.019 ±0.876 | ns | 0.9986 | 3 (2M, 1F) | 115.7 ±1.53 | 88.5% ±1.8 | 2.1% ±0.5 | 9.4% ±2.3 | 3 (2M, 1F) |
| *Crebbp* | Crebbp-JD3.2 | 3.532 ±0.239 | -0.435 ±0.246 | ** | 0.0015 | 6 (3M, 3F) | 116.2 ±1.17 | 54.6% ±2.0 | 10.9% ±0.8 | 34.6% ±2.8 | 3 (1M, 2F) | 4.269 ±0.246 | ns | 0.8969 | 3 (1M, 2F) | 117.0 ±1.00 | 68.1% ±2.2 | 12.8% ±0.5 | 19.2% ±2.4 | 3 (1M, 2F) |
| *Dhfr* | Dhfr-JD3.1 | 4.179 ±0.286 | 0.232 ±0.294 | ns | 0.9999 | 6 (6M) | 118.2 ±0.98 | 60.1% ±3.5 | 5.1% ±1.4 | 34.9% ±4.5 | 6 (6M) | | | | | | | | | |
| *Dnmt1* | Dnmt1-JD3.1 | 4.354 ±0.321 | 0.412 ±0.331 | ns | 0.9838 | 10 (10M) | 116.8 ±2.30 | 61.1% ±3.0 | 3.9% ±0.6 | 34.9% ±2.5 | 3 (3M) | 5.055 ±0.089 | ns | 0.9983 | 3 (3M) | 118.7 ±0.58 | 83.7% ±5.5 | 4.7% ±0.4 | 11.5% ±5.1 | 3 (3M) |
| *Dnmt3a* | Dnmt3a-JD3.1 | 4.075 ±0.207 | 0.125 ±0.214 | ns | 0.9994 | 9 (9M) | 117.2 ±1.64 | 71.6% ±1.3 | 2.4% ±0.2 | 26.0% ±1.5 | 2 (2M) | 5.079 ±0.171 | ns | 0.9980 | 3 (3M) | 116.3 ±1.15 | 88.0% ±1.5 | 2.0% ±0.1 | 10.0% ±1.7 | 2 (2M) |
| Empty | | 4.159 ±0.294 | 0.211 ±0.303 | N/A | N/A | 30 (21M, 9F) | 115.3 ±2.44 | | | | | 4.729 ±0.304 | N/A | N/A | 16 (10M, 6F) | 115.3 ±2.89 | | | | |
| *Ercc1* | Ercc1-JD3.1 | 3.594 ±0.228 | -0.371 ±0.235 | ** | 0.0012 | 8 (5M, 3F) | 114.1 ±0.64 | 43.3% ±7.7 | 4.3% ±2.0 | 52.4% ±9.1 | 10 (6M, 4F) | 4.383 ±0.324 | ns | 0.9981 | 3 (2M, 1F) | 114.0 ±0.00 | 67.8% ±13.1 | 5.7% ±3.6 | 26.5% ±16.2 | 3 (2M, 1F) |
| *Ercc3* | Ercc3-JD2.1 | 3.730 ±0.260 | -0.231 ±0.268 | * | 0.0219 | 10 (10M) | 117.2 ±1.55 | 49.8% ±4.4 | 8.4% ±1.9 | 41.8% ±5.6 | 10 (10M) | | | | | | | | | |
| *Ercc5* | Ercc5-JD3.2 | 3.615 ±0.363 | -0.349 ±0.374 | ** | 0.0023 | 8 (4M, 4F) | 114.8 ±2.25 | 27.3% ±10.1 | 1.6% ±0.6 | 71.1% ±9.9 | 10 (6M, 4F) | 4.458 ±0.198 | ns | 0.9987 | 3 (2M, 1F) | 113.7 ±0.58 | 39.1% ±1.8 | 1.4% ±0.2 | 59.4% ±1.8 | 3 (2M, 1F) |
| *Exo1* | Exo1-JD2.1 | 4.251 ±0.362 | 0.305 ±0.373 | ns | 0.9994 | 6 (6M) | 117.3 ±1.37 | 45.5% ±6.1 | 9.0% ±1.6 | 45.5% ±7.3 | 8 (8M) | | | | | | | | | |
| *Fan1* | Fan1-JD2.1 | 7.442 ±0.530 | 3.593 ±0.546 | **** | <0.0001 | 16 (14M, 2F) | 115.7 ±1.45 | 73.7% ±5.4 | 1.9% ±0.5 | 24.3% ±5.4 | 13 (10M, 3F) | 8.401 ±1.249 | **** | <0.0001 | 3 (1M, 2F) | 115.0 ±1.00 | 83.8% ±6.7 | 1.5% ±1.0 | 14.8% ±7.2 | 4 (1M, 3F) |
| *Fen1* | Fen1-JD3.1 | 3.893 ±0.174 | -0.062 ±0.179 | ns | 0.9644 | 6 (6M) | 116.3 ±2.34 | 50.7% ±3.6 | 3.9% ±0.8 | 45.4% ±4.1 | 5 (5M) | 4.595 ±0.169 | ns | 0.9995 | 3 (3M) | 117.0 ±1.73 | 0.0% ±0.0 | 0.0% ±0.0 | 0.0% ±0.0 | (F) |
| *Gpr151* | Gpr151-JD3.1 | 4.275 ±0.325 | 0.331 ±0.335 | ns | 0.9989 | 12 (6M, 6F) | 115.7 ±1.87 | 67.6% ±2.6 | 0.6% ±0.5 | 31.8% ±2.7 | 12 (6M, 6F) | 4.651 ±0.355 | ns | 0.9997 | 3 (2M, 1F) | 115.7 ±2.52 | 87.6% ±4.4 | 0.4% ±0.5 | 11.9% ±4.4 | 3 (2M, 1F) |
| *Hdac6* | Hdac6-JD3.2 | 4.141 ±0.359 | 0.192 ±0.370 | ns | >0.9999 | 6 (3M, 3F) | 115.3 ±1.37 | 68.7% ±1.1 | 2.3% ±0.5 | 29.0% ±0.9 | 3 (1M, 2F) | 4.792 ±0.506 | ns | 0.9997 | 4 (1M, 3F) | 115.0 ±1.41 | 84.2% ±6.2 | 3.4% ±0.9 | 12.4% ±7.0 | 3 (1M, 2F) |
| *Hmgb1* | Hmgb1-JD3.2 | 5.151 ±0.433 | 1.233 ±0.446 | **** | <0.0001 | 12 (6M, 6F) | 116.5 ±1.88 | 66.6% ±2.9 | 1.7% ±0.7 | 31.7% ±2.8 | 12 (6M, 6F) | 6.121 ±0.331 | **** | <0.0001 | 3 (1M, 2F) | 116.7 ±2.52 | 86.0% ±2.1 | 1.8% ±0.3 | 12.2% ±1.9 | 3 (1M, 2F) |
| *Htt* | Htt-ex6.1 | 4.061 ±0.208 | 0.110 ±0.214 | ns | 0.9994 | 6 (3M, 3F) | 115.8 ±1.47 | 67.8% ±2.5 | 2.0% ±0.3 | 30.2% ±2.5 | 6 (3M, 3F) | 4.776 ±0.223 | ns | 0.9999 | 3 (2M, 1F) | 115.3 ±0.58 | 87.5% ±1.4 | 2.4% ±0.3 | 10.1% ±1.5 | 3 (2M, 1F) |
| *LacZ* | LacZ-JD3.1 | 4.293 ±0.294 | 0.349 ±0.303 | ns | 0.9986 | 12 (5M, 7F) | 115.6 ±2.35 | | | | | 5.054 ±0.308 | ns | 0.8705 | 8 (4M, 4F) | 115.8 ±2.19 | | | | |
| *Lig1* | Lig1-JD3.2 | 4.277 ±0.276 | 0.333 ±0.285 | ns | 0.9990 | 10 (8M, 2F) | 115.9 ±1.52 | 45.1% ±4.5 | 8.5% ±1.5 | 46.4% ±5.6 | 11 (9M, 2F) | 4.990 ±0.128 | ns | 0.9988 | 3 (1M, 2F) | 116.3 ±0.58 | 75.3% ±2.7 | 12.4% ±1.2 | 12.3% ±3.6 | 3 (1M, 2F) |
| *Lig3* | Lig3-JD3.1 | 3.955 ±0.264 | 0.001 ±0.272 | ns | 0.9984 | 5 (5M) | 117.8 ±0.84 | 65.4% ±2.1 | 0.4% ±0.4 | 34.2% ±2.3 | 6 (6M) | 4.532 ±0.361 | ns | 0.9990 | 5 (5M) | 117.8 ±0.84 | 91.8% ±3.1 | 0.6% ±0.4 | 7.6% ±3.2 | 5 (5M) |
| *Lig4* | Lig4-JD3.1 | 4.919 ±0.109 | 0.994 ±0.112 | *** | 0.0001 | 5 (3M, 2F) | 116.8 ±1.64 | 57.3% ±3.3 | 2.6% ±0.7 | 40.1% ±3.9 | 6 (3M, 3F) | 5.891 ±0.482 | ** | 0.0043 | 2 (2M) | 116.5 ±2.12 | 82.9% ±1.1 | 5.1% ±0.8 | 12.0% ±1.1 | 3 (2M, 1F) |
| *Mcm9* | Mcm9-JD3.1 | 4.537 ±0.342 | 0.601 ±0.352 | ns | 0.1290 | 9 (9M) | 118.4 ±0.73 | 69.2% ±0.8 | 5.2% ±0.7 | 25.6% ±1.2 | 4 (4M) | 4.693 ±0.486 | ns | 0.9999 | 4 (4M) | 118.5 ±0.58 | 86.5% ±1.1 | 8.8% ±0.5 | 4.7% ±1.2 | 4 (4M) |
| *Mlh1* | Mlh1-JD4 | 2.160 ±0.141 | -1.848 ±0.145 | **** | <0.0001 | 11 (8M, 3F) | 115.6 ±1.91 | 74.6% ±4.5 | 0.3% ±0.5 | 25.1% ±4.3 | 10 (7M, 3F) | | | | | | | | | |
| *Mlh3* | Mlh3-JD2 | 1.864 ±0.223 | -2.152 ±0.229 | **** | <0.0001 | 7 (7M) | 115.4 ±2.23 | 51.6% ±3.5 | 13.3% ±1.6 | 35.0% ±2.4 | 7 (5M, 2F) | 1.816 | **** | <0.0001 | 1 (1M) | 114 | 64.3% ±1.4 | 20.2% ±1.2 | 15.5% ±2.6 | 4 (2M, 2F) |
| *Msh2* | Msh2-JD1 | 1.857 ±0.159 | -2.160 ±0.164 | **** | <0.0001 | 7 (7M) | 115.9 ±1.95 | 57.1% ±6.9 | 12.8% ±3.3 | 30.1% ±4.8 | 3 (3M) | | | | | | | | | |
| *Msh3* | Msh3-JD2.3 | 1.797 ±0.183 | -2.221 ±0.189 | **** | <0.0001 | 12 (9M, 3F) | 117.0 ±1.65 | 75.9% ±1.9 | 3.1% ±0.7 | 21.0% ±2.1 | 11 (8M, 3F) | 2.077 ±0.238 | **** | <0.0001 | 5 (2M, 3F) | 117.4 ±1.14 | 90.9% ±3.3 | 2.0% ±0.5 | 7.1% ±3.5 | 5 (2M, 3F) |
| *Msh4* | Msh4-JD3.1 | 4.122 ±0.180 | 0.173 ±0.185 | ns | 0.9998 | 6 (6M) | 116.7 ±2.07 | 63.7% ±5.2 | 1.8% ±0.5 | 34.5% ±5.0 | 6 (6M) | | | | | | | | | |
| *Msh5* | Msh5-JD3.1 | 4.032 ±0.325 | 0.080 ±0.335 | ns | 0.9990 | 8 (7M, 1F) | 117.0 ±2.00 | 78.9% ±1.9 | 0.8% ±0.2 | 20.3% ±2.0 | 5 (5M) | | | | | | | | | |
| *Msh6* | Msh6-JD2.2 | 4.805 ±0.413 | 0.876 ±0.426 | **** | <0.0001 | 16 (16M) | 117.2 ±1.87 | 54.0% ±4.8 | 11.1% ±5.3 | 34.9% ±2.7 | 16 (14M, 2F) | 5.119 ±0.715 | ns | 0.4722 | 9 (9M) | 118.1 ±0.93 | 69.1% ±6.0 | 12.7% ±6.0 | 18.2% ±9.9 | 9 (7M, 2F) |
| *Ogg1* | Ogg1-JD3.1 | 4.029 ±0.328 | 0.077 ±0.338 | ns | 0.9991 | 6 (6M) | 116.0 ±2.37 | 59.1% ±9.3 | 3.1% ±0.7 | 37.8% ±9.9 | 6 (6M) | 4.737 ±0.131 | ns | >0.9999 | 3 (3M) | 117.0 ±2.65 | 0.0% ±0.0 | 0.0% ±0.0 | 0.0% ±0.0 | (F) |
| *Parp1* | Parp1-JD3.1 | 4.199 ±0.358 | 0.253 ±0.369 | ns | 0.9997 | 6 (3M, 3F) | 116.2 ±2.32 | 62.3% ±2.6 | 2.5% ±0.5 | 35.2% ±3.0 | 3 (1M, 2F) | 4.676 ±0.438 | ns | 0.9998 | 3 (1M, 2F) | 116.7 ±1.53 | 75.5% ±13.7 | 1.1% ±1.1 | 23.4% ±14.1 | 3 (1M, 2F) |
| *Pcna* | Pcna-JD3.1 | 3.339 ±0.374 | -0.634 ±0.385 | **** | <0.0001 | 17 (17M) | 117.6 ±1.46 | 19.0% ±4.1 | 2.0% ±2.5 | 79.0% ±5.9 | 13 (13M) | 4.554 ±0.215 | ns | 0.9990 | 6 (6M) | 118.5 ±0.55 | 45.4% ±8.1 | 15.7% ±10.1 | 39.0% ±18.1 | 9 (9M) |
| *Pms1* | Pms1-JD2.2 | 2.155 ±0.074 | -1.853 ±0.076 | **** | <0.0001 | 6 (6M) | 116.3 ±2.34 | 59.2% ±2.6 | 9.3% ±2.0 | 31.6% ±4.1 | 7 (7M) | | | | | | | | | |
| *Pms2* | Pms2-JD2.2 | 5.358 ±0.590 | 1.446 ±0.608 | **** | <0.0001 | 12 (10M, 2F) | 117.1 ±1.88 | 66.8% ±3.2 | 1.7% ±0.8 | 31.5% ±3.5 | 15 (11M, 4F) | 6.760 ±0.392 | **** | <0.0001 | 2 (1M, 1F) | 118.0 ±0.00 | 82.7% ±6.6 | 1.9% ±0.8 | 15.4% ±6.4 | 8 (4M, 4F) |
| *Polb* | Polb-JD3.1 | 3.532 ±0.172 | -0.435 ±0.178 | *** | 0.0004 | 7 (7M) | 116.9 ±1.95 | 58.7% ±4.2 | 2.8% ±0.6 | 38.4% ±4.6 | 6 (6M) | | | | | | | | | |
| *Pold1* | Pold1-JD3.1 | 2.133 ±0.147 | -1.876 ±0.152 | **** | <0.0001 | 6 (3M, 3F) | 118.2 ±1.17 | 36.1% ±8.0 | 1.5% ±1.7 | 62.4% ±8.6 | 7 (4M, 3F) | 2.874 ±0.315 | **** | <0.0001 | 6 (3M, 3F) | 118.2 ±1.17 | 57.3% ±16.1 | 3.0% ±2.5 | 39.7% ±15.8 | 7 (4M, 3F) |
| *Pold2* | Pold2-JD3.1 | 2.744 ±0.298 | -1.246 ±0.306 | **** | <0.0001 | 12 (6M, 6F) | 116.0 ±1.81 | 40.2% ±3.2 | 7.3% ±2.6 | 52.5% ±4.1 | 12 (6M, 6F) | 3.043 ±0.065 | **** | <0.0001 | 3 (2M, 1F) | 116.7 ±2.31 | 60.3% ±9.0 | 10.5% ±0.5 | 29.1% ±8.6 | 3 (2M, 1F) |
| *Pold3* | Pold3-JD3.1 | 2.186 ±0.182 | -1.821 ±0.187 | **** | <0.0001 | 6 (4M, 2F) | 118.7 ±0.52 | 38.0% ±17.6 | 1.8% ±1.9 | 60.2% ±18.5 | 6 (4M, 2F) | 2.863 ±0.105 | **** | <0.0001 | 6 (4M, 2F) | 118.7 ±0.52 | 64.4% ±15.7 | 3.9% ±2.4 | 31.7% ±16.9 | 6 (4M, 2F) |
| *Pold4* | Pold4-JD3.2 | 3.344 ±0.334 | -0.628 ±0.344 | **** | <0.0001 | 12 (6M, 6F) | 115.8 ±2.05 | 61.2% ±3.7 | 1.0% ±0.5 | 37.8% ±3.5 | 12 (6M, 6F) | 3.986 ±0.424 | ns | 0.1063 | 3 (2M, 1F) | 117.3 ±2.08 | 84.3% ±1.7 | 1.5% ±0.1 | 14.2% ±1.6 | 3 (2M, 1F) |
| *Pole* | Pole-JD3.1 | 3.626 ±0.430 | -0.338 ±0.443 | *** | 0.0002 | 12 (6M, 6F) | 116.2 ±2.25 | 38.0% ±16.8 | 2.9% ±1.3 | 59.1% ±16.2 | 12 (6M, 6F) | 4.288 ±0.212 | ns | 0.9341 | 3 (1M, 2F) | 115.0 ±1.73 | 68.0% ±1.4 | 3.1% ±0.4 | 28.8% ±1.8 | 3 (1M, 2F) |
| *Polh* | Polh-JD3.1 | 4.391 ±0.342 | 0.450 ±0.352 | ns | 0.9178 | 10 (10M) | 118.6 ±0.52 | 66.7% ±0.7 | 0.0% ±0.0 | 33.3% ±0.7 | 3 (3M) | 5.055 ±0.642 | ns | 0.9860 | 4 (4M) | 118.5 ±0.58 | 95.1% ±0.7 | 0.0% ±0.0 | 4.9% ±0.7 | 3 (3M) |
| *Poln* | Poln-JD3.1 | 3.973 ±0.405 | 0.020 ±0.418 | ns | 0.9847 | 11 (5M, 6F) | 116.2 ±1.33 | 59.7% ±3.6 | 0.5% ±0.6 | 39.8% ±3.9 | 11 (5M, 6F) | 4.685 ±0.263 | ns | 0.9999 | 3 (2M, 1F) | 115.7 ±2.52 | 81.1% ±2.4 | 0.8% ±0.2 | 18.1% ±2.5 | 3 (2M, 1F) |
| *Polq* | Polq-JD3.1 | 4.370 ±0.157 | 0.429 ±0.162 | ns | 0.9981 | 6 (3M, 3F) | 114.8 ±2.56 | 50.1% ±1.8 | 3.2% ±0.6 | 46.7% ±2.2 | 6 (3M, 3F) | | | | | | | | | |
| *Rad18* | Rad18-JD3.2 | 4.355 ±0.278 | 0.413 ±0.286 | ns | 0.9985 | 5 (3M, 2F) | 115.4 ±1.14 | 59.2% ±9.6 | 3.6% ±1.2 | 37.3% ±10.8 | 3 (1M, 2F) | 5.071 ±0.436 | ns | 0.9981 | 3 (1M, 2F) | 115.0 ±1.00 | 83.1% ±4.9 | 5.2% ±0.4 | 11.8% ±4.6 | 3 (1M, 2F) |
| *Rnaseh1* | Rnaseh1-JD3.1 | 4.123 ±0.381 | 0.174 ±0.393 | ns | 0.9998 | 5 (5M) | 117.8 ±1.30 | 67.9% ±2.1 | 4.9% ±0.8 | 27.2% ±1.6 | 3 (3M) | 4.958 ±0.533 | ns | 0.9988 | 4 (4M) | 117.5 ±1.29 | 85.8% ±3.0 | 5.4% ±0.8 | 8.7% ±3.6 | 3 (3M) |
| *Rnaseh2a* | Rnaseh2a-JD3.1 | 4.062 ±0.318 | 0.111 ±0.327 | ns | 0.9994 | 6 (6M) | 117.5 ±1.05 | 69.0% ±0.9 | 0.7% ±0.2 | 30.3% ±0.9 | 4 (4M) | 4.744 ±0.534 | ns | >0.9999 | 5 (5M) | 117.6 ±1.14 | 84.5% ±0.0 | 0.6% ±0.0 | 14.9% ±0.0 | 1 (1M) |
| *Rrm2b* | Rrm2b-JD2.1 | 3.794 ±0.474 | -0.165 ±0.489 | ns | 0.4731 | 6 (6M) | 116.0 ±2.37 | 49.4% ±5.2 | 2.8% ±1.4 | 47.8% ±6.0 | 7 (7M) | | | | | | | | | |
| *Rrm2b* | Rrm2b-JD3.1 | | | | | | | | | | | 3.939 ±0.334 | ns | 0.0604 | 3 (3M) | 117.3 ±1.53 | 69.0% ±8.9 | 15.1% ±1.0 | 15.9% ±9.9 | 3 (3M) |
| *Setd2* | Setd2-JD3.1 | 3.704 ±0.183 | -0.257 ±0.189 | * | 0.0174 | 9 (9M) | 118.3 ±1.32 | 51.3% ±5.2 | 1.0% ±0.9 | 47.8% ±6.0 | 3 (3M) | 4.546 ±0.158 | ns | 0.9990 | 5 (5M) | 118.6 ±0.55 | 87.9% ±0.3 | 3.2% ±0.8 | 8.9% ±1.0 | 3 (3M) |
| *Setdb1* | Setdb1-JD3.1 | 3.722 ±0.183 | -0.239 ±0.189 | * | 0.0492 | 8 (8M) | 117.9 ±0.99 | 57.2% ±3.6 | 2.1% ±0.4 | 40.7% ±3.5 | 8 (8M) | 4.421 ±0.238 | ns | 0.9844 | 5 (5M) | 118.4 ±0.89 | 91.3% ±0.2 | 3.4% ±0.4 | 5.1% ±0.7 | 5 (5M) |
| *Setx* | Setx-JD3.2 | 4.056 ±0.377 | 0.105 ±0.389 | ns | 0.9994 | 5 (3M, 2F) | 113.0 ±1.22 | 46.6% ±4.4 | 7.2% ±1.0 | 46.3% ±5.4 | 6 (3M, 3F) | 4.739 ±0.412 | ns | >0.9999 | 3 (1M, 2F) | 113.3 ±1.53 | 60.6% ±10.1 | 8.8% ±1.9 | 30.6% ±11.8 | 3 (1M, 2F) |
| *Sirt6* | Sirt6-JD3.1 | 4.084 ±0.502 | 0.134 ±0.517 | ns | 0.9996 | 6 (3M, 3F) | 116.2 ±1.60 | 63.7% ±4.0 | 3.0% ±0.4 | 33.2% ±3.7 | 3 (1M, 2F) | 4.566 ±0.154 | ns | 0.9994 | 3 (1M, 2F) | 115.7 ±1.15 | 80.5% ±1.9 | 2.7% ±0.2 | 16.8% ±2.2 | 3 (1M, 2F) |
| *Slx4* | Slx4-JD3.1 | 4.024 ±0.267 | 0.072 ±0.274 | ns | 0.9991 | 6 (3M, 3F) | 113.8 ±1.47 | 60.2% ±2.3 | 1.6% ±0.6 | 38.2% ±2.3 | 6 (3M, 3F) | 4.522 ±0.245 | ns | 0.9991 | 3 (1M, 2F) | 113.3 ±1.53 | 80.8% ±2.2 | 3.5% ±0.3 | 15.7% ±2.4 | 3 (1M, 2F) |
| *Smarcad1* | Smarcad1-JD3.1 | 4.109 ±0.293 | 0.160 ±0.302 | ns | 0.9996 | 12 (5M, 7F) | 114.0 ±1.76 | 65.5% ±1.8 | 4.4% ±0.6 | 30.0% ±1.9 | 13 (5M, 8F) | 4.599 ±0.349 | ns | 0.9993 | 6 (2M, 4F) | 114.5 ±1.87 | 83.4% ±2.4 | 5.2% ±0.7 | 11.4% ±3.0 | 6 (2M, 4F) |
| *Supt4a* | Supt4a-JD3.1 | 4.215 ±0.426 | 0.268 ±0.439 | ns | 0.9996 | 12 (6M, 6F) | 113.6 ±1.24 | 42.7% ±3.7 | 3.4% ±2.3 | 53.9% ±4.0 | 13 (6M, 7F) | 4.662 ±0.229 | ns | 0.9997 | 5 (3M, 2F) | 113.8 ±1.10 | 68.7% ±7.4 | 9.2% ±4.0 | 22.1% ±8.1 | 6 (3M, 3F) |
| *Tcea2* | Tcea2-JD3.1 | 4.024 ±0.123 | 0.072 ±0.127 | ns | 0.9991 | 6 (3M, 3F) | 113.2 ±0.75 | 61.9% ±2.6 | 3.2% ±1.0 | 34.9% ±3.1 | 6 (3M, 3F) | 4.636 ±0.047 | ns | 0.9997 | 3 (1M, 2F) | 112.7 ±0.58 | 77.3% ±5.3 | 4.4% ±1.4 | 18.3% ±4.7 | 3 (1M, 2F) |
| *Tcerg1* | Tcerg1-JD3.1 | 4.237 ±0.387 | 0.291 ±0.398 | ns | 0.9993 | 14 (11M, 3F) | 117.1 ±1.73 | 56.9% ±3.2 | 7.6% ±1.2 | 35.4% ±4.3 | 8 (5M, 3F) | 5.058 ±0.241 | ns | 0.9784 | 5 (5M) | 118.0 ±0.71 | 81.9% ±0.1 | 11.4% ±0.9 | 6.7% ±1.0 | 3 (3M) |
| *Tdp1* | Tdp1-JD3.1 | 4.249 ±0.277 | 0.304 ±0.286 | ns | 0.9994 | 6 (4M, 2F) | 114.3 ±1.51 | 60.5% ±3.1 | 5.2% ±0.3 | 34.3% ±3.3 | 6 (4M, 2F) | 4.770 ±0.199 | ns | 0.9999 | 2 (2M) | 115.0 ±2.83 | 80.7% ±2.9 | 7.2% ±0.5 | 12.0% ±3.1 | 3 (2M, 1F) |
| *Ubr5* | Ubr5-JD3.1 | 3.879 ±0.308 | -0.077 ±0.317 | ns | 0.9289 | 6 (6M) | 118.2 ±1.17 | 54.4% ±3.4 | 0.0% ±0.0 | 45.6% ±3.4 | 6 (6M) | 4.670 ±0.421 | ns | 0.9997 | 6 (6M) | 118.2 ±1.17 | 93.8% ±1.6 | 0.4% ±0.9 | 5.8% ±1.2 | 5 (5M) |
| Untreated | | 3.799 ±0.320 | -0.160 ±0.329 | **** | <0.0001 | 77 (56M, 21F) | 116.0 ±2.06 | | | | | 4.352 ±0.327 | ns | 0.1253 | 23 (18M, 5F) | 116.1 ±2.11 | | | | |
| *Ung* | Ung-JD3.2 | 3.660 ±0.265 | -0.303 ±0.273 | ns | 0.0921 | 5 (3M, 2F) | 113.8 ±0.45 | 71.4% ±2.7 | 3.4% ±0.5 | 25.2% ±2.9 | 10 (6M, 4F) | 4.312 ±0.269 | ns | 0.9663 | 3 (2M, 1F) | 113.7 ±0.58 | 85.8% ±2.9 | 3.0% ±0.4 | 11.1% ±3.2 | 3 (2M, 1F) |
| *Wrn* | Wrn-JD3.1 | 3.897 ±0.249 | -0.059 ±0.256 | ns | 0.9979 | 4 (4M) | 116.3 ±2.22 | 50.1% ±1.1 | 3.3% ±0.5 | 46.6% ±1.5 | 4 (4M) | | | | | | | | | |
| *Xpa* | Xpa-JD3.1 | 4.477 ±0.291 | 0.538 ±0.299 | ns | 0.2713 | 11 (11M) | 117.7 ±1.27 | 54.5% ±5.7 | 0.1% ±0.2 | 45.4% ±5.8 | 6 (6M) | 4.769 ±0.180 | ns | 0.9999 | 2 (2M) | 118.5 ±0.71 | 78.7% ±0.2 | 0.9% ±0.2 | 20.3% ±0.4 | 2 (2M) |

***Supplementary Table 3: List of genes tested with AAV8 delivery in liver***

| Gene | Role in CAG expansions | p-value † (Liver) | Sig † | Rationale for testing<br>*Note that references to known modifier effects are not exhaustive but represent most relevant data providing rationale for the current study. For additional references see Wheeler&Dion 2021, Journal of Huntington's Disease 10, 123–148.* | References |
|---|---|---|---|---|---|
| *Atm* | | 0.9995 | *ns* | DNA damage response gene. Modifies huntingtin toxicity (Lu et al. 2014). | Lu, X.-H. et al. Targeting ATM ameliorates mutant Huntingtin toxicity in cell and animal models of Huntington's disease. Sci. Transl. Med. 6, 268ra178 (2014). |
| *Brip1* | | 0.9996 | *ns* | Encodes FANCJ, a DNA helicase that maintains the integrity/stability of microsatellite repeats including CAG/CTG (Matsuzaki et al. 2015, Barthelemy et al. 2016). | Matsuzaki, K., Borel, V., Adelman, C. A., Schindler, D. & Boulton, S. J. FANCJ suppresses microsatellite instability and lymphomagenesis independent of the Fanconi anemia pathway. Genes Dev. 29, 2532–2546 (2015).; Barthelemy, J., Hanenberg, H. & Leffak, M. FANCJ is essential to maintain microsatellite structure genome-wide during replication stress. Nucleic Acids Res. 44, 6803–6816 (2016). |
| *Ccdc82* | | >0.9999 | *ns* | Candidate genetic modifier of HD age of onset (GeM-HD Consortium 2019). | Genetic Modifiers of Huntington's Disease (GeM-HD) Consortium. Identification of Genetic Factors that Modify Clinical Onset of Huntington's Disease. Cell 162, 516–526 (2015). |
| *Crebbp* | Enhancer | 0.0015 | ** | Encodes CBP, a H3K56 acetyltansferase. H3K56 acetylation is imporant in genome stability (Vempati et al. 2010). Modifies CAG/CTG instability in human cells (Debacker et al. 2012), in *Drosophila* (Jung&Bonini et al. 2007) and exhibits structural similarity to Rtt109 that modifies CAG/CTG instability in yeast (Yang&Freudenreich 2010). | Vempati, R. K. et al. p300-mediated acetylation of histone H3 lysine 56 functions in DNA damage response in mammals. J. Biol. Chem. 285, 28553–28564 (2010).; Debacker, K. et al. Histone deacetylase complexes promote trinucleotide repeat expansions. PLoS Biol. 10, e1001257 (2012).; Jung, J. & Bonini, N. CREB-binding protein modulates repeat instability in a Drosophila model for polyQ disease. Science 315, 1857–1859 (2007).; Yang, J. H. & Freudenreich, C. H. The Rtt109 histone acetyltransferase facilitates error-free replication to prevent CAG/CTG repeat contractions. DNA Repair (Amst) 9, 414–420 (2010). |
| *Dhfr* | | 0.9999 | *ns* | Candidate genetic modifier of HD age of onset (GeM-HD Consortium 2019). | Genetic Modifiers of Huntington's Disease (GeM-HD) Consortium. Identification of Genetic Factors that Modify Clinical Onset of Huntington's Disease. Cell 162, 516–526 (2015). |
| *Dnmt1* | | 0.9838 | *ns* | DNA demetylation destabilizes CAG/CTG repeats (Gorbunova et al. 2004, Dion et al. 2008). | Gorbunova, V., Seluanov, A., Mittelman, D. & Wilson, J. H. Genome-wide demethylation destabilizes CTG.CAG trinucleotide repeats in mammalian cells. Hum. Mol. Genet. 13, 2979–2989 (2004).; Dion, V., Lin, Y., Hubert, L., Waterland, R. A. & Wilson, J. H. Dnmt1 deficiency promotes CAG repeat expansion in the mouse germline. Hum. Mol. Genet. 17, 1306–1317 (2008). |
| *Dnmt3a* | | 0.9994 | *ns* | DNA demetylation destabilizes CAG/CTG repeats (Gorbunova et al. 2004, Dion et al. 2008). | Gorbunova, V., Seluanov, A., Mittelman, D. & Wilson, J. H. Genome-wide demethylation destabilizes CTG.CAG trinucleotide repeats in mammalian cells. Hum. Mol. Genet. 13, 2979–2989 (2004).; Dion, V., Lin, Y., Hubert, L., Waterland, R. A. & Wilson, J. H. Dnmt1 deficiency promotes CAG repeat expansion in the mouse germline. Hum. Mol. Genet. 17, 1306–1317 (2008). |
| *Ercc1* | Enhancer | 0.0012 | ** | ERCC1/ERCC4 (XPF) is a structure-specific nuclease responsible for the 5'-incision during DNA repair. Acts with SLX4 in the repair of interstrand cross-links. Modifies CAG/CTG instabilty in human cells (Lin&Wilson 2007). | Lin, Y. & Wilson, J. H. Transcription-induced CAG repeat contraction in human cells is mediated in part by transcription-coupled nucleotide excision repair. Mol. Cell. Biol. 27, 6209–6217 (2007). |
| *Ercc3* | Enhancer | 0.0219 | * | ERCC3 (XPB) is a 3'-5' DNA helicase, component of the TFIIH core complex, involved in general and transcription-coupled nucleotide excision repair. | |
| *Ercc5* | Enhancer | 0.0023 | ** | ERCC5 (XPG) is a single-stranded structure-specific DNA endonuclease involved in DNA excision repair. Modifies CAG/CTG instabilty in human cells (Lin&Wilson 2007) and Drosophiia (Jung&Bonini et al. 2007). | Lin, Y. & Wilson, J. H. Transcription-induced CAG repeat contraction in human cells is mediated in part by transcription-coupled nucleotide excision repair. Mol. Cell. Biol. 27, 6209–6217 (2007).; Jung, J. & Bonini, N. CREB-binding protein modulates repeat instability in a Drosophila model for polyQ disease. Science 315, 1857–1859 (2007). |
| *Exo1* | | 0.9994 | *ns* | Exonuclease involved in mismatch repair. | |
| *Fan1* | Supressor | <0.0001 | **** | Candidate genetic modifier of HD age of onset (GeM-HD Consortium 2015, 2019). Modifier of somatic CAG expansion in *Htt* .Q111 mice (Loupe et al. 2020). | Genetic Modifiers of Huntington's Disease (GeM-HD) Consortium. CAG repeat not polyglutamine length determines timing of huntington's disease onset. Cell 178, 887-900.e14 (2019).; Genetic Modifiers of Huntington's Disease (GeM-HD) Consortium. Identification of Genetic Factors that Modify Clinical Onset of Huntington's Disease. Cell 162, 516–526 (2015).; Loupe, J. M. et al. Promotion of somatic CAG repeat expansion by Fan1 knock-out in Huntington's disease knock-in mice is blocked by Mlh1 knock-out. Hum. Mol. Genet. 29, 3044–3053 (2020). |
| *Fen1* | | 0.9644 | *ns* | Orthologue (*RAD27*) modifies CAG/CTG instability in yeast (Freudenreich et al. 1998, Schweitzer et al. 1998, Yang&Freudenreich 2010). Influences male germline CAG/CTG instability in mice (Spiro et al. 2003). Promotes CAG expansion in cell-free base exsision repair (Liu et al. 2009). | Freudenreich, C. H., Kantrow, S. M. & Zakian, V. A. Expansion and length-dependent fragility of CTG repeats in yeast. Science 279, 853–856 (1998).; Schweitzer, J. K. & Livingston, D. M. Expansions of CAG repeat tracts are frequent in a yeast mutant defective in Okazaki fragment maturation. Hum. Mol. Genet. 7, 69–74 (1998).; Yang, J. H. & Freudenreich, C. H. The Rtt109 histone acetyltransferase facilitates error-free replication to prevent CAG/CTG repeat contractions. DNA Repair (Amst) 9, 414–420 (2010).; Spiro, C. & McMurray, C. T. Nuclease-deficient FEN-1 blocks Rad51/BRCA1-mediated repair and causes trinucleotide repeat instability. Mol. Cell. Biol. 23, 6063–6074 (2003).; Liu, Y. et al. Coordination between polymerase beta and FEN1 can modulate CAG repeat expansion. J. Biol. Chem. 284, 28352–28366 (2009). |
| *Gpr151* | | 0.9989 | *ns* | Candidate genetic modifier of HD age of onset (GeM-HD Consortium 2019). | Genetic Modifiers of Huntington's Disease (GeM-HD) Consortium. Identification of Genetic Factors that Modify Clinical Onset of Huntington's Disease. Cell 162, 516–526 (2015). |
| *Hdac6* | | >0.9999 | *ns* | Regulates mismatch repair (Zhang et al. 2014, Zhang et al. 2019). | Zhang, M. et al. HDAC6 deacetylates and ubiquitinates MSH2 to maintain proper levels of MutSα. Mol. Cell 55, 31–46 (2014).; Zhang, M. et al. HDAC6 regulates DNA damage response via deacetylating MLH1. J. Biol. Chem. 294, 5813–5826 (2019). |
| *Hmgb1* | Supressor | <0.0001 | **** | Involved in mismatch repair (Yuan et al. 2004, Zhang et al. 2005, Genschel et al. 2009). Binds slipped repeat structures (Slean et al. 2013). Promotes CAG expansion in cell-free base exsision repair (Liu et al. 2009). | Yuan, F., Gu, L., Guo, S., Wang, C. & Li, G.-M. Evidence for involvement of HMGB1 protein in human DNA mismatch repair. J. Biol. Chem. 279, 20935–20940 (2004).; Zhang, Y. et al. Reconstitution of 5'-directed human mismatch repair in a purified system. Cell 122, 693–705 (2005).; Genschel, J. & Modrich, P. Functions of MutLalpha, replication protein A (RPA), and HMGB1 in 5'-directed mismatch repair. J. Biol. Chem. 284, 21536–21544 (2009).; Slean, M. M. et al. Interconverting conformations of slipped-DNA junctions formed by trinucleotide repeats affect repair outcome. Biochemistry 52, 773–785 (2013).; Liu, Y. et al. Coordination between polymerase beta and FEN1 can modulate CAG repeat expansion. J. Biol. Chem. 284, 28352–28366 (2009). |
| *Htt* | | 0.9994 | *ns* | Candidate genetic modifier of HD age of onset (GeM-HD Consortium 2019) | Genetic Modifiers of Huntington's Disease (GeM-HD) Consortium. Identification of Genetic Factors that Modify Clinical Onset of Huntington's Disease. Cell 162, 516–526 (2015). |
| *Lig1* | | 0.9990 | *ns* | DNA ligase involved in mismatch repair. Candidate genetic modifier of HD age of onset (GeM-HD Consortium 2019). Modifier of CAG/CTG instability in the maternal germline in mice (Tome et al. 2011). | Genetic Modifiers of Huntington's Disease (GeM-HD) Consortium. Identification of Genetic Factors that Modify Clinical Onset of Huntington's Disease. Cell 162, 516–526 (2015).; Tomé, S. et al. Maternal germline-specific effect of DNA ligase I on CTG/CAG instability. Hum. Mol. Genet. 20, 2131–2143 (2011). |
| *Lig3* | | 0.9984 | *ns* | DNA ligase involved in nucleotide excision and base excision repair. | |
| *Lig4* | Supressor | 0.0001 | *** | DNA ligase involved in non-homologous end-joining. Modifer of *Fmr1* CGG repeat instability (Gazy et al. 2019). | Gazy, I., Hayward, B., Potapova, S., Zhao, X. & Usdin, K. Double-strand break repair plays a role in repeat instability in a fragile X mouse model. DNA Repair (Amst) 74, 63–69 (2019). |
| *Mcm9* | | 0.1290 | *ns* | DNA helicase involved in mismatch repair (Traver et al. 2015). | Traver, S. et al. MCM9 is required for mammalian DNA mismatch repair. Mol. Cell 59, 831–839 (2015). |
| *Mlh1* | Enhancer | <0.0001 | **** | Modifier of somatic CAG expansion in *Htt.* Q111 mice (Mouro Pinto et al. 2013). Candidate genetic modifier of HD age of onset (GeM-HD Consortium 2015, 2019, Lee et al. 2017). | Mouro Pinto, R. et al. Mismatch repair genes Mlh1 and Mlh3 modify CAG instability in Huntington's disease mice: genome-wide and candidate approaches. PLoS Genet. 9, e1003930 (2013).; Genetic Modifiers of Huntington's Disease (GeM-HD) Consortium. CAG repeat not polyglutamine length determines timing of huntington's disease onset. Cell 178, 887-900.e14 (2019).; Genetic Modifiers of Huntington's Disease (GeM-HD) Consortium. Identification of Genetic Factors that Modify Clinical Onset of Huntington's Disease. Cell 162, 516–526 (2015).; Lee, J.-M. et al. A modifier of Huntington's disease onset at the MLH1 locus. Hum. Mol. Genet. 26, 3859–3867 (2017). |
| *Mlh3* | Enhancer | <0.0001 | **** | Modifier of somatic CAG expansion in *Htt* .Q111 mice and HD patient fibroblasts (Mouro Pinto et al. 2013, Roy et al. 2021). Modifier of somatic CAG expansion in HD patient blood (Ciosi et al. 2019). | Mouro Pinto, R. et al. Mismatch repair genes Mlh1 and Mlh3 modify CAG instability in Huntington's disease mice: genome-wide and candidate approaches. PLoS Genet. 9, e1003930 (2013).; Roy, J. C. L. et al. Somatic CAG expansion in Huntington's disease is dependent on the MLH3 endonuclease domain, which can be excluded via splice redirection. Nucleic Acids Res. 49, 3907–3918 (2021).; Ciosi, M. et al. A genetic association study of glutamine-encoding DNA sequence structures, somatic CAG expansion, and DNA repair gene variants, with Huntington disease clinical outcomes. EBioMedicine 48, 568–580 (2019). |
| *Msh2* | Enhancer | <0.0001 | **** | Modifier of somatic CAG expansion in *Htt.* Q111 mice (Manley et al. 2019, Wheeler et al. 2003, Kovalenko et al. 2012). | Manley, K., Shirley, T. L., Flaherty, L. & Messer, A. Msh2 deficiency prevents in vivo somatic instability of the CAG repeat in Huntington disease transgenic mice. Nat. Genet. 23, 471–473 (1999).; Wheeler, V. C. et al. Mismatch repair gene Msh2 modifies the timing of early disease in Hdh(Q111) striatum. Hum. Mol. Genet. 12, 273–281 (2003).; Kovalenko, M. et al. Msh2 acts in medium-spiny striatal neurons as an enhancer of CAG instability and mutant huntingtin phenotypes in Huntington's disease knock-in mice. PLoS ONE 7, e44273 (2012). |
| *Msh3* | Enhancer | <0.0001 | **** | Modifier of somatic CAG expansion in *Htt.* Q111 mice (Dragileva et al. 2009). Candidate genetic modifier of HD age of onset, progression and somatic CAG expansion in HD patient blood (GeM-HD Consortium 2019, Hensman Moss et al. 2017, Ciosi et al. 2019, Flower et al. 2019) | Dragileva, E. et al. Intergenerational and striatal CAG repeat instability in Huntington's disease knock-in mice involve different DNA repair genes. Neurobiol. Dis. 33, 37–47 (2009).; Genetic Modifiers of Huntington's Disease (GeM-HD) Consortium. Identification of Genetic Factors that Modify Clinical Onset of Huntington's Disease. Cell 162, 516–526 (2015).; Hensman Moss, D. J. et al. Identification of genetic variants associated with Huntington's disease progression: a genome-wide association study. Lancet Neurol. 16, 701–711 (2017).; Ciosi, M. et al. A genetic association study of glutamine-encoding DNA sequence structures, somatic CAG expansion, and DNA repair gene variants, with Huntington disease clinical outcomes. EBioMedicine 48, 568–580 (2019).; Flower, M. et al. MSH3 modifies somatic instability and disease severity in Huntington's and myotonic dystrophy type 1. Brain 142, 1876–1886 (2019). |
| *Msh4* | | 0.9998 | *ns* | Member of the DNA mismatch repair MutS family that acts as a dimer with MSH5 in meiotic recombination. | |
| *Msh5* | | 0.9990 | *ns* | Member of the DNA mismatch repair MutS family that acts as a dimer with MSH4 in meiotic recombination. | |
| *Msh6* | Supressor | <0.0001 | **** | Variable effects on CAG/CTG instability (summarized in Wheeler&Dion 2021) | Wheeler, V. C. & Dion, V. Modifiers of CAG/CTG Repeat Instability: Insights from Mammalian Models. J Huntingtons Dis 10, 123–148 (2021). |
| *Ogg1* | | 0.9991 | *ns* | Modifier of somatic CAG expansion in R6/1 *HTT* exon1 transgenic and *Htt* .Q150 mice (Kovtun et al. 2007, Budworth et al. 2015). | Kovtun, I. V. et al. OGG1 initiates age-dependent CAG trinucleotide expansion in somatic cells. Nature 447, 447–452 (2007).; Budworth, H. et al. Suppression of somatic expansion delays the onset of pathophysiology in a mouse model of huntington's disease. PLoS Genet. 11, e1005267 (2015). |
| *Parp1* | | 0.9997 | *ns* | Poly-ADP-ribosyltransferase that plays a key role in DNA repair. Modifies CAG/CTG instability in human cells (Hubert et al. 2011). | Hubert, L., Lin, Y., Dion, V. & Wilson, J. H. Topoisomerase 1 and single-strand break repair modulate transcription-induced CAG repeat contraction in human cells. Mol. Cell. Biol. 31, 3105–3112 (2011). |
| *Pcna* | Enhancer | <0.0001 | **** | Accessory factor involved in mismatch repair and CAG/CTG loop processing (Pluciennik et al. 2013, Phadte et al. 2023). | Pluciennik, A. et al. Extrahelical (CAG)/(CTG) triplet repeat elements support proliferating cell nuclear antigen loading and MutLα endonuclease activation. Proc Natl Acad Sci USA 110, 12277–12282 (2013).; Phadte, A. S. et al. FAN1 removes triplet repeat extrusions via a PCNA- and RFC-dependent mechanism. Proc Natl Acad Sci USA 120, e2302103120 (2023). |
| *Pms1* | Enhancer | <0.0001 | **** | Candidate genetic modifier of HD age of onset (GeM-HD Consortium 2019). | Genetic Modifiers of Huntington's Disease (GeM-HD) Consortium. Identification of Genetic Factors that Modify Clinical Onset of Huntington's Disease. Cell 162, 516–526 (2015). |
| *Pms2* | Supressor | <0.0001 | **** | Candidate genetic modifier of HD age of onset (GeM-HD Consortium 2019). | Genetic Modifiers of Huntington's Disease (GeM-HD) Consortium. Identification of Genetic Factors that Modify Clinical Onset of Huntington's Disease. Cell 162, 516–526 (2015). |
| *Polb* | Enhancer | 0.0004 | *** | DNA polymerase beta involved in base excision repair. Interacts with MutSβ to promote gap-filling synthesis in base excision repair (Lai et al. 2016, Guo et al. 2016). Modifies *Fmr1* CGG instabilty (Lokanga et al. 2015). | Lai, Y. et al. Crosstalk between MSH2-MSH3 and polβ promotes trinucleotide repeat expansion during base excision repair. Nat. Commun. 7, 12465 (2016).; Guo, J., Gu, L., Leffak, M. & Li, G.-M. MutSβ promotes trinucleotide repeat expansion by recruiting DNA polymerase β to nascent (CAG)n or (CTG)n hairpins for error-prone DNA synthesis. Cell Res. 26, 775–786 (2016).; Lokanga, R. A., Senejani, A. G., Sweasy, J. B. & Usdin, K. Heterozygosity for a hypomorphic Polβ mutation reduces the expansion frequency in a mouse model of the Fragile X-related disorders. PLoS Genet. 11, e1005181 (2015). |
| *Pold1* | Enhancer | <0.0001 | **** | Subunit of DNA polymerase delta, catalyzing gap-filling DNA synthesis in mismatch repair and involved in EXO1-independent mismatch repair (Goellner et al. 2015). Amongst top scoring genes in significantly enriched pathways from HD GWAS (GeM-HD Consortium 2019, Table S5). | Goellner, E. M., Putnam, C. D. & Kolodner, R. D. Exonuclease 1-dependent and independent mismatch repair. DNA Repair (Amst) 32, 24–32 (2015).; Genetic Modifiers of Huntington's Disease (GeM-HD) Consortium. Identification of Genetic Factors that Modify Clinical Onset of Huntington's Disease. Cell 162, 516–526 (2015). |
| *Pold2* | Enhancer | <0.0001 | **** | Subunit of DNA polymerase delta, catalyzing gap-filling DNA synthesis in mismatch repair and involved in EXO1-independent mismatch repair (Goellner et al. 2015). | Goellner, E. M., Putnam, C. D. & Kolodner, R. D. Exonuclease 1-dependent and independent mismatch repair. DNA Repair (Amst) 32, 24–32 (2015). |
| *Pold3* | Enhancer | <0.0001 | **** | Subunit of DNA polymerase delta, catalyzing gap-filling DNA synthesis in mismatch repair and involved in EXO1-independent mismatch repair (Goellner et al. 2015). | Goellner, E. M., Putnam, C. D. & Kolodner, R. D. Exonuclease 1-dependent and independent mismatch repair. DNA Repair (Amst) 32, 24–32 (2015). |
| *Pold4* | Enhancer | <0.0001 | **** | Subunit of DNA polymerase delta, catalyzing gap-filling DNA synthesis in mismatch repair and involved in EXO1-independent mismatch repair (Goellner et al. 2015). | Goellner, E. M., Putnam, C. D. & Kolodner, R. D. Exonuclease 1-dependent and independent mismatch repair. DNA Repair (Amst) 32, 24–32 (2015). |
| *Pole* | Enhancer | 0.0002 | *** | DNA polymerase epsilon acts in cell-free reconstituted mismatch repair (Bowen and Kolodner 2017). | Bowen, N. & Kolodner, R. D. Reconstitution of Saccharomyces cerevisiae DNA polymerase ε-dependent mismatch repair with purified proteins. Proc Natl Acad Sci USA 114, 3607–3612 (2017). |
| *Polh* | | 0.9178 | *ns* | Tranlesion DNA polymerase eta involved in noncanonical mismatch repair (Pena-Diaz et al. 2012) | Peña-Diaz, J. et al. Noncanonical mismatch repair as a source of genomic instability in human cells. Mol. Cell 47, 669–680 (2012). |
| *Poln* | | 0.9847 | *ns* | Amongst top scoring genes in significantly enriched pathways from HD GWAS (GeM-HD Consortium 2019, Table S5). | Genetic Modifiers of Huntington's Disease (GeM-HD) Consortium. Identification of Genetic Factors that Modify Clinical Onset of Huntington's Disease. Cell 162, 516–526 (2015). |
| *Polq* | | 0.9981 | *ns* | Amongst top scoring genes in significantly enriched pathways from HD GWAS (GeM-HD Consortium 2019, Table S5). | Genetic Modifiers of Huntington's Disease (GeM-HD) Consortium. Identification of Genetic Factors that Modify Clinical Onset of Huntington's Disease. Cell 162, 516–526 (2015). |
| *Rad18* | | 0.9985 | *ns* | Amongst top scoring genes in significantly enriched pathways from HD GWAS (GeM-HD Consortium 2019, Table S5). Modifier of CAG/CTG instability in human cells (Frizzell et al. 2014). Implicated in noncanonical mismatch repair (Pena-Diaz et al. 2012). | Genetic Modifiers of Huntington's Disease (GeM-HD) Consortium. Identification of Genetic Factors that Modify Clinical Onset of Huntington's Disease. Cell 162, 516–526 (2015).; Frizzell, A. et al. RTEL1 inhibits trinucleotide repeat expansions and fragility. Cell Rep. 6, 827–835 (2014).; Peña-Diaz, J. et al. Noncanonical mismatch repair as a source of genomic instability in human cells. Mol. Cell 47, 669–680 (2012). |
| *Rnaseh1* | | 0.9998 | *ns* | Processes R-loops. Modifies instability in human cells (Lin et al. 2010) | Lin, Y., Dent, S. Y. R., Wilson, J. H., Wells, R. D. & Napierala, M. R loops stimulate genetic instability of CTG.CAG repeats. Proc Natl Acad Sci USA 107, 692–697 (2010). |
| *Rnaseh2a* | | 0.9994 | *ns* | Processes R-loops. Modifies instability in human cells (Lin et al. 2010) | Lin, Y., Dent, S. Y. R., Wilson, J. H., Wells, R. D. & Napierala, M. R loops stimulate genetic instability of CTG.CAG repeats. Proc Natl Acad Sci USA 107, 692–697 (2010). |
| *Rrm2b* | | 0.4731 | *ns* | Candidate genetic modifier of HD age of onset (GeM-HD Consortium 2015, 2019). Mild modifier of somatic CAG expansion in *Htt* .Q111 mice (Loupe et al. 2020). | Genetic Modifiers of Huntington's Disease (GeM-HD) Consortium. CAG repeat not polyglutamine length determines timing of huntington's disease onset. Cell 178, 887-900.e14 (2019).; Genetic Modifiers of Huntington's Disease (GeM-HD) Consortium. Identification of Genetic Factors that Modify Clinical Onset of Huntington's Disease. Cell 162, 516–526 (2015).; Loupe, J. M. et al. Promotion of somatic CAG repeat expansion by Fan1 knock-out in Huntington's disease knock-in mice is blocked by Mlh1 knock-out. Hum. Mol. Genet. 29, 3044–3053 (2020). |
| *Setd2* | Enhancer | 0.0174 | * | Trimethylates H3 lysine 36 (Wagner&Carpenter 2012). H3K36me3 plays a role in mismatch repair by recruting MutSα to chromatin (Huang&Li 2020) | Wagner, E. J. & Carpenter, P. B. Understanding the language of Lys36 methylation at histone H3. Nat. Rev. Mol. Cell Biol. 13, 115–126 (2012).; Huang, Y. & Li, G.-M. DNA mismatch repair in the context of chromatin. Cell Biosci. 10, 10 (2020). |
| *Setdb1* | Enhancer | 0.0492 | * | Trimethylates H3 lysine 9. Coordinates histone methylation with new histone deposition in damaged heterochromatin, thus protecting cells from genome instability (Fortuny et al. 2021). Expression is increased in HD (Ryu et al. 2006). Regulates topological chromatin domains (Jiang et al. 2017) and disease-associated repeats co-localize with chromatin domain boundaries (Sun et al. 2019). | Fortuny, A. et al. Imaging the response to DNA damage in heterochromatin domains reveals core principles of heterochromatin maintenance. Nat. Commun. 12, 2428 (2021).; Ryu, H. et al. ESET/SETDB1 gene expression and histone H3 (K9) trimethylation in Huntington's disease. Proc Natl Acad Sci USA 103, 19176–19181 (2006).; Jiang, Y. et al. The methyltransferase SETDB1 regulates a large neuron-specific topological chromatin domain. Nat. Genet. 49, 1239–1250 (2017).; Sun, Y. et al. A Dissection of Oligomerization by the TRIM28 Tripartite Motif and the Interaction with Members of the Krab-ZFP Family. J. Mol. Biol. 431, 2511–2527 (2019). |
| *Setx* | | 0.9994 | *ns* | Probable RNA/DNA helicase involved in diverse aspects of RNA metabolism and genomic integrity. Modifier of CAG/CTG instability in human cells (Nakatani et al. 2015) | Nakatani, R., Nakamori, M., Fujimura, H., Mochizuki, H. & Takahashi, M. P. Large expansion of CTG•CAG repeats is exacerbated by MutSβ in human cells. Sci. Rep. 5, 11020 (2015). |
| *Sirt6* | | 0.9996 | *ns* | Regulates histone aceylation and genome integrity (Tennen&Chua 2011) | Tennen, R. I. & Chua, K. F. Chromatin regulation and genome maintenance by mammalian SIRT6. Trends Biochem. Sci. 36, 39–46 (2011). |
| *Slx4* | | 0.9991 | *ns* | Part of a multi-subunit complex incorporating endonucleases for the resolution of DNA structures that interacts with MutSβ (Svendsen et al. 2009). MutSβ stimulates the cleavage of CAG loops by SLX1-SLX4 (Young et al. 2020). | Svendsen, J. M. et al. Mammalian BTBD12/SLX4 assembles a Holliday junction resolvase and is required for DNA repair. Cell 138, 63–77 (2009).; Young, S. J. et al. Mutsβ stimulates holliday junction resolution by the SMX complex. Cell Rep. 33, 108289 (2020). |
| *Smarcad1* | | 0.9996 | *ns* | Recruited by MSH2 to mismatch-containing DNA to exclude nucleosomes from the repair site (Terui et al. 2018). | Terui, R. et al. Nucleosomes around a mismatched base pair are excluded via an Msh2-dependent reaction with the aid of SNF2 family ATPase Smarcad1. Genes Dev. 32, 806–821 (2018). |
| *Supt4a* | | 0.9996 | *ns* | Selectively transcribes long CAG repeat tracts (Liu et al. 2012). | Liu, C.-R. et al. Spt4 is selectively required for transcription of extended trinucleotide repeats. Cell 148, 690–701 (2012). |
| *Tcea2* | | 0.9991 | *ns* | Transcriptional elongation factor. Modifier of CAG/CTG instability in human cells (Lin&Wilson 2007). | Lin, Y. & Wilson, J. H. Transcription-induced CAG repeat contraction in human cells is mediated in part by transcription-coupled nucleotide excision repair. Mol. Cell. Biol. 27, 6209–6217 (2007). |
| *Tcerg1* | | 0.9993 | *ns* | Candidate genetic modifier of HD age of onset (GeM-HD Consortium 2015, 2019). | Genetic Modifiers of Huntington's Disease (GeM-HD) Consortium. CAG repeat not polyglutamine length determines timing of huntington's disease onset. Cell 178, 887-900.e14 (2019).; Genetic Modifiers of Huntington's Disease (GeM-HD) Consortium. Identification of Genetic Factors that Modify Clinical Onset of Huntington's Disease. Cell 162, 516–526 (2015). |
| *Tdp1* | | 0.9994 | *ns* | Single strand break repair protein. Modifier of CAG/CTG instability in human cells (Hubert et al. 2011). | Hubert, L., Lin, Y., Dion, V. & Wilson, J. H. Topoisomerase 1 and single-strand break repair modulate transcription-induced CAG repeat contraction in human cells. Mol. Cell. Biol. 31, 3105–3112 (2011). |
| *Ubr5* | | 0.9289 | *ns* | Candidate genetic modifier of HD age of onset (GeM-HD Consortium 2015, 2019) | Genetic Modifiers of Huntington's Disease (GeM-HD) Consortium. CAG repeat not polyglutamine length determines timing of huntington's disease onset. Cell 178, 887-900.e14 (2019).; Genetic Modifiers of Huntington's Disease (GeM-HD) Consortium. Identification of Genetic Factors that Modify Clinical Onset of Huntington's Disease. Cell 162, 516–526 (2015). |
| *Ung* | | 0.0921 | *ns* | Base excision repair protein. Modifier of CAG/CTG instability in yeast (Su&Freudenreich 2017) | Su, X. A. & Freudenreich, C. H. Cytosine deamination and base excision repair cause R-loop-induced CAG repeat fragility and instability in Saccharomyces cerevisiae. Proc Natl Acad Sci USA 114, E8392–E8401 (2017). |
| *Wrn* | | 0.9979 | *ns* | Interacts with mismatch repair factors (Saydam et al. 2007). | Saydam, N. et al. Physical and functional interactions between Werner syndrome helicase and mismatch-repair initiation factors. Nucleic Acids Res. 35, 5706–5716 (2007). |
| *Xpa* | | 0.2713 | *ns* | Modifier of Atxn1 CAG instability in neuronal tissues (Hubert et al. 2011b) | Hubert, L., Lin, Y., Dion, V. & Wilson, J. H. Xpa deficiency reduces CAG trinucleotide repeat instability in neuronal tissues in a mouse model of SCA1. Hum. Mol. Genet. 20, 4822–4830 (2011). |

† *One way ANOVA; "Empty" used as reference group; Dunnett's multiple comparisons correction.*

***Supplementary Table 4: GO terms for target genes***

http://pantherdb.org/

Accessed on: 7/25/2023

| Gene (Mouse) | Gene (Human) | Protein (Human) | Gene Name | Custom Category used in Fig 3 | PANTHER Family/Subfamily | PANTHER Protein Class | GO Biological Process (Slim) | GO Biological Process (Complete) |
|---|---|---|---|---|---|---|---|---|
| *Atm* | *ATM* | Q13315 | Serine-protein kinase ATM | transcription/regulation by RNA Polymerase II | SERINE-PROTEIN KINASE ATM (PTHR11139:SF72) | non-receptor serine/threonine protein kinase(PC00167) | TORC1 signaling(GO:0038202);DNA damage checkpoint signaling(GO:0000077);negative regulation of TORC1 signaling(GO:1904262);cell cycle phase transition(GO:0044770);telomere maintenance(GO:0000723);protein phosphorylation(GO:0006468) | DNA damage checkpoint signaling(GO:0000077);pexophagy(GO:0000425);telomere maintenance(GO:0000723);double-strand break repair via homologous recombination(GO:0000724);ovarian follicle development(GO:0001541);response to hypoxia(GO:0001666);somitogenesis(GO:0001756);pre-B cell allelic exclusion(GO:0002331);double-strand break repair(GO:0006302);double-strand break repair via nonhomologous end joining(GO:0006303);protein phosphorylation(GO:0006468);DNA damage response(GO:0006974);DNA damage response, signal transduction by p53 class mediator resulting in cell cycle arrest(GO:0006977);mitotic spindle assembly checkpoint signaling(GO:0007094);mitotic G2 DNA damage checkpoint signaling(GO:0007095);reciprocal meiotic recombination(GO:0007131);male meiotic nuclear division(GO:0007140);female meiotic nuclear division(GO:0007143);signal transduction(GO:0007165);brain development(GO:0007420);heart development(GO:0007507);determination of adult lifespan(GO:0008340);intrinsic apoptotic signaling pathway in response to DNA damage(GO:0008630);post-embryonic development(GO:0009791);response to ionizing radiation(GO:0010212);regulation of autophagy(GO:0010506);positive regulation of gene expression(GO:0010628);peptidyl-serine phosphorylation(GO:0018105);positive regulation of cell migration(GO:0030335);negative regulation of B cell proliferation(GO:0030889);regulation of telomere maintenance via telomerase(GO:0032210);positive regulation of telomere maintenance via telomerase(GO:0032212);V(D)J recombination(GO:0033151);cellular response to reactive oxygen species(GO:0034614);multicellular organism growth(GO:0035264);phosphatidylinositol-3-phosphate biosynthetic process(GO:0036092);peptidyl-serine autophosphorylation(GO:0036289);lipoprotein catabolic process(GO:0042159);signal transduction in response to DNA damage(GO:0042770);regulation of apoptotic process(GO:0042981);positive regulation of apoptotic process(GO:0043065);positive regulation of DNA damage response, signal transduction by p53 class mediator(GO:0043517);positive regulation of neuron apoptotic process(GO:0043525);meiotic telomere clustering(GO:0045141);positive regulation of cell adhesion(GO:0045785);positive regulation of transcription by RNA polymerase II(GO:0045944);protein autophosphorylation(GO:0046777);thymus development(GO:0048538);oocyte development(GO:0048599);neuron apoptotic process(GO:0051402);regulation of cell cycle(GO:0051726);histone mRNA catabolic process(GO:0071044);cellular response to retinoic acid(GO:0071300);cellular response to gamma radiation(GO:0071480);cellular response to X-ray(GO:0071481);cellular response to nitrosative stress(GO:0071500);cellular senescence(GO:0090398);replicative senescence(GO:0090399);establishment of RNA localization to telomere(GO:0097694);establishment of protein-containing complex localization to telomere(GO:0097695);regulation of cellular response to heat(GO:1900034);regulation of signal transduction by p53 class mediator(GO:1901796);positive regulation of DNA catabolic process(GO:1903626);regulation of microglial cell activation(GO:1903978);negative regulation of TORC1 signaling(GO:1904262);negative regulation of telomere capping(GO:1904354);positive regulation of telomere maintenance via telomere lengthening(GO:1904358);positive regulation of telomerase catalytic core complex assembly(GO:1904884);regulation of cellular response to gamma radiation(GO:1905843) |
| *Brip1 (Fancj)* | *BRIP1 (FANCJ)* | Q9BX63 | Fanconi anemia group J protein | | FANCONI ANEMIA GROUP J PROTEIN (PTHR11472:SF47) | DNA helicase(PC00011) | reciprocal meiotic recombination(GO:0007131);nucleotide-excision repair(GO:0006289);meiotic telophase I(GO:0007134);double-strand break repair(GO:0006302) | DNA damage checkpoint signaling(GO:0000077);DNA repair(GO:0006281);nucleotide-excision repair(GO:0006289);double-strand break repair(GO:0006302);regulation of transcription by RNA polymerase II(GO:0006357);spermatogonial cell division(GO:0007284);spermatid development(GO:0007286);negative regulation of cell population proliferation(GO:0008285);response to toxic substance(GO:0009636);negative regulation of gene expression(GO:0010629);meiotic DNA double-strand break processing involved in reciprocal meiotic recombination(GO:0010705);DNA duplex unwinding(GO:0032508);homologous recombination(GO:0035825);chiasma assembly(GO:0051026);cellular response to vitamin(GO:0071295);cellular response to hypoxia(GO:0071456);seminiferous tubule development(GO:0072520);cellular response to angiotensin(GO:1904385);double-strand break repair involved in meiotic recombination(GO:1990918) |
| *Ccdc82* | *CCDC82* | Q8N4S0 | Coiled-coil domain-containing protein 82 | GWAS candidate | COILED-COIL DOMAIN-CONTAINING PROTEIN 82 (PTHR14689:SF0) | | | |
| *Crebbp* | *CREBBP* | Q92793 | CREB-binding protein | transcription/regulation by RNA Polymerase II | CREB-BINDING PROTEIN (PTHR13808:SF34) | histone modifying enzyme(PC00261) | transcription by RNA polymerase II(GO:0006366);histone acetylation(GO:0016573);positive regulation of transcription by RNA polymerase II(GO:0045944) | negative regulation of transcription by RNA polymerase II(GO:0000122);response to hypoxia(GO:0001666);stimulatory C-type lectin receptor signaling pathway(GO:0002223);chromatin remodeling(GO:0006338);DNA-templated transcription(GO:0006351);regulation of DNA-templated transcription(GO:0006355);protein acetylation(GO:0006473);signal transduction(GO:0007165);regulation of smoothened signaling pathway(GO:0008589);negative regulation of transcription by RNA polymerase I(GO:0016479);histone acetylation(GO:0016573);N-terminal peptidyl-lysine acetylation(GO:0018076);positive regulation of transforming growth factor beta receptor signaling pathway(GO:0030511);protein destabilization(GO:0031648);cellular response to nutrient levels(GO:0031669);cellular response to UV(GO:0034644);NIK/NF-kappaB signaling(GO:0038061);homeostatic process(GO:0042592);embryonic digit morphogenesis(GO:0042733);positive regulation of DNA-templated transcription(GO:0045893);positive regulation of transcription by RNA polymerase II(GO:0045944);rhythmic process(GO:0048511);protein-containing complex assembly(GO:0065003);regulation of cellular response to heat(GO:1900034);positive regulation of protein localization to nucleus(GO:1900182) |
| *Dhfr* | *DHFR* | P00374 | Dihydrofolate reductase | GWAS candidate | DIHYDROFOLATE REDUCTASE (PTHR48069:SF6) | reductase(PC00198) | cellular amide metabolic process(GO:0043603);cellular nitrogen compound biosynthetic process(GO:0044271);heterocycle biosynthetic process(GO:0018130);organic cyclic compound biosynthetic process(GO:1901362);carboxylic acid metabolic process(GO:0019752);folic acid-containing compound metabolic process(GO:0006760);organonitrogen compound biosynthetic process(GO:1901566);aromatic compound biosynthetic process(GO:0019438) | glycine biosynthetic process(GO:0006545);tetrahydrobiopterin biosynthetic process(GO:0006729);one-carbon metabolic process(GO:0006730);negative regulation of translation(GO:0017148);axon regeneration(GO:0031103);response to methotrexate(GO:0031427);dihydrofolate metabolic process(GO:0046452);tetrahydrofolate metabolic process(GO:0046653);tetrahydrofolate biosynthetic process(GO:0046654);folic acid metabolic process(GO:0046655);positive regulation of nitric-oxide synthase activity(GO:0051000);regulation of removal of superoxide radicals(GO:2000121) |
| *Dnmt1* | *DNMT1* | P26358 | DNA (cytosine-5)-methyltransferase 1 | transcription/regulation by RNA Polymerase II | DNA (CYTOSINE-5)-METHYLTRANSFERASE 1 (PTHR10629:SF52) | DNA methyltransferase(PC00013) | | negative regulation of transcription by RNA polymerase II(GO:0000122);DNA methylation-dependent heterochromatin formation(GO:0006346);DNA-templated transcription(GO:0006351);DNA methylation on cytosine within a CG sequence(GO:0010424);positive regulation of gene expression(GO:0010628);negative regulation of gene expression(GO:0010629);methylation(GO:0032259);DNA methylation involved in embryo development(GO:0043045);negative regulation of gene expression via CpG island methylation(GO:0044027);cellular response to amino acid stimulus(GO:0071230);C-5 methylation of cytosine(GO:0090116);cellular response to bisphenol A(GO:1903926);positive regulation of vascular associated smooth muscle cell proliferation(GO:1904707);negative regulation of vascular associated smooth muscle cell apoptotic process(GO:1905460);negative regulation of vascular associated smooth muscle cell differentiation involved in phenotypic switching(GO:1905931) |
| *Dnmt3a* | *DNMT3A* | Q9Y6K1 | DNA (cytosine-5)-methyltransferase 3A | transcription/regulation by RNA Polymerase II | DNA (CYTOSINE-5)-METHYLTRANSFERASE 3A (PTHR23068:SF10) | DNA methyltransferase(PC00013) | negative regulation of transcription, DNA-templated(GO:0045892);transcription, DNA-templated(GO:0006351);DNA metabolic process(GO:0006259);macromolecule methylation(GO:0043414) | negative regulation of transcription by RNA polymerase II(GO:0000122);mitotic cell cycle(GO:0000278);DNA methylation(GO:0006306);DNA methylation-dependent heterochromatin formation(GO:0006346);regulation of gene expression by genomic imprinting(GO:0006349);spermatogenesis(GO:0007283);response to xenobiotic stimulus(GO:0009410);response to toxic substance(GO:0009636);response to ionizing radiation(GO:0010212);response to lead ion(GO:0010288);regulation of gene expression(GO:0010468);neuron differentiation(GO:0030182);response to estradiol(GO:0032355);DNA methylation on cytosine(GO:0032776);response to vitamin A(GO:0033189);response to cocaine(GO:0042220);DNA methylation involved in embryo development(GO:0043045);negative regulation of DNA-templated transcription(GO:0045892);cellular response to amino acid stimulus(GO:0071230);cellular response to ethanol(GO:0071361);cellular response to hypoxia(GO:0071456);genomic imprinting(GO:0071514);C-5 methylation of cytosine(GO:0090116);hepatocyte apoptotic process(GO:0097284);cellular response to bisphenol A(GO:1903926) |
| *Ercc1* | *ERCC1* | P07992 | DNA excision repair protein ERCC-1 | | DNA EXCISION REPAIR PROTEIN ERCC-1 (PTHR12749:SF0) | endodeoxyribonuclease(PC00093) | cellular response to light stimulus(GO:0071482);meiosis I cell cycle process(GO:0061982);nucleotide-excision repair, DNA incision(GO:0033683);response to UV(GO:0009411);mismatch repair(GO:0006298);mitotic recombination(GO:0006312);meiotic nuclear division(GO:0140013) | pyrimidine dimer repair by nucleotide-excision repair(GO:0000720);DNA repair(GO:0006281);nucleotide-excision repair(GO:0006289);double-strand break repair via nonhomologous end joining(GO:0006303);mitotic recombination(GO:0006312);syncytium formation(GO:0006949);response to oxidative stress(GO:0006979);spermatogenesis(GO:0007283);response to nutrient(GO:0007584);cell population proliferation(GO:0008283);determination of adult lifespan(GO:0008340);male gonad development(GO:0008584);UV protection(GO:0009650);response to sucrose(GO:0009744);response to X-ray(GO:0010165);negative regulation of telomere maintenance(GO:0032205);post-embryonic hemopoiesis(GO:0035166);multicellular organism growth(GO:0035264);response to immobilization stress(GO:0035902);interstrand cross-link repair(GO:0036297);isotype switching(GO:0045190);response to cadmium ion(GO:0046686);insulin-like growth factor receptor signaling pathway(GO:0048009);oogenesis(GO:0048477);embryonic organ development(GO:0048568);positive regulation of transcription initiation by RNA polymerase II(GO:0060261);telomeric DNA-containing double minutes formation(GO:0061819);UV-damage excision repair(GO:0070914);replicative senescence(GO:0090399);t-circle formation(GO:0090656);positive regulation of t-circle formation(GO:1904431);negative regulation of protection from non-homologous end joining at telomere(GO:1905765) |
| *Ercc3* | *ERCC3* | P19447 | General transcription and DNA repair factor IIH helicase subunit XPB | transcription/regulation by RNA Polymerase II | GENERAL TRANSCRIPTION AND DNA REPAIR FACTOR IIH HELICASE SUBUNIT XPB (PTHR11274:SF0) | DNA helicase(PC00011) | | nucleotide-excision repair, DNA duplex unwinding(GO:0000717);response to hypoxia(GO:0001666);DNA topological change(GO:0006265);DNA repair(GO:0006281);transcription-coupled nucleotide-excision repair(GO:0006283);nucleotide-excision repair(GO:0006289);transcription by RNA polymerase II(GO:0006366);transcription initiation at RNA polymerase II promoter(GO:0006367);transcription elongation by RNA polymerase II(GO:0006368);apoptotic process(GO:0006915);response to oxidative stress(GO:0006979);protein localization(GO:0008104);response to UV(GO:0009411);UV protection(GO:0009650);DNA duplex unwinding(GO:0032508);hair cell differentiation(GO:0035315);positive regulation of apoptotic process(GO:0043065);embryonic organ development(GO:0048568);regulation of mitotic cell cycle phase transition(GO:1901990) |
| *Ercc5* | *ERCC5* | P28715 | DNA excision repair protein ERCC-5 | | DNA EXCISION REPAIR PROTEIN ERCC-5 (PTHR16171:SF7) | DNA metabolism protein(PC00009) | nucleotide-excision repair, DNA incision(GO:0033683) | double-strand break repair via homologous recombination(GO:0000724);transcription-coupled nucleotide-excision repair(GO:0006283);base-excision repair, AP site formation(GO:0006285);nucleotide-excision repair(GO:0006289);response to UV(GO:0009411);response to UV-C(GO:0010225);negative regulation of apoptotic process(GO:0043066);regulation of catalytic activity(GO:0050790) |
| *Exo1* | *EXO1* | Q9UQ84 | Exonuclease 1 | | EXONUCLEASE 1 (PTHR11081:SF8) | exodeoxyribonuclease(PC00098) | | humoral immune response mediated by circulating immunoglobulin(GO:0002455);DNA repair(GO:0006281);mismatch repair(GO:0006298);DNA recombination(GO:0006310);somatic hypermutation of immunoglobulin genes(GO:0016446);isotype switching(GO:0045190);meiotic cell cycle(GO:0051321);RNA phosphodiester bond hydrolysis, endonucleolytic(GO:0090502);t-circle formation(GO:0090656) |
| *Fan1* | *FAN1* | Q9Y2M0 | Fanconi-associated nuclease 1 | GWAS candidate | FANCONI-ASSOCIATED NUCLEASE 1 (PTHR15749:SF4) | | | double-strand break repair via homologous recombination(GO:0000724);DNA repair(GO:0006281);nucleotide-excision repair(GO:0006289);interstrand cross-link repair(GO:0036297) |
| *Fen1* | *FEN1* | P39748 | Flap endonuclease 1 | | FLAP ENDONUCLEASE 1 (PTHR11081:SF58) | exodeoxyribonuclease(PC00098) | | double-strand break repair via homologous recombination(GO:0000724);DNA replication(GO:0006260);DNA repair(GO:0006281);base-excision repair, gap-filling(GO:0006287);double-strand break repair(GO:0006302);memory(GO:0007613);UV protection(GO:0009650);telomere maintenance via semi-conservative replication(GO:0032201);DNA replication, removal of RNA primer(GO:0043137);positive regulation of sister chromatid cohesion(GO:0045876);nucleic acid phosphodiester bond hydrolysis(GO:0090305);RNA phosphodiester bond hydrolysis, endonucleolytic(GO:0090502) |
| *Gpr151* | *GPR151* | Q8TDV0 | G-protein coupled receptor 151 | GWAS candidate | G-PROTEIN COUPLED RECEPTOR 151 (PTHR45695:SF1) | transmembrane signal receptor(PC00197) | G protein-coupled receptor signaling pathway(GO:0007186) | response to ischemia(GO:0002931);G protein-coupled receptor signaling pathway(GO:0007186);neuropeptide signaling pathway(GO:0007218);response to acidic pH(GO:0010447);positive regulation of immune response(GO:0050778) |
| *Hdac6* | *HDAC6* | Q9UBN7 | Histone deacetylase 6 | transcription/regulation by RNA Polymerase II | HISTONE DEACETYLASE 6 (PTHR48252:SF71) | histone modifying enzyme(PC00261) | | negative regulation of transcription by RNA polymerase II(GO:0000122);protein polyubiquitination(GO:0000209);response to amphetamine(GO:0001975);chromatin remodeling(GO:0006338);protein deacetylation(GO:0006476);protein quality control for misfolded or incompletely synthesized proteins(GO:0006515);nitrogen compound metabolic process(GO:0006807);intracellular protein transport(GO:0006886);autophagy(GO:0006914);negative regulation of microtubule depolymerization(GO:0007026);regulation of autophagy(GO:0010506);negative regulation of macromolecule metabolic process(GO:0010605);positive regulation of epithelial cell migration(GO:0010634);negative regulation of hydrogen peroxide metabolic process(GO:0010727);regulation of macroautophagy(GO:0016241);histone deacetylation(GO:0016575);axonal transport of mitochondrion(GO:0019896);negative regulation of protein-containing complex assembly(GO:0031333);regulation of protein stability(GO:0031647);protein destabilization(GO:0031648);lysosome localization(GO:0032418);protein-containing complex disassembly(GO:0032984);positive regulation of peptidyl-serine phosphorylation(GO:0033138);cellular response to stress(GO:0033554);peptidyl-lysine deacetylation(GO:0034983);response to immobilization stress(GO:0035902);cellular response to topologically incorrect protein(GO:0035967);epigenetic regulation of gene expression(GO:0040029);ubiquitin-dependent protein catabolic process via the multivesicular body sorting pathway(GO:0043162);negative regulation of protein-containing complex disassembly(GO:0043242);regulation of fat cell differentiation(GO:0045598);negative regulation of proteolysis(GO:0045861);negative regulation of DNA-templated transcription(GO:0045892);collateral sprouting(GO:0048668);negative regulation of axon extension involved in axon guidance(GO:0048843);positive regulation of dendrite morphogenesis(GO:0050775);negative regulation of oxidoreductase activity(GO:0051354);response to corticosterone(GO:0051412);response to misfolded protein(GO:0051788);positive regulation of synaptic transmission, glutamatergic(GO:0051968);cilium assembly(GO:0060271);regulation of microtubule-based movement(GO:0060632);regulation of androgen receptor signaling pathway(GO:0060765);dendritic spine morphogenesis(GO:0060997);cilium disassembly(GO:0061523);parkin-mediated stimulation of mitophagy in response to mitochondrial depolarization(GO:0061734);regulation of establishment of protein localization(GO:0070201);cellular response to hydrogen peroxide(GO:0070301);aggresome assembly(GO:0070842);polyubiquitinated misfolded protein transport(GO:0070845);Hsp90 deacetylation(GO:0070846);response to growth factor(GO:0070848);cellular response to misfolded protein(GO:0071218);cellular response to parathyroid hormone stimulus(GO:0071374);response to dexamethasone(GO:0071548);tubulin deacetylation(GO:0090042);positive regulation of tubulin deacetylation(GO:0090044);positive regulation of hydrogen peroxide-mediated programmed cell death(GO:1901300);negative regulation of protein acetylation(GO:1901984);regulation of autophagy of mitochondrion(GO:1903146);positive regulation of cholangiocyte proliferation(GO:1904056);positive regulation of signaling receptor activity(GO:2000273) |
| *Hmgb1* | *HMGB1* | P09429 | High mobility group protein B1 | transcription/regulation by RNA Polymerase II | HIGH MOBILITY GROUP PROTEIN B1 (PTHR48112:SF11) | chromatin/chromatin-binding, or -regulatory protein(PC00077) | transcription by RNA polymerase II(GO:0006366);regulation of transcription by RNA polymerase II(GO:0006357) | negative regulation of transcription by RNA polymerase II(GO:0000122);myeloid dendritic cell activation(GO:0001773);activation of innate immune response(GO:0002218);dendritic cell chemotaxis(GO:0002407);inflammatory response to antigenic stimulus(GO:0002437);regulation of tolerance [illegible] |
| *Htt* | *HTT* | P42858 | Huntingtin | GWAS candidate | HUNTINGTIN (PTHR10170:SF10) | | | [illegible] |
| *Lig1* | *LIG1* | P18858 | DNA ligase 1 | GWAS candidate | DNA LIGASE 1 (PTHR45674:SF4) | DNA ligase(PC00012) | | [illegible] |
| *Lig3* | *LIG3* | P49916 | DNA ligase 3 | | DNA LIGASE 3 (PTHR45674:SF9) | DNA ligase(PC00012) | DNA strand elongation involved in DNA replication(GO:0006271);DNA biosynthetic process(GO:0071897);RNA biosynthetic process(GO:0032774);RNA catabolic process(GO:0006401) | [illegible] |
| *Lig4* | *LIG4* | P49917 | DNA ligase 4 | | DNA LIGASE 4 (PTHR45997:SF1) | DNA metabolism protein(PC00009) | nucleotide-excision repair(GO:0006289);double-strand break repair via nonhomologous end joining(GO:0006303) | [illegible] |
| *Mcm9* | *MCM9* | Q9NXL9 | DNA helicase MCM9 | | DNA HELICASE MCM9 (PTHR11630:SF48) | DNA metabolism protein(PC00009) | double-strand break repair via homologous recombination(GO:0000724) | [illegible] |
| *Mlh1* | *MLH1* | P40692 | DNA mismatch repair protein Mlh1 | GWAS candidate | DNA MISMATCH REPAIR PROTEIN MLH1 (PTHR10073:SF12) | DNA metabolism protein(PC00009) | mismatch repair(GO:0006298) | [illegible] |
| *Mlh3* | *MLH3* | Q9UHC1 | DNA mismatch repair protein Mlh3 | | DNA MISMATCH REPAIR PROTEIN MLH3 (PTHR10073:SF47) | DNA metabolism protein(PC00009) | mismatch repair(GO:0006298) | [illegible] |
| *Msh2* | *MSH2* | P43246 | DNA mismatch repair protein Msh2 | | DNA MISMATCH REPAIR PROTEIN MSH2 (PTHR11361:SF35) | DNA metabolism protein(PC00009) | mismatch repair(GO:0006298);mitotic recombination(GO:0006312) | [illegible] |
| *Msh3* | *MSH3* | P20585 | DNA mismatch repair protein Msh3 | GWAS candidate | DNA MISMATCH REPAIR PROTEIN MSH3 (PTHR11361:SF122) | DNA metabolism protein(PC00009) | mismatch repair(GO:0006298);mitotic recombination(GO:0006312) | [illegible] |
| *Msh4* | *MSH4* | O15457 | MutS protein homolog 4 | | MUTS PROTEIN HOMOLOG 4 (PTHR11361:SF21) | DNA metabolism protein(PC00009) | reciprocal meiotic recombination(GO:0007131);meiotic telophase I(GO:0007134) | [illegible] |
| *Msh5* | *MSH5* | O43196 | MutS protein homolog 5 | | MUTS PROTEIN HOMOLOG 5 (PTHR11361:SF20) | DNA metabolism protein(PC00009) | reciprocal meiotic recombination(GO:0007131);cellular component assembly(GO:0022607);meiotic telophase I(GO:0007134);homologous chromosome pairing at meiosis(GO:0007129) | [illegible] |
| *Msh6* | *MSH6* | P52701 | DNA mismatch repair protein Msh6 | | DNA MISMATCH REPAIR PROTEIN MSH6 (PTHR11361:SF148) | DNA metabolism protein(PC00009) | mismatch repair(GO:0006298) | [illegible] |
| *Ogg1* | *OGG1* | O15527 | N-glycosylase/DNA lyase | | N-GLYCOSYLASE/DNA LYASE (PTHR10242:SF2) | DNA metabolism protein(PC00009) | base-excision repair(GO:0006284) | [illegible] |
| *Parp1* | *PARP1* | P09874 | Poly [ADP-ribose] polymerase 1 | transcription/regulation by RNA Polymerase II | POLY [ADP-RIBOSE] POLYMERASE 1 (PTHR10459:SF112) | DNA metabolism protein(PC00009) | | [illegible] response(GO:0006974);mitochondrion organization(GO:0007005);transforming growth factor beta receptor signaling pathway(GO:0007179);response to gamma radiation(GO:0010332);positive regulation of cardiac muscle hypertrophy(GO:0010613);regulation of SMAD protein complex assembly(GO:0010990);carbohydrate biosynthetic process(GO:0016051);protein autoprocessing(GO:0016540);signal transduction involved in regulation of gene expression(GO:0023019);macrophage differentiation(GO:0030225);DNA ADP-ribosylation(GO:0030592);mitochondrial DNA metabolic process(GO:0032042);positive regulation of DNA-templated transcription, elongation(GO:0032786);cellular response to insulin stimulus(GO:0032869);regulation of protein localization(GO:0032880);positive regulation of intracellular estrogen receptor signaling pathway(GO:0033148);negative regulation of transcription elongation by RNA polymerase II(GO:0034244);cellular response to oxidative stress(GO:0034599);cellular response to UV(GO:0034644);protein modification process(GO:0036211);positive regulation of I-kappaB kinase/NF-kappaB signaling(GO:0043123);mitochondrial DNA repair(GO:0043504);regulation of DNA methylation(GO:0044030);innate immune response(GO:0045087);regulation of circadian sleep/wake cycle, non-REM sleep(GO:0045188);negative regulation of innate immune response(GO:0045824);negative regulation of DNA-templated transcription(GO:0045892);positive regulation of transcription by RNA polymerase II(GO:0045944);decidualization(GO:0046697);regulation of catalytic activity(GO:0050790);regulation of DNA metabolic process(GO:0051052);positive regulation of mitochondrial depolarization(GO:0051901);positive regulation of SMAD protein signal transduction(GO:0060391);positive regulation of necroptotic process(GO:0060545);protein poly-ADP-ribosylation(GO:0070212);protein auto-ADP-ribosylation(GO:0070213);protein localization to chromatin(GO:0071168);cellular response to zinc ion(GO:0071294);replication fork reversal(GO:0071932);positive regulation of protein localization to nucleus(GO:1900182);positive regulation of neuron death(GO:1901216);regulation of oxidative stress-induced neuron intrinsic apoptotic signaling pathway(GO:1903376);positive regulation of single strand break repair(GO:1903518);response to aldosterone(GO:1904044);negative regulation of adipose tissue development(GO:1904178);negative regulation of telomere maintenance via telomere lengthening(GO:1904357);cellular response to amyloid-beta(GO:1904646);positive regulation of myofibroblast differentiation(GO:1904762);regulation of base-excision repair(GO:1905051);positive regulation of double-strand break repair via homologous recombination(GO:1905168);cellular response to nerve growth factor stimulus(GO:1990090);ATP generation from poly-ADP-D-ribose(GO:1990966);positive regulation of transcription regulatory region DNA binding(GO:2000679);negative regulation of ATP biosynthetic process(GO:2001170) |
| *Pcna* | *PCNA* | P12004 | Proliferating cell nuclear antigen | transcription/regulation by RNA Polymerase II | PROLIFERATING CELL NUCLEAR ANTIGEN (PTHR11352:SF5) | DNA polymerase processivity factor(PC00015) | DNA strand elongation involved in DNA replication(GO:0006271);RNA biosynthetic process(GO:0032774);mismatch repair(GO:0006298);translesion synthesis(GO:0019985);RNA catabolic process(GO:0006401) | negative regulation of transcription by RNA polymerase II(GO:0000122);leading strand elongation(GO:0006272);base-excision repair, gap-filling(GO:0006287);mismatch repair(GO:0006298);heart development(GO:0007507);translesion synthesis(GO:0019985);epithelial cell differentiation(GO:0030855);replication fork processing(GO:0031297);positive regulation of deoxyribonuclease activity(GO:0032077);response to estradiol(GO:0032355);cellular response to UV(GO:0034644);estrous cycle(GO:0044849);positive regulation of DNA repair(GO:0045739);positive regulation of DNA replication(GO:0045740);response to cadmium ion(GO:0046686);cellular response to hydrogen peroxide(GO:0070301);cellular response to xenobiotic stimulus(GO:0071466);response to dexamethasone(GO:0071548);liver regeneration(GO:0097421);positive regulation of DNA-directed DNA polymerase activity(GO:1900264);response to L-glutamate(GO:1902065);mitotic telomere maintenance via semi-conservative replication(GO:1902990) |
| *Pms1* | *PMS1* | P54277 | PMS1 protein homolog 1 | GWAS candidate | PMS1 PROTEIN HOMOLOG 1 (PTHR10073:SF54) | DNA metabolism protein(PC00009) | mismatch repair(GO:0006298) | mismatch repair(GO:0006298);response to xenobiotic stimulus(GO:0009410) |
| *Pms2* | *PMS2* | P54278 | Mismatch repair endonuclease PMS2 | GWAS candidate | MISMATCH REPAIR ENDONUCLEASE PMS2 (PTHR10073:SF52) | DNA metabolism protein(PC00009) | mismatch repair(GO:0006298) | mismatch repair(GO:0006298);response to xenobiotic stimulus(GO:0009410);somatic hypermutation of immunoglobulin genes(GO:0016446) |
| *Polb* | *POLB* | P06746 | DNA polymerase beta | | DNA POLYMERASE BETA (PTHR11276:SF42) | DNA-directed DNA polymerase(PC00018) | double-strand break repair via nonhomologous end joining(GO:0006303) | in utero embryonic development(GO:0001701);DNA replication(GO:0006260);DNA-templated DNA replication(GO:0006261);DNA repair(GO:0006281);base-excision repair(GO:0006284);base-excision repair, gap-filling(GO:0006287);pyrimidine dimer repair(GO:0006290);double-strand break repair via nonhomologous end joining(GO:0006303);inflammatory response(GO:0006954);DNA damage response(GO:0006974);salivary gland morphogenesis(GO:0007435);intrinsic apoptotic signaling pathway in response to DNA damage(GO:0008630);response to gamma radiation(GO:0010332);somatic hypermutation of immunoglobulin genes(GO:0016446);response to ethanol(GO:0045471);lymph node development(GO:0048535);spleen development(GO:0048536);homeostasis of number of cells(GO:0048872);neuron apoptotic process(GO:0051402);response to hyperoxia(GO:0055093);immunoglobulin heavy chain V-D-J recombination(GO:0071707);DNA biosynthetic process(GO:0071897) |
| *Pold1* | *POLD1* | P28340 | DNA polymerase delta catalytic subunit | | DNA POLYMERASE DELTA CATALYTIC SUBUNIT (PTHR10322:SF23) | DNA metabolism protein(PC00009) | | DNA synthesis involved in DNA repair(GO:0000731);DNA replication(GO:0006260);DNA-templated DNA replication(GO:0006261);DNA repair(GO:0006281);base-excision repair, gap-filling(GO:0006287);nucleotide-excision repair, DNA gap filling(GO:0006297);response to UV(GO:0009411);cellular response to UV(GO:0034644);DNA replication proofreading(GO:0045004);fatty acid homeostasis(GO:0055089);error-free translesion synthesis(GO:0070987);DNA biosynthetic process(GO:0071897) |
| *Pold2* | *POLD2* | P49005 | DNA polymerase delta subunit 2 | | DNA POLYMERASE DELTA SUBUNIT 2 (PTHR10416:SF0) | DNA-directed DNA polymerase(PC00018) | DNA strand elongation involved in DNA replication(GO:0006271);DNA biosynthetic process(GO:0071897) | DNA replication(GO:0006260);DNA-templated DNA replication(GO:0006261);DNA strand elongation involved in DNA replication(GO:0006271);DNA repair(GO:0006281);error-prone translesion synthesis(GO:0042276);DNA biosynthetic process(GO:0071897) |
| *Pold3* | *POLD3* | Q15054 | DNA polymerase delta subunit 3 | | DNA POLYMERASE DELTA SUBUNIT 3 (PTHR17598:SF13) | DNA metabolism protein(PC00009) | cellular response to light stimulus(GO:0071482);DNA strand elongation involved in DNA replication(GO:0006271);nucleotide-excision repair(GO:0006289);response to UV(GO:0009411);DNA synthesis involved in DNA repair(GO:0000731) | DNA synthesis involved in DNA repair(GO:0000731);DNA replication(GO:0006260);DNA-templated DNA replication(GO:0006261);DNA strand elongation involved in DNA replication(GO:0006271);nucleotide-excision repair, DNA gap filling(GO:0006297);mismatch repair(GO:0006298);error-prone translesion synthesis(GO:0042276);DNA biosynthetic process(GO:0071897);DNA synthesis involved in UV-damage excision repair(GO:1904161) |
| *Pold4* | *POLD4* | Q9HCU8 | DNA polymerase delta subunit 4 | | DNA POLYMERASE DELTA SUBUNIT 4 (PTHR14303:SF0) | DNA-directed DNA polymerase(PC00018) | DNA-dependent DNA replication(GO:0006261);DNA synthesis involved in DNA repair(GO:0000731) | DNA synthesis involved in DNA repair(GO:0000731);positive regulation of endothelial cell proliferation(GO:0001938);DNA-templated DNA replication(GO:0006261) |
| *Pole* | *POLE* | Q07864 | DNA polymerase epsilon catalytic subunit A | | DNA POLYMERASE EPSILON CATALYTIC SUBUNIT A (PTHR10670:SF0) | DNA metabolism protein(PC00009) | base-excision repair(GO:0006284);DNA strand elongation involved in DNA replication(GO:0006271);DNA biosynthetic process(GO:0071897);mitotic nuclear division(GO:0140014);RNA biosynthetic process(GO:0032774);nucleotide-excision repair(GO:0006289);RNA catabolic process(GO:0006401) | G1/S transition of mitotic cell cycle(GO:0000082);mitotic cell cycle(GO:0000278);DNA synthesis involved in DNA repair(GO:0000731);DNA replication(GO:0006260);DNA-templated DNA replication(GO:0006261);leading strand elongation(GO:0006272);DNA repair(GO:0006281);base-excision repair, gap-filling(GO:0006287);nucleotide-excision repair, DNA gap filling(GO:0006297);DNA replication proofreading(GO:0045004);embryonic organ development(GO:0048568);DNA biosynthetic process(GO:0071897) |
| *Polh* | *POLH* | Q9Y253 | DNA polymerase eta | | DNA POLYMERASE ETA (PTHR45873:SF1) | DNA metabolism protein(PC00009) | response to radiation(GO:0009314);translesion synthesis(GO:0019985) | DNA synthesis involved in DNA repair(GO:0000731);DNA replication(GO:0006260);DNA repair(GO:0006281);regulation of DNA repair(GO:0006282);pyrimidine dimer repair(GO:0006290);response to radiation(GO:0009314);response to UV-C(GO:0010225);error-prone translesion synthesis(GO:0042276);error-free translesion synthesis(GO:0070987);cellular response to UV-C(GO:0071494) |
| *Poln* | *POLN* | Q7Z5Q5 | DNA polymerase nu | | DNA POLYMERASE NU (PTHR10133:SF27) | DNA-directed DNA polymerase(PC00018) | double-strand break repair(GO:0006302) | double-strand break repair via homologous recombination(GO:0000724);DNA-templated DNA replication(GO:0006261);DNA repair(GO:0006281);double-strand break repair(GO:0006302);translesion synthesis(GO:0019985);interstrand cross-link repair(GO:0036297);DNA biosynthetic process(GO:0071897) |
| *Polq* | *POLQ* | O75417 | DNA polymerase theta | | DNA POLYMERASE THETA (PTHR10133:SF62) | DNA-directed DNA polymerase(PC00018) | double-strand break repair(GO:0006302) | DNA-templated DNA replication(GO:0006261);DNA repair(GO:0006281);base-excision repair(GO:0006284);double-strand break repair(GO:0006302);DNA damage response(GO:0006974);somatic hypermutation of immunoglobulin genes(GO:0016446);DNA duplex unwinding(GO:0032508);protein homooligomerization(GO:0051260);DNA biosynthetic process(GO:0071897);double-strand break repair via alternative nonhomologous end joining(GO:0097681);negative regulation of double-strand break repair via homologous recombination(GO:2000042) |
| *Rad18* | *RAD18* | Q9NS91 | E3 ubiquitin-protein ligase RAD18 | | E3 UBIQUITIN-PROTEIN LIGASE RAD18 (PTHR14134:SF2) | ubiquitin-protein ligase(PC00234) | protein monoubiquitination(GO:0006513);postreplication repair(GO:0006301) | DNA repair(GO:0006281);postreplication repair(GO:0006301);protein monoubiquitination(GO:0006513);DNA damage response(GO:0006974);protein autoubiquitination(GO:0051865);positive regulation of chromosome segregation(GO:0051984) |
| *Rnaseh1* | *RNASEH1* | O60930 | Ribonuclease H1 | | RIBONUCLEASE H1 (PTHR10642:SF26) | endoribonuclease(PC00094) | RNA phosphodiester bond hydrolysis, endonucleolytic(GO:0090502);DNA replication(GO:0006260);DNA biosynthetic process(GO:0071897);RNA catabolic process(GO:0006401) | RNA catabolic process(GO:0006401);DNA replication, removal of RNA primer(GO:0043137);RNA phosphodiester bond hydrolysis, endonucleolytic(GO:0090502) |
| *Rnaseh2a* | *RNASEH2A* | O75792 | Ribonuclease H2 subunit A | | RIBONUCLEASE H2 SUBUNIT A (PTHR10954:SF7) | endoribonuclease(PC00094) | DNA replication(GO:0006260);DNA biosynthetic process(GO:0071897);mismatch repair(GO:0006298);RNA catabolic process(GO:0006401) | DNA replication(GO:0006260);mismatch repair(GO:0006298);RNA catabolic process(GO:0006401);DNA replication, removal of RNA primer(GO:0043137);RNA phosphodiester bond hydrolysis, endonucleolytic(GO:0090502) |
| *Rrm2b* | *RRM2B* | Q7LG56 | Ribonucleoside-diphosphate reductase subunit M2 B | GWAS candidate | RIBONUCLEOSIDE-DIPHOSPHATE REDUCTASE SUBUNIT M2 B (PTHR23409:SF19) | reductase(PC00198) | nucleotide biosynthetic process(GO:0009165);carbohydrate derivative biosynthetic process(GO:1901137) | DNA synthesis involved in DNA repair(GO:0000731);kidney development(GO:0001822);renal system process(GO:0003014);mitochondrial DNA replication(GO:0006264);DNA repair(GO:0006281);response to oxidative stress(GO:0006979);ribonucleoside diphosphate metabolic process(GO:0009185);deoxyribonucleoside triphosphate metabolic process(GO:0009200);deoxyribonucleotide biosynthetic process(GO:0009263);2'-deoxyribonucleotide biosynthetic process(GO:0009265);positive regulation of G2/M transition of mitotic cell cycle(GO:0010971);response to amine(GO:0014075);positive regulation of G0 to G1 transition(GO:0070318);negative regulation of intrinsic apoptotic signaling pathway by p53 class mediator(GO:1902254) |
| *Setd2* | *SETD2* | Q9BYW2 | Histone-lysine N-methyltransferase SETD2 | transcription/regulation by RNA Polymerase II; chromatin organization/remodeling | HISTONE-LYSINE N-METHYLTRANSFERASE SETD2 (PTHR46711:SF1) | histone modifying enzyme(PC00261) | | angiogenesis(GO:0001525);morphogenesis of a branching structure(GO:0001763);neural tube closure(GO:0001843);mismatch repair(GO:0006298);chromatin remodeling(GO:0006338);regulation of DNA-templated transcription(GO:0006355);transcription elongation by RNA polymerase II(GO:0006368);regulation of gene expression(GO:0010468);positive regulation of autophagy(GO:0010508);regulation of double-strand break repair via homologous recombination(GO:0010569);regulation of mRNA export from nucleus(GO:0010793);peptidyl-lysine trimethylation(GO:0018023);forebrain [illegible] |
| *Setdb1* | *SETDB1* | Q15047 | Histone-lysine N-methyltransferase SETDB1 | chromatin organization/remodeling | HISTONE-LYSINE N-METHYLTRANSFERASE SETDB1 (PTHR46024:SF2) | histone modifying enzyme(PC00261) | histone lysine methylation(GO:0034968);regulation of chromosome organization(GO:0033044);heterochromatin assembly(GO:0031507);positive regulation of cellular component biogenesis(GO:0044089);positive regulation of organelle organization(GO:0010638);regulation of chromatin assembly(GO:0010847);gene expression(GO:0010467);DNA metabolic process(GO:0006259) | [illegible] |
| *Setx* | *SETX* | Q7Z333 | Probable helicase senataxin | transcription/regulation by RNA Polymerase II | HELICASE SENATAXIN-RELATED (PTHR10887:SF495) | RNA helicase(PC00032) | | [illegible] |
| *Sirt6* | *SIRT6* | Q8N6T7 | NAD-dependent protein deacetylase sirtuin-6 | transcription/regulation by RNA Polymerase II | NAD-DEPENDENT PROTEIN DEACETYLASE SIRTUIN-6 (PTHR47651:SF12) | histone modifying enzyme(PC00261) | | [illegible] |
| *Slx4* | *SLX4* | Q8IY92 | Structure-specific endonuclease subunit SLX4 | | STRUCTURE-SPECIFIC ENDONUCLEASE SUBUNIT SLX4 (PTHR21541:SF3) | | nucleic acid phosphodiester bond hydrolysis(GO:0090305);DNA double-strand break processing(GO:0000729);interstrand cross-link repair(GO:0036297);resolution of meiotic recombination intermediates(GO:0000712);meiotic telophase I(GO:0007134) | [illegible] |
| *Smarcad1* | *SMARCAD1* | Q9H4L7 | SWI/SNF-related matrix-associated actin-dependent regulator of chromatin subfamily A containing DEAD/H box 1 | chromatin organization/remodeling | SWI/SNF-RELATED MATRIX-ASSOCIATED ACTIN-DEPENDENT REGULATOR OF CHROMATIN SUBFAMILY A CONTAINING DEAD/H BOX 1 (PTHR10799:SF964) | DNA helicase(PC00011) | | [illegible] |
| *Supt4a* | *SUPT4H1* | P63272 | Transcription elongation factor SPT4 | transcription/regulation by RNA Polymerase II; chromatin organization/remodeling | TRANSCRIPTION ELONGATION FACTOR SPT4 (PTHR12882:SF1) | general transcription factor(PC00259) | regulation of transcription elongation from RNA polymerase II promoter(GO:0034243);mRNA processing(GO:0006397);chromatin organization(GO:0006325);transcription elongation from RNA polymerase II promoter(GO:0006368) | [illegible] |
| *Tcea2* | *TCEA2* | Q15560 | Transcription elongation factor A protein 2 | | TRANSCRIPTION ELONGATION FACTOR A PROTEIN 2 (PTHR11477:SF3) | general transcription factor(PC00259) | transcription, DNA-templated(GO:0006351) | [illegible] |
| *Tcerg1* | *TCERG1* | O14776 | Transcription elongation regulator 1 | GWAS candidate | TRANSCRIPTION ELONGATION REGULATOR 1 (PTHR15377:SF7) | general transcription factor(PC00259) | | [illegible] |
| *Tdp1* | *TDP1* | Q9NUW8 | Tyrosyl-DNA phosphodiesterase 1 | | TYROSYL-DNA PHOSPHODIESTERASE 1 (PTHR12415:SF0) | phosphodiesterase(PC00185) | double-strand break repair(GO:0006302) | [illegible] |
| *Ubr5* | *UBR5* | O95071 | E3 ubiquitin-protein ligase UBR5 | GWAS candidate | E3 UBIQUITIN-PROTEIN LIGASE UBR5 (PTHR46276:SF1) | ubiquitin-protein ligase(PC00234) | regulation of double-strand break repair(GO:2000779);histone H2A ubiquitination(GO:0033522);protein K63-linked ubiquitination(GO:0070534);negative regulation of protein ubiquitination(GO:0031397);regulation of histone modification(GO:0031056);double-strand break repair(GO:0006302) | [illegible] |

| | | | | | | | |
|---|---|---|---|---|---|---|---|
| *Ung* | *UNG* | P13051 | Uracil-DNA glycosylase | URACIL-DNA GLYCOSYLASE (PTHR11264:SF0) | DNA glycosylase(PC00010) | base-excision repair(GO:0006284) | DNA repair(GO:0006281);base-excision repair(GO:0006284);somatic hypermutation of immunoglobulin genes(GO:0016446);negative regulation of apoptotic process(GO:0043066);depyrimidination(GO:0045008);isotype switching(GO:0045190);base-excision repair, AP site formation via deaminated base removal(GO:0097510) |
| *Wrn* | *WRN* | Q14191 | Werner syndrome ATP-dependent helicase | WERNER SYNDROME ATP-DEPENDENT HELICASE (PTHR13710:SF146) | DNA helicase(PC00011) | DNA biosynthetic process(GO:0071897);double-strand break repair via homologous recombination(GO:0000724);DNA unwinding involved in DNA replication(GO:0006268) | telomere maintenance(GO:0000723);double-strand break repair via homologous recombination(GO:0000724);DNA synthesis involved in DNA repair(GO:0000731);DNA metabolic process(GO:0006259);DNA replication(GO:0006260);DNA unwinding involved in DNA replication(GO:0006268);base-excision repair(GO:0006284);double-strand break repair(GO:0006302);DNA damage response(GO:0006974);response to oxidative stress(GO:0006979);brain development(GO:0007420);determination of adult lifespan(GO:0008340);cellular response to starvation(GO:0009267);response to UV-C(GO:0010225);replication fork processing(GO:0031297);telomere maintenance via semi-conservative replication(GO:0032201);DNA duplex unwinding(GO:0032508);regulation of growth rate(GO:0040009);G-quadruplex DNA unwinding(GO:0044806);positive regulation of hydrolase activity(GO:0051345);telomeric D-loop disassembly(GO:0061820);cellular response to gamma radiation(GO:0071480);cellular senescence(GO:0090398);replicative senescence(GO:0090399);t-circle formation(GO:0090656);positive regulation of strand invasion(GO:0098530);protein localization to nucleolus(GO:1902570) |
| *Xpa* | *XPA* | P23025 | DNA repair protein complementing XP-A cells | DNA REPAIR PROTEIN COMPLEMENTING XP-A CELLS (PTHR10142:SF0) | damaged DNA-binding protein(PC00086) | cellular response to light stimulus(GO:0071482);base-excision repair(GO:0006284);nucleotide-excision repair, DNA incision(GO:0033683);chromosome organization(GO:0051276);interstrand cross-link repair(GO:0036297);response to UV(GO:0009411) | nucleotide-excision repair, DNA damage recognition(GO:0000715);DNA repair(GO:0006281);base-excision repair(GO:0006284);nucleotide-excision repair(GO:0006289);UV protection(GO:0009650);protein localization to nucleus(GO:0034504);UV-damage excision repair(GO:0070914);nucleotide-excision repair involved in interstrand cross-link repair(GO:1901255) |

***Supplementary Table 5: In vivo CRISPR editing and CAG instability analysis following PHP.eB treatment***

| Gene | sgRNA ID | Striatum | | | | | | | | |
|---|---|---|---|---|---|---|---|---|---|---|
| | | CAG instability analysis | | | | | CRISPR editing quantification by NGS | | | |
| | | **Index (+ve)** | **ANOVA** | **p-value (vs Empty)** | **Number of animals** | **CAG** | **Frameshift (%)** | **Non-frameshift (%)** | **Unedited (%)** | **Number of animals** |
| *Empty* | | 11.876 ±0.931 | | | 13 (9M, 4F) | 115.7 ±1.9 | | | | |
| *Fan1* | Fan1-JD2.1 | 13.812 ±2.081 | * | 0.0140 | 8 (4M, 4F) | 115.9 ±2.2 | 69.8% ±5.2 | 0.2% ±0.4 | 30.0% ±5.1 | 5 (2M, 3F) |
| *Mlh1* | Mlh1-JD4 | 3.676 ±0.593 | **** | <0.0001 | 10 (6M, 4F) | 116.8 ±1.2 | 66.7% ±12.2 | 0.2% ±0.4 | 33.1% ±12.5 | 3 (2M, 1F) |
| *Mlh3* | Mlh3-JD2 | 3.798 ±0.508 | **** | <0.0001 | 5 (3M, 2F) | 115.8 ±1.3 | 56.7% ±2.9 | 12.3% ±0.7 | 31.0% ±3.5 | 6 (4M, 2F) |
| *Msh3* | Msh3-JD2.3 | 4.672 ±1.277 | **** | <0.0001 | 8 (4M, 4F) | 116.1 ±1.5 | 71.9% ±4.6 | 1.9% ±0.2 | 26.2% ±4.8 | 4 (1M, 3F) |
| *Msh6* | Msh6-JD2.2 | 10.993 ±1.452 | ns | 0.7639 | 6 (3M, 3F) | 115.2 ±1.7 | 45.7% ±6.9 | 22.7% ±10.7 | 31.7% ±6.8 | 7 (3M, 4F) |
| *Pms1* | Pms1-JD2.2 | 7.817 ±0.847 | **** | <0.0001 | 9 (4M, 5F) | 115.2 ±1.4 | 53.6% ±9.5 | 4.2% ±1.4 | 42.2% ±10.8 | 6 (3M, 3F) |
| Pms2 | Pms2-JD2.2 | 15.330 ±2.025 | **** | <0.0001 | 9 (4M, 5F) | 115.6 ±2.0 | 64.9% ±4.2 | 0.5% ±0.2 | 34.6% ±4.4 | 5 (1M, 4F) |
| *Pold1* | Pold1-JD3.1 | 7.049 ±1.720 | **** | <0.0001 | 9 (4M, 5F) | 116.1 ±1.6 | 45.6% ±9.1 | 0.8% ±0.7 | 53.7% ±9.5 | 6 (2M, 4F) |
| *Pold3* | Pold3-JD3.1 | 5.531 ±1.211 | **** | <0.0001 | 9 (5M, 4F) | 115.4 ±2.3 | 70.2% ±0.8 | 0.3% ±0.3 | 29.4% ±1.0 | 3 (2M, 1F) |
| *Untreated* | | 11.431 ±1.065 | ns | 0.9772 | 14 (10M, 4F) | 116.0 ±2.1 | | | | |

| Gene | sgRNA ID | Liver | | | | | | | | |
|---|---|---|---|---|---|---|---|---|---|---|
| | | CAG instability analysis | | | | | CRISPR editing quantification by NGS | | | |
| | | **Index (+ve)** | **ANOVA** | **p-value (vs Empty)** | **Number of animals** | **CAG** | **Frameshift (%)** | **Non-frameshift (%)** | **Unedited (%)** | **Number of animals** |
| *Empty* | | 9.400 ±0.593 | | | 13 (9M, 4F) | 115.7 ±1.9 | | | | |
| *Fan1* | Fan1-JD2.1 | 15.218 ±1.739 | **** | <0.0001 | 8 (4M, 4F) | 115.9 ±2.2 | 64.9% ±3.6 | 1.6% ±1.1 | 33.5% ±4.7 | 5 (2M, 3F) |
| *Mlh1* | Mlh1-JD4 | 2.045 ±0.142 | **** | <0.0001 | 10 (6M, 4F) | 116.8 ±1.2 | 69.0% ±4.6 | 0.2% ±0.3 | 30.8% ±4.5 | 6 (3M, 3F) |
| *Mlh3* | Mlh3-JD2 | 1.989 ±0.197 | **** | <0.0001 | 5 (3M, 2F) | 115.8 ±1.3 | 46.4% ±2.0 | 14.5% ±1.2 | 39.2% ±2.5 | 5 (3M, 2F) |
| *Msh3* | Msh3-JD2.3 | 2.056 ±0.613 | **** | <0.0001 | 8 (4M, 4F) | 116.1 ±1.5 | 66.9% ±3.7 | 2.0% ±0.6 | 31.1% ±4.0 | 6 (3M, 3F) |
| *Msh6* | Msh6-JD2.2 | 9.892 ±0.292 | ns | 0.8566 | 7 (3M, 4F) | 115.4 ±1.7 | 53.9% ±3.8 | 8.9% ±2.7 | 37.2% ±3.4 | 8 (4M, 4F) |
| *Pms1* | Pms1-JD2.2 | 3.546 ±0.741 | **** | <0.0001 | 9 (4M, 5F) | 115.2 ±1.4 | 56.6% ±2.5 | 7.6% ±2.4 | 35.8% ±4.2 | 6 (3M, 3F) |
| Pms2 | Pms2-JD2.2 | 11.696 ±0.791 | **** | <0.0001 | 9 (4M, 5F) | 115.3 ±2.1 | 65.2% ±2.4 | 2.0% ±0.8 | 32.8% ±2.6 | 7 (4M, 3F) |
| *Pold1* | Pold1-JD3.1 | 5.954 ±1.158 | **** | <0.0001 | 9 (4M, 5F) | 116.1 ±1.6 | 24.1% ±6.7 | 2.0% ±1.6 | 74.0% ±7.4 | 6 (3M, 3F) |
| *Pold3* | Pold3-JD3.1 | 5.115 ±1.517 | **** | <0.0001 | 9 (5M, 4F) | 115.4 ±2.3 | 36.4% ±8.7 | 3.2% ±2.6 | 60.4% ±10.2 | 6 (3M, 3F) |
| *Untreated* | | 9.408 ±0.271 | ns | >0.9999 | 14 (10M, 4F) | 116.0 ±2.1 | | | | |

***Supplementary Table 6: CAG instability analysis following single vs dual AAV8 treatment***

| Gene | Age (wks) | CAG instability analysis | | | *p*-values from one-way ANOVA (Dunnett's multiple comparison correction) | | | | | | | | | | | | | | | | | | | | | | | | | | | | |
|---|---|---|---|---|---|---|---|---|---|---|---|---|---|---|---|---|---|---|---|---|---|---|---|---|---|---|---|---|---|---|---|---|---|
| | | Liver Index (+ve) | Number of animals | CAG | Untreated (6wks) | Untreated (12wks) | Empty | *LacZ* | *Msh2* | *Msh3* | *Mlh1* | *Mlh3* | *Pms1* | *Msh6* | *Pms2* | *Fan1* | *Msh3 + Mlh3* | *Msh3 + Pms1* | *Mlh3 + Pms1* | *Msh3 + Msh6* | *Mlh3 + Msh6* | *Pms1 + Msh6* | *Msh3 + Pms2* | *Mlh3 + Pms2* | *Pms1 + Pms2* | *Msh2 + Fan1* | *Msh3 + Fan1* | *Mlh1 + Fan1* | *Mlh3 + Fan1* | *Pms1 + Fan1* | *Msh6 + Pms2* | *Msh6 + Fan1* | *Pms2 + Fan1* |
| Untreated | 6 | 1.775 ±0.265 | 28 (23M, 5F) | 119.2 ±3.5 | | <0.0001 | <0.0001 | <0.0001 | >0.9999 | >0.9999 | 0.1265 | >0.9999 | 0.606 | <0.0001 | <0.0001 | <0.0001 | >0.9999 | >0.9999 | >0.9999 | >0.9999 | >0.9999 | 0.9627 | >0.9999 | 0.9956 | 0.4047 | >0.9999 | >0.9999 | 0.8613 | >0.9999 | 0.9986 | <0.0001 | <0.0001 | <0.0001 |
| Untreated | 12 | 3.799 ±0.319 | 77 (56M, 21F) | 116.0 ±2.1 | <0.0001 | | <0.0001 | 0.0002 | <0.0001 | <0.0001 | <0.0001 | <0.0001 | <0.0001 | <0.0001 | <0.0001 | <0.0001 | <0.0001 | <0.0001 | <0.0001 | <0.0001 | <0.0001 | <0.0001 | <0.0001 | <0.0001 | <0.0001 | <0.0001 | <0.0001 | <0.0001 | <0.0001 | <0.0001 | 0.4689 | <0.0001 | <0.0001 |
| Empty | 12 | 4.159 ±0.294 | 30 (21M, 9F) | 115.3 ±2.4 | <0.0001 | <0.0001 | | >0.9999 | <0.0001 | <0.0001 | <0.0001 | <0.0001 | <0.0001 | <0.0001 | <0.0001 | <0.0001 | <0.0001 | <0.0001 | <0.0001 | <0.0001 | <0.0001 | <0.0001 | <0.0001 | <0.0001 | <0.0001 | <0.0001 | <0.0001 | <0.0001 | <0.0001 | <0.0001 | >0.9999 | <0.0001 | <0.0001 |
| *LacZ* | 12 | 4.293 ±0.294 | 12 (5M, 7F) | 115.6 ±2.4 | <0.0001 | 0.0002 | >0.9999 | | <0.0001 | <0.0001 | <0.0001 | <0.0001 | <0.0001 | 0.0077 | <0.0001 | <0.0001 | <0.0001 | <0.0001 | <0.0001 | <0.0001 | <0.0001 | <0.0001 | <0.0001 | <0.0001 | <0.0001 | <0.0001 | <0.0001 | <0.0001 | <0.0001 | <0.0001 | >0.9999 | <0.0001 | <0.0001 |
| *Msh2* | 12 | 1.857 ±0.159 | 7 (7M) | 115.9 ±2.0 | >0.9999 | <0.0001 | <0.0001 | <0.0001 | | >0.9999 | 0.9718 | >0.9999 | 0.9967 | <0.0001 | <0.0001 | <0.0001 | >0.9999 | >0.9999 | >0.9999 | >0.9999 | >0.9999 | >0.9999 | >0.9999 | >0.9999 | 0.911 | >0.9999 | >0.9999 | 0.9992 | >0.9999 | >0.9999 | <0.0001 | <0.0001 | <0.0001 |
| *Msh3* | 12 | 1.797 ±0.183 | 12 (9M, 3F) | 117.0 ±1.7 | >0.9999 | <0.0001 | <0.0001 | <0.0001 | >0.9999 | | 0.5456 | >0.9999 | 0.8828 | <0.0001 | <0.0001 | <0.0001 | >0.9999 | >0.9999 | >0.9999 | >0.9999 | >0.9999 | 0.992 | >0.9999 | 0.9995 | 0.6295 | >0.9999 | >0.9999 | 0.9656 | >0.9999 | >0.9999 | <0.0001 | <0.0001 | <0.0001 |
| *Mlh1* | 12 | 2.160 ±0.141 | 11 (8M, 3F) | 115.6 ±1.9 | 0.1265 | <0.0001 | <0.0001 | <0.0001 | 0.9718 | 0.5456 | | 0.9793 | >0.9999 | <0.0001 | <0.0001 | <0.0001 | 0.9945 | 0.9945 | 0.9417 | 0.7614 | 0.8989 | >0.9999 | 0.2486 | >0.9999 | >0.9999 | 0.9865 | 0.9815 | >0.9999 | >0.9999 | >0.9999 | <0.0001 | <0.0001 | <0.0001 |
| *Mlh3* | 12 | 1.864 ±0.223 | 7 (7M) | 115.4 ±2.2 | >0.9999 | <0.0001 | <0.0001 | <0.0001 | >0.9999 | >0.9999 | 0.9793 | | 0.9978 | <0.0001 | <0.0001 | <0.0001 | >0.9999 | >0.9999 | >0.9999 | >0.9999 | >0.9999 | >0.9999 | >0.9999 | >0.9999 | 0.9239 | >0.9999 | >0.9999 | 0.9995 | >0.9999 | >0.9999 | <0.0001 | <0.0001 | <0.0001 |
| *Pms1* | 12 | 2.155 ±0.074 | 6 (6M) | 116.3 ±2.3 | 0.606 | <0.0001 | <0.0001 | <0.0001 | 0.9967 | 0.8828 | >0.9999 | 0.9978 | | <0.0001 | <0.0001 | <0.0001 | 0.9991 | 0.9988 | 0.9804 | 0.9455 | 0.9751 | >0.9999 | 0.5433 | >0.9999 | >0.9999 | 0.9965 | 0.9949 | >0.9999 | >0.9999 | >0.9999 | <0.0001 | <0.0001 | <0.0001 |
| *Msh6* | 12 | 4.805 ±0.413 | 16 (16M) | 117.2 ±1.9 | <0.0001 | <0.0001 | <0.0001 | 0.0077 | <0.0001 | <0.0001 | <0.0001 | <0.0001 | <0.0001 | | 0.0019 | <0.0001 | <0.0001 | <0.0001 | <0.0001 | <0.0001 | <0.0001 | <0.0001 | <0.0001 | <0.0001 | <0.0001 | <0.0001 | <0.0001 | <0.0001 | <0.0001 | <0.0001 | 0.0632 | <0.0001 | <0.0001 |
| *Pms2* | 12 | 5.358 ±0.590 | 12 (10M, 2F) | 117.1 ±1.9 | <0.0001 | <0.0001 | <0.0001 | <0.0001 | <0.0001 | <0.0001 | <0.0001 | <0.0001 | <0.0001 | 0.0019 | | <0.0001 | <0.0001 | <0.0001 | <0.0001 | <0.0001 | <0.0001 | <0.0001 | <0.0001 | <0.0001 | <0.0001 | <0.0001 | <0.0001 | <0.0001 | <0.0001 | <0.0001 | <0.0001 | 0.0654 | <0.0001 |
| *Fan1* | 12 | 7.442 ±0.530 | 16 (14M, 2F) | 115.7 ±1.4 | <0.0001 | <0.0001 | <0.0001 | <0.0001 | <0.0001 | <0.0001 | <0.0001 | <0.0001 | <0.0001 | <0.0001 | <0.0001 | | <0.0001 | <0.0001 | <0.0001 | <0.0001 | <0.0001 | <0.0001 | <0.0001 | <0.0001 | <0.0001 | <0.0001 | <0.0001 | <0.0001 | <0.0001 | <0.0001 | <0.0001 | <0.0001 | 0.8219 |
| *Msh3+Mlh3* | 12 | 1.836 ±0.063 | 4 (2M, 2F) | 117.5 ±2.4 | >0.9999 | <0.0001 | <0.0001 | <0.0001 | >0.9999 | >0.9999 | 0.9945 | >0.9999 | 0.9991 | <0.0001 | <0.0001 | <0.0001 | | >0.9999 | >0.9999 | >0.9999 | >0.9999 | >0.9999 | >0.9999 | >0.9999 | 0.952 | >0.9999 | >0.9999 | 0.9997 | >0.9999 | >0.9999 | <0.0001 | <0.0001 | <0.0001 |
| *Msh3+Pms1* | 12 | 1.799 ±0.135 | 3 (3M) | 118.7 ±0.6 | >0.9999 | <0.0001 | <0.0001 | <0.0001 | >0.9999 | >0.9999 | 0.9945 | >0.9999 | 0.9988 | <0.0001 | <0.0001 | <0.0001 | >0.9999 | | >0.9999 | >0.9999 | >0.9999 | 0.9998 | >0.9999 | >0.9999 | 0.9491 | >0.9999 | >0.9999 | 0.9995 | >0.9999 | >0.9999 | <0.0001 | <0.0001 | <0.0001 |
| *Mlh3+Pms1* | 12 | 1.725 ±0.102 | 3 (3M) | 117.3 ±2.1 | >0.9999 | <0.0001 | <0.0001 | <0.0001 | >0.9999 | >0.9999 | 0.9417 | >0.9999 | 0.9804 | <0.0001 | <0.0001 | <0.0001 | >0.9999 | >0.9999 | | >0.9999 | >0.9999 | 0.9965 | >0.9999 | 0.9997 | 0.8263 | >0.9999 | >0.9999 | 0.9906 | >0.9999 | >0.9999 | <0.0001 | <0.0001 | <0.0001 |
| *Msh3+Msh6* | 12 | 1.797 ±0.188 | 8 (8M) | 117.8 ±2.4 | >0.9999 | <0.0001 | <0.0001 | <0.0001 | >0.9999 | >0.9999 | 0.7614 | >0.9999 | 0.9455 | <0.0001 | <0.0001 | <0.0001 | >0.9999 | >0.9999 | >0.9999 | | >0.9999 | 0.996 | >0.9999 | 0.9998 | 0.7238 | >0.9999 | >0.9999 | 0.9833 | >0.9999 | >0.9999 | <0.0001 | <0.0001 | <0.0001 |
| *Mlh3+Msh6* | 12 | 1.780 ±0.086 | 5 (5M) | 117.6 ±2.1 | >0.9999 | <0.0001 | <0.0001 | <0.0001 | >0.9999 | >0.9999 | 0.8989 | >0.9999 | 0.9751 | <0.0001 | <0.0001 | <0.0001 | >0.9999 | >0.9999 | >0.9999 | >0.9999 | | 0.9975 | >0.9999 | 0.9998 | 0.7996 | >0.9999 | >0.9999 | 0.9909 | >0.9999 | >0.9999 | <0.0001 | <0.0001 | <0.0001 |
| *Pms1+Msh6* | 12 | 2.165 ±0.013 | 3 (3M) | 117.0 ±2.6 | 0.9627 | <0.0001 | <0.0001 | <0.0001 | >0.9999 | 0.992 | >0.9999 | >0.9999 | >0.9999 | <0.0001 | <0.0001 | <0.0001 | >0.9999 | 0.9998 | 0.9965 | 0.996 | 0.9975 | | 0.8715 | >0.9999 | >0.9999 | 0.9995 | 0.9993 | >0.9999 | >0.9999 | >0.9999 | <0.0001 | <0.0001 | <0.0001 |
| *Msh3+Pms2* | 12 | 1.653 ±0.132 | 6 (6M) | 116.8 ±1.7 | >0.9999 | <0.0001 | <0.0001 | <0.0001 | >0.9999 | >0.9999 | 0.2486 | >0.9999 | 0.5433 | <0.0001 | <0.0001 | <0.0001 | >0.9999 | >0.9999 | >0.9999 | >0.9999 | >0.9999 | 0.8715 | | 0.9626 | 0.3103 | >0.9999 | >0.9999 | 0.7346 | >0.9999 | 0.9803 | <0.0001 | <0.0001 | <0.0001 |
| *Mlh3+Pms2* | 12 | 2.107 ±0.205 | 3 (3M) | 118.0 ±0.0 | 0.9956 | <0.0001 | <0.0001 | <0.0001 | >0.9999 | 0.9995 | >0.9999 | >0.9999 | >0.9999 | <0.0001 | <0.0001 | <0.0001 | >0.9999 | >0.9999 | 0.9997 | 0.9998 | 0.9998 | >0.9999 | 0.9626 | | >0.9999 | >0.9999 | >0.9999 | >0.9999 | >0.9999 | >0.9999 | <0.0001 | <0.0001 | <0.0001 |
| *Pms1+Pms2* | 12 | 2.337 ±0.154 | 3 (3M) | 118.3 ±0.6 | 0.4047 | <0.0001 | <0.0001 | <0.0001 | 0.911 | 0.6295 | >0.9999 | 0.9239 | >0.9999 | <0.0001 | <0.0001 | <0.0001 | 0.952 | 0.9491 | 0.8263 | 0.7238 | 0.7996 | >0.9999 | 0.3103 | >0.9999 | | 0.9193 | 0.9051 | >0.9999 | 0.9967 | >0.9999 | <0.0001 | <0.0001 | <0.0001 |
| *Msh2+Fan1* | 12 | 1.775 ±0.178 | 3 (3M) | 117.3 ±1.5 | >0.9999 | <0.0001 | <0.0001 | <0.0001 | >0.9999 | >0.9999 | 0.9865 | >0.9999 | 0.9965 | <0.0001 | <0.0001 | <0.0001 | >0.9999 | >0.9999 | >0.9999 | >0.9999 | >0.9999 | 0.9995 | >0.9999 | >0.9999 | 0.9193 | | >0.9999 | 0.9984 | >0.9999 | >0.9999 | <0.0001 | <0.0001 | <0.0001 |
| *Msh3+Fan1* | 12 | 1.765 ±0.217 | 3 (3M) | 118.0 ±1.0 | >0.9999 | <0.0001 | <0.0001 | <0.0001 | >0.9999 | >0.9999 | 0.9815 | >0.9999 | 0.9949 | <0.0001 | <0.0001 | <0.0001 | >0.9999 | >0.9999 | >0.9999 | >0.9999 | >0.9999 | 0.9993 | >0.9999 | >0.9999 | 0.9051 | >0.9999 | | 0.9977 | >0.9999 | >0.9999 | <0.0001 | <0.0001 | <0.0001 |
| *Mlh1+Fan1* | 12 | 2.165 ±0.239 | 4 (4M) | 118.0 ±1.2 | 0.8613 | <0.0001 | <0.0001 | <0.0001 | 0.9992 | 0.9656 | >0.9999 | 0.9995 | >0.9999 | <0.0001 | <0.0001 | <0.0001 | 0.9997 | 0.9995 | 0.9906 | 0.9833 | 0.9909 | >0.9999 | 0.7346 | >0.9999 | >0.9999 | 0.9984 | 0.9977 | | >0.9999 | >0.9999 | <0.0001 | <0.0001 | <0.0001 |
| *Mlh3+Fan1* | 12 | 1.899 ±0.275 | 3 (3M) | 118.0 ±1.0 | >0.9999 | <0.0001 | <0.0001 | <0.0001 | >0.9999 | >0.9999 | >0.9999 | >0.9999 | >0.9999 | <0.0001 | <0.0001 | <0.0001 | >0.9999 | >0.9999 | >0.9999 | >0.9999 | >0.9999 | >0.9999 | >0.9999 | >0.9999 | 0.9967 | >0.9999 | >0.9999 | >0.9999 | | >0.9999 | <0.0001 | <0.0001 | <0.0001 |
| *Pms1+Fan1* | 12 | 2.084 ±0.202 | 3 (3M) | 116.3 ±2.5 | 0.9986 | <0.0001 | <0.0001 | <0.0001 | >0.9999 | >0.9999 | >0.9999 | >0.9999 | >0.9999 | <0.0001 | <0.0001 | <0.0001 | >0.9999 | >0.9999 | >0.9999 | >0.9999 | >0.9999 | >0.9999 | 0.9803 | >0.9999 | >0.9999 | >0.9999 | >0.9999 | >0.9999 | >0.9999 | | <0.0001 | <0.0001 | <0.0001 |
| *Msh6+Pms2* | 12 | 4.213 ±0.268 | 5 (5M) | 117.4 ±2.3 | <0.0001 | 0.4689 | >0.9999 | >0.9999 | <0.0001 | <0.0001 | <0.0001 | <0.0001 | <0.0001 | 0.0632 | <0.0001 | <0.0001 | <0.0001 | <0.0001 | <0.0001 | <0.0001 | <0.0001 | <0.0001 | <0.0001 | <0.0001 | <0.0001 | <0.0001 | <0.0001 | <0.0001 | <0.0001 | <0.0001 | | <0.0001 | <0.0001 |
| *Msh6+Fan1* | 12 | 6.101 ±0.176 | 3 (3M) | 117.7 ±2.3 | <0.0001 | <0.0001 | <0.0001 | <0.0001 | <0.0001 | <0.0001 | <0.0001 | <0.0001 | <0.0001 | <0.0001 | 0.0654 | <0.0001 | <0.0001 | <0.0001 | <0.0001 | <0.0001 | <0.0001 | <0.0001 | <0.0001 | <0.0001 | <0.0001 | <0.0001 | <0.0001 | <0.0001 | <0.0001 | <0.0001 | <0.0001 | | <0.0001 |
| *Pms2+Fan1* | 12 | 7.915 ±0.186 | 3 (3M) | 118.0 ±1.0 | <0.0001 | <0.0001 | <0.0001 | <0.0001 | <0.0001 | <0.0001 | <0.0001 | <0.0001 | <0.0001 | <0.0001 | <0.0001 | 0.8219 | <0.0001 | <0.0001 | <0.0001 | <0.0001 | <0.0001 | <0.0001 | <0.0001 | <0.0001 | <0.0001 | <0.0001 | <0.0001 | <0.0001 | <0.0001 | <0.0001 | <0.0001 | <0.0001 | |
